# A chromosome-scale super-pangenome of the lichen genus *Peltigera* reveals genome architecture and expanded interaction repertoires shared across pathogenic and mutualistic fungi

**DOI:** 10.64898/2026.06.16.732702

**Authors:** Vivien Joisten-Rosenthal, Tolga Arslan, Max Heinen, Ciaran Kelly, Yukiyo Sato, Diego Garfias-Gallegos, Lucas Hüttebräucker, Stefan Robertz, Vicente Ramírez, Jochen Hecht, Francy J Pérez-Llanos, Carlos Pardo De la Hoz, Jolanta Miadlikowska, Xiaoran Zhou, Dirk Joisten, Jan P Buchmann, Max Schmidt, Jasmin Almer, Reynir Freyr Reynisson, Silke Werth, Philip Nakonz, Michael Feldbrügge, Markus Pauly, François Lutzoni, Bart P.H.J. Thomma, Florian Altegoer, Björn Usadel

**Author notes:** Heinrich-Heine University Düsseldorf, Faculty of Mathematics and Natural Sciences, Institute of Quantitative and Theoretical Biology, Düsseldorf, Germany (Stefan Robertz). Institute of Plant Science and Resources, Okayama University, Kurashiki, Okayama 710-0046, Japan (Yukiyo Sato). HS Geisenheim University, Department of Plant Breeding, Geisenheim, Germany. Corresponding authors: Vivien Joisten-Rosenthal, Björn Usadel.

## Abstract

Fungi engage in associations with other organisms across a continuum from pathogenic to mutualistic lifestyles. Hence, they require a compendium of molecular capacities, including partner recognition, extracellular signaling, nutrient exchange, immune modulation, and control of microbial competitors. In filamentous pathogens such traits are frequently associated with compartmentalized genomes, including rapidly evolving secreted proteins known as effectors and expanded receptor families, but it remains unclear whether similar genomic principles shape mutualistic fungal symbioses. Here, we generated a chromosome-scale super-pangenome for the lichen-forming genus *Peltigera*, comprising 41 mycobiont assemblies representing eleven species, together with genomes of associated *Nostoc* and, in tripartite species, *Coccomyxa* photobionts. The mycobiont genomes revealed extensive variation in genome size, transposable element content, biosynthetic gene clusters, and lineage-specific gene content, with pronounced expansions in tripartite species. Across *Peltigera*, secreted protein encoding genes were preferentially located in TE-rich regions. We further identified *Starship*-like transposon elements, expanded antimicrobial protein repertoires, and a large, previously underestimated repertoire of fungal GPCRs dominated by Pth11-like receptors. Layer-specific transcriptomics of a *P. rufescens* thallus showed differential expression of several interaction-associated gene families, e.g. lectins, antimicrobial proteins and Pth11-like GPCRs. These data indicate that pathogenic and mutualistic fungi might exhibit shared genomic principles, including genome compartmentalization, mobile-element-associated diversification, and the expansion of molecular repertoires involved in recognition, extracellular control and signaling.

## Introduction

Lichens and their symbioses represent stable, long-term associations that require continuous coordination between interacting partners. In lichens, a filamentous fungus (termed mycobiont) undergoes tightly coordinated interactions with one or few photosynthetic partners (termed photobionts), mostly unicellular green algae, cyanobacteria, or both (Lutzoni and Miadlikowska, 2009; Spribille et al., 2022; Scharnagl et al., 2023). The evolutionary success of lichens is reflected by their remarkably widespread occurrence and diversity. Lichens colonize a wide range of habitats, from tropical forests to extreme environments such as arctic tundra, deserts, and alpine ecosystems, where they often dominate primary productivity and biomass, but they are also found in aquatic ecosystems, both freshwater and marine habitats (Lutzoni and Miadlikowska, 2009; Asplund and Wardle, 2017; Muggia and Grube, 2018). They play key roles in ecosystem functioning, including carbon and nitrogen cycling, soil formation, and contribute to habitat structuring by providing resources and microenvironments for diverse microbial and invertebrate communities.

Within the lichen, photosynthetically derived carbon, typically in the form of glucose exported from cyanobacteria or polyols exported from algal photobionts, are transferred to the mycobiont and converted into fungal storage compounds (Spribille et al., 2022). Recent genomic and transcriptomic studies have identified candidate transport systems, including sugar- and polyol transporters as well as amino acid permeases, that are differentially regulated during symbiosis and likely mediate bidirectional nutrient fluxes (Yoshino et al., 2019; Resl et al., 2022). In cyanolichens, comparative studies of *Nostoc* symbionts have revealed transcriptional and genomic signatures consistent with nitrogen provisioning, including reduced nitrogen assimilation capacity and loss of a high-affinity ammonium permease, suggesting a metabolic state that favors transfer of fixed nitrogen to the mycobiont (Garfias-Gallegos et al., 2025).

Beyond resource exchange, the molecular mechanisms that govern lichen establishment and development remain largely unresolved (Joneson et al., 2011). However, lectins have long been proposed as mediators of partner recognition and maintenance due to their ability to bind defined carbohydrate motifs. In *Peltigera membranacea*, expression of the galectin-like proteins LEC-1 and LEC-2 is influenced by the presence of the photobiont, supporting a role for lectins in symbiotic interactions (Manoharan et al., 2012; Miao et al., 2012). Similarly, in *P. canina* the mycobiont secretes an arginase with lectin-like activity that interacts with a polygalactosylated urease from compatible photobionts preventing cellular uptake of the arginase, suggesting highly specific carbohydrate-mediated recognition processes (Díaz et al., 2009; Nazem-Bokaee et al., 2021). In incompatible photobionts the arginase is taken up negatively influencing N-metabolism resulting in cell wall disruption and thus death.

In addition to symbiont recognition and resource exchange, stable lichen symbiosis requires tight coordination of growth and differentiation between symbiotic partners, yet the signaling processes underlying this coordination are only beginning to be explored. Potential insights come from work on *Umbilicaria muhlenbergii*, where G-protein signaling and cyclic AMP (cAMP) pathways have been proposed to coregulate dimorphism and symbiosis (Wang et al., 2020). While dimorphic fungi such as *U. muhlenbergii* seem to be the exception among the lichen forming fungi, comparative genomic analyses suggest that components of these signaling cascades are differentially regulated in other lichenized fungi such as *Cladonia grayi* (Armaleo et al., 2019) and *Xanthoria parietina (Tagirdzhanova et al., 2025b)*. However, while these studies establish that canonical fungal signaling pathways are involved in lichen development, the upstream signals and respective receptors that coordinate interactions between symbiotic partners remain unknown. In parallel, recent work has begun to uncover additional molecular layers of interaction that might be mediated by effector-like proteins which have been identified in lichen-forming fungi such as *X. parietina*, suggesting that the mycobiont may actively modulate photobiont physiology and potentially other members of the lichen-associated microbiome (Tagirdzhanova et al., 2025b).

Studies of filamentous plant pathogens have shown that genes involved in host interaction are often embedded in dynamic, repeat-rich genome compartments, where gene family expansion, rapid divergence, and structural variation can drive lifestyle adaptation (Dong et al., 2015; Fouché et al., 2018). Whether similar genomic principles shape mutualistic fungal symbioses remains largely unresolved, in part because this question requires assemblies that preserve long-range genomic context. This is especially relevant for effector-like secreted proteins, small secreted peptides, antimicrobial proteins, and expanded receptor families, which can be lineage-specific, highly divergent, or associated with repetitive regions, making them difficult to annotate and compare in fragmented genomes. However with some recent exceptions (Adams et al., 2023; Cameron et al., 2026; Dong et al., 2026), the majority of lichen genome assemblies are highly fragmented, thus lacking sufficient depth to characterize gene abundance and link it to lifestyle adaptations.

To overcome these limitations, we generated genome assemblies for 41 samples with most of them being chromosome scale genomes covering eleven *Peltigera* species representing five of the eight recognized sections (Miadlikowska and Lutzoni, 2000) and combined these single genomes into a gene based comparative framework (i.e. super-pangenome). In our case, this allowed us to distinguish genomic features shared across *Peltigera* from those specific to individual species or lineages, including variation in gene content, genome architecture, and repeat-rich regions. This genomic resource enables the systematic identification of interaction-associated repertoires, including secreted potential antimicrobial and effector-like proteins as well as signaling receptor families. In parallel, the improved contiguity allows us to resolve transposable element landscapes and assess their contribution to genome architecture and gene diversification. By integrating gene inventories with their genomic context, our approach provides the resolution required to link interaction-related repertoires with the evolutionary forces shaping them.

## Results

### *Peltigera* sampling and genome sequencing

To obtain a comprehensive genomic overview of the genus *Peltigera*, we sampled *Peltigera* lichen across Central and Western Europe and included three samples obtained previously (Resl et al., 2022), (Supplementary Table S1). In total, we obtained 41 samples representing nine bipartite cyanolichen species (containing *Nostoc* as their photobiont): *Peltigera didactyla*, *P. canina, P. monticola, P. neorufescens, P. ponojensis, P. praetextata, and P. rufescens* (section *Peltigera*), *P. hymenina* (section Polydactylon), *P. neckeri* (section *Horizontales*), as well as two tripartite lichen species *P. aphthosa* (section *Peltidea*) and *P. leucophlebia* (section *Chloropeltigera*) which harbour both a cyanobacterial (*Nostoc*) and a green algal (*Coccomyxa*) photobiont (Table 1). *P. hymenina, P. leucophlebia, P. neorufescens* and *P. ponojensis* were each represented by a single sample only, whereas we obtained multiples samples for the other seven species, including eight samples for *P. rufescens* and ten for *P. monticola* (Table 1).

**Table 1:**
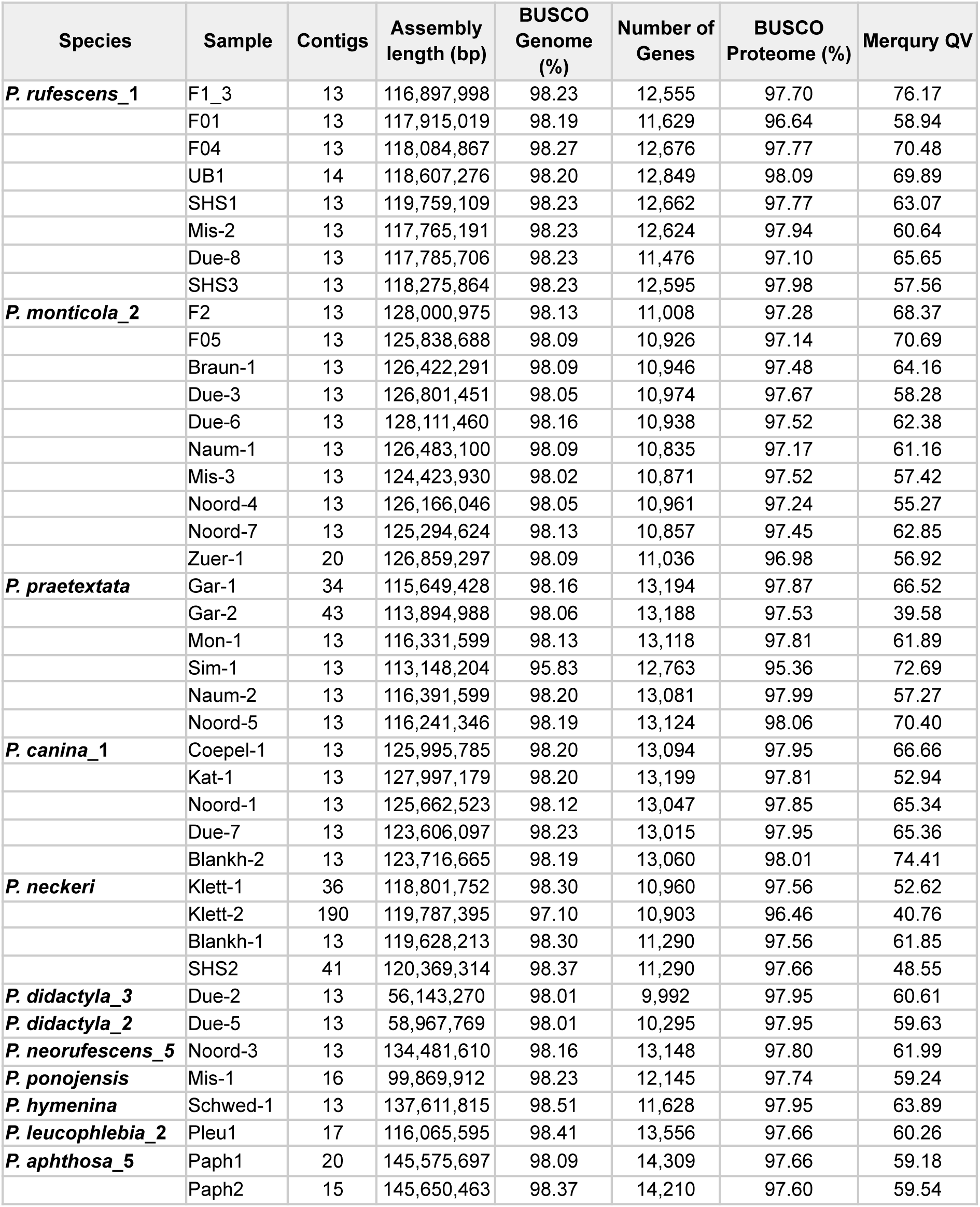
Assembly statistics, BUSCO completeness and Merqury QV for all *Peltigera* mycobiont genomes.

| Species | Sample | Contigs | Assembly length (bp) | BUSCO Genome (%) | Number of Genes | BUSCO Proteome (%) | Merqury QV |
| --- | --- | --- | --- | --- | --- | --- | --- |
| *P. rufescens_1* | F1_3 | 13 | 116,897,998 | 98.23 | 12,555 | 97.70 | 76.17 |
|  | F01 | 13 | 117,915,019 | 98.19 | 11,629 | 96.64 | 58.94 |
|  | F04 | 13 | 118,084,867 | 98.27 | 12,676 | 97.77 | 70.48 |
|  | UB1 | 14 | 118,607,276 | 98.20 | 12,849 | 98.09 | 69.89 |
|  | SHS1 | 13 | 119,759,109 | 98.23 | 12,662 | 97.77 | 63.07 |
|  | Mis-2 | 13 | 117,765,191 | 98.23 | 12,624 | 97.94 | 60.64 |
|  | Due-8 | 13 | 117,785,706 | 98.23 | 11,476 | 97.10 | 65.65 |
|  | SHS3 | 13 | 118,275,864 | 98.23 | 12,595 | 97.98 | 57.56 |
| *P. monticola_2* | F2 | 13 | 128,000,975 | 98.13 | 11,008 | 97.28 | 68.37 |
|  | F05 | 13 | 125,838,688 | 98.09 | 10,926 | 97.14 | 70.69 |
|  | Braun-1 | 13 | 126,422,291 | 98.09 | 10,946 | 97.48 | 64.16 |
|  | Due-3 | 13 | 126,801,451 | 98.05 | 10,974 | 97.67 | 58.28 |
|  | Due-6 | 13 | 128,111,460 | 98.16 | 10,938 | 97.52 | 62.38 |
|  | Naum-1 | 13 | 126,483,100 | 98.09 | 10,835 | 97.17 | 61.16 |
|  | Mis-3 | 13 | 124,423,930 | 98.02 | 10,871 | 97.52 | 57.42 |
|  | Noord-4 | 13 | 126,166,046 | 98.05 | 10,961 | 97.24 | 55.27 |
|  | Noord-7 | 13 | 125,294,624 | 98.13 | 10,857 | 97.45 | 62.85 |
|  | Zuer-1 | 20 | 126,859,297 | 98.09 | 11,036 | 96.98 | 56.92 |
| *P. praetextata* | Gar-1 | 34 | 115,649,428 | 98.16 | 13,194 | 97.87 | 66.52 |
|  | Gar-2 | 43 | 113,894,988 | 98.06 | 13,188 | 97.53 | 39.58 |
|  | Mon-1 | 13 | 116,331,599 | 98.13 | 13,118 | 97.81 | 61.89 |
|  | Sim-1 | 13 | 113,148,204 | 95.83 | 12,763 | 95.36 | 72.69 |
|  | Naum-2 | 13 | 116,391,599 | 98.20 | 13,081 | 97.99 | 57.27 |
|  | Noord-5 | 13 | 116,241,346 | 98.19 | 13,124 | 98.06 | 70.40 |
| *P. canina_1* | Coepel-1 | 13 | 125,995,785 | 98.20 | 13,094 | 97.95 | 66.66 |
|  | Kat-1 | 13 | 127,997,179 | 98.20 | 13,199 | 97.81 | 52.94 |
|  | Noord-1 | 13 | 125,662,523 | 98.12 | 13,047 | 97.85 | 65.34 |
|  | Due-7 | 13 | 123,606,097 | 98.23 | 13,015 | 97.95 | 65.36 |
|  | Blankh-2 | 13 | 123,716,665 | 98.19 | 13,060 | 98.01 | 74.41 |
| *P. neckeri* | Klett-1 | 36 | 118,801,752 | 98.30 | 10,960 | 97.56 | 52.62 |
|  | Klett-2 | 190 | 119,787,395 | 97.10 | 10,903 | 96.46 | 40.76 |
|  | Blankh-1 | 13 | 119,628,213 | 98.30 | 11,290 | 97.56 | 61.85 |
|  | SHS2 | 41 | 120,369,314 | 98.37 | 11,290 | 97.66 | 48.55 |
| *P. didactyla_3* | Due-2 | 13 | 56,143,270 | 98.01 | 9,992 | 97.95 | 60.61 |
| *P. didactyla_2* | Due-5 | 13 | 58,967,769 | 98.01 | 10,295 | 97.95 | 59.63 |
| *P. neorufescens_5* | Noord-3 | 13 | 134,481,610 | 98.16 | 13,148 | 97.80 | 61.99 |
| *P. ponojensis* | Mis-1 | 16 | 99,869,912 | 98.23 | 12,145 | 97.74 | 59.24 |
| *P. hymenina* | Schwed-1 | 13 | 137,611,815 | 98.51 | 11,628 | 97.95 | 63.89 |
| *P. leucophlebia_2* | Pleu1 | 17 | 116,065,595 | 98.41 | 13,556 | 97.66 | 60.26 |
| *P. aphthosa_5* | Paph1 | 20 | 145,575,697 | 98.09 | 14,309 | 97.66 | 59.18 |
|  | Paph2 | 15 | 145,650,463 | 98.37 | 14,210 | 97.60 | 59.54 |

We then subjected these samples to long-read nanopore and HiFi sequencing, generating between 6.9 and 235.0 Gbases of Nanopore data (with quality score of 10) and between 8.3 and 73.2 Gbases of HiFi data per sample, respectively (Supplementary Table S2).

### Mycobiont genome assembly and annotation

Mycobiont and photobiont genomes were assembled using a hybrid approach combining Oxford Nanopore and PacBio HiFi data and subsequently corrected using Inspector (Chen et al., 2021b). Non-mycobiont or photobiont contigs were removed using fcs-gx (Astashyn et al., 2024). After extracting the photobiont genomes (i.e. *Nostoc spp.* and *Coccomyxa spp.)*, the final mycobiont genomes were assessed using EukCC (Saary et al., 2020) which indicated completeness values ranging from 96.99% for *P. praetextata Sim-1* to 100% for multiple samples of *P. canina, P. praetextata* and *P. monticola*. For each species at least one assembly reached at least 99.25% completeness (Supplementary Table S3).

For several species, the initial assemblies already resolved the genome into 13 contigs likely representing complete chromosomes. We generated Pore-C chromosome conformation capture data for nine samples representing six species whose genomes remained more fragmented: *P. rufescens* (F1_3, F01, F04), *P. monticola* (F2, F05), *P. praetextata* (Sim-1), *P. canina* (Coepel-1), *P. neorufescens* (Noord-3), and *P. neckeri* (Blankh-1). This combined approach yielded assemblies consisting of 13 scaffolds or contigs for eight bipartite *Peltigera* species (Table 1). We were able to detect telomeric sequences on multiple chromosomes and five assemblies representing four species achieved telomere-to-telomere (T2T) completeness across all 13 chromosomes (Figure 1). In total, 30 of the 41 assemblies resolved the genome into 13 chromosomes (Table 1). For the two tripartite species, *P. leucophlebia* yielded 17 contigs and *P. aphthosa* yielded 15 and 20 contigs for its two isolates.

**Figure 1:**
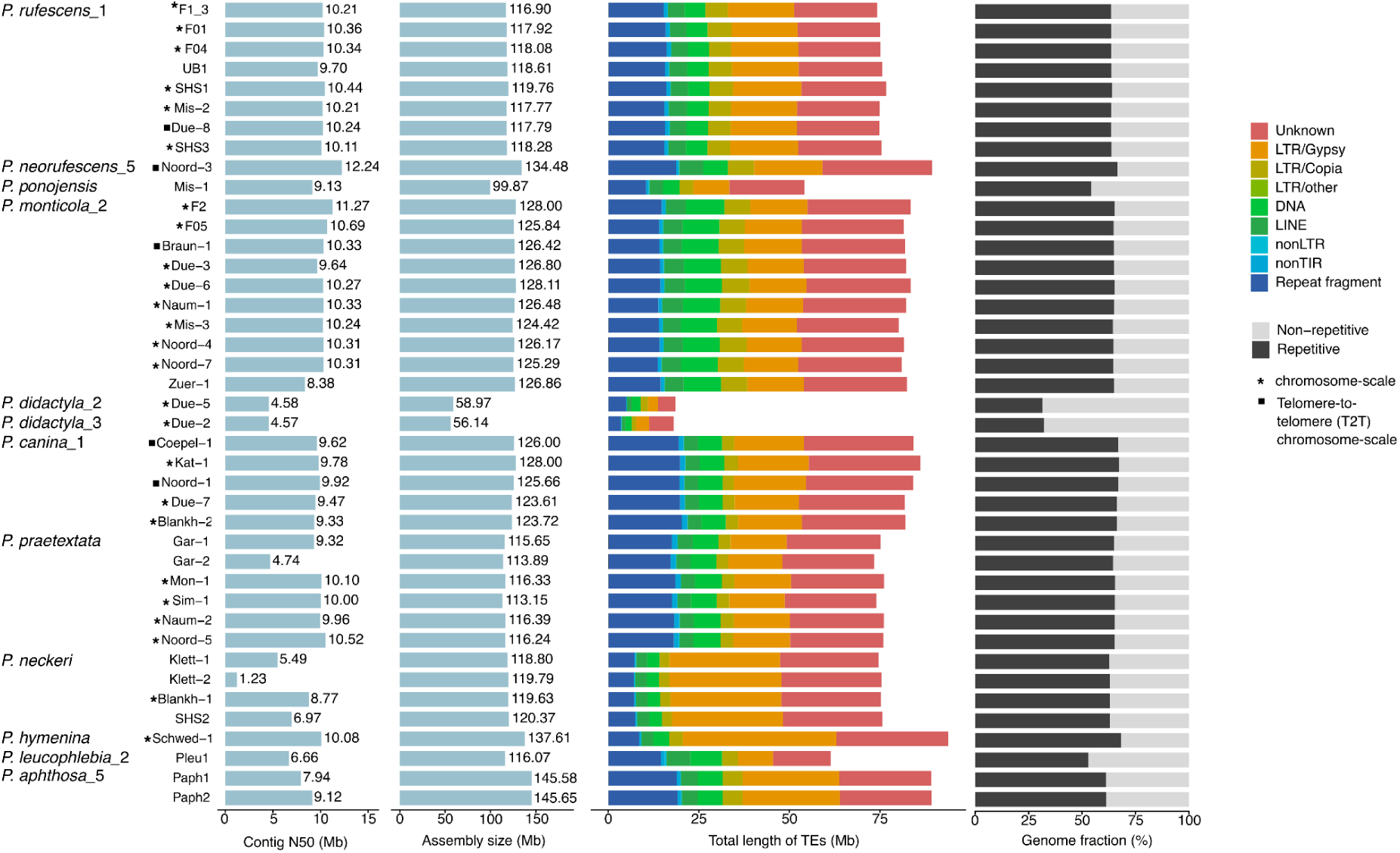
Genome size correlates with repeat content across *Peltigera* mycobiont genomes. Bar plots show contig N50 values of the genome assemblies and total assembly size. The third panel illustrates the genomic occupancy of transposable elements and repeats, expressed as the total length of TE- and repeat-derived sequences across each genome. The fourth panel shows the genomic fractions occupied by repetitive and non-repetitive sequences. Asterisks (*) indicate chromosome-scale assemblies, and square symbols indicate that all chromosomes assembled telomere-to-telomere.

The mycobiont assemblies exhibited genome sizes centered around 120 Mb, but showed substantial interspecific variation. The smallest genomes were those of *P. didactyla*, at approximately 56 and 59 Mb for its two lineages i.e. roughly half the size of other *Peltigera* species. At the other extreme, *P. aphthosa* reached approximately 145.5 Mb, closely followed by *P. hymenina* at 137.6 Mb. Species-level N50 values ranged from 4.6 Mb for *P. didactyla* to 12.2 Mb for *P. neorufescens*, with an overall average N50 of 9.1 Mb and N90 of 5.9 Mb across all 41 assemblies (Figure 1, Supplementary Table S4). Assembly quality was assessed using multiple complementary metrics. Merqury (Rhie et al., 2020) per-base accuracy scores (QV) averaged above 60 across all samples, with values ranging from 39.58 (*P. praetextata* Gar-1) to 76.17 (*P. rufescens* F1_3); for each species, at least one assembly exceeded QV 59 (Table 1). Genome-level BUSCO (Simão et al., 2015) completeness against the ascomycota dataset ranged from 95.83% (*P. praetextata* Sim-1) to 98.51% (*P. hymenina* Schwed-1), with every species reaching at least 98% in its best assembly. CRAQ (Li et al., 2023) regional quality scores (R-AQI) exceeded 90 for all assemblies except *P. neckeri* Klett-2 (84.22), and structural quality scores (S-AQI) ranged from 91.44 (Klett-2) to 100 (Supplementary Table S5). These metrics confirmed that the assemblies are of reference genome quality and likely suitable for resolving repeat-rich and gene-sparse regions that are typically fragmented in short-read lichen genome assemblies.

Gene annotation was performed by combining predictions from BRAKER3 (Gabriel et al., 2024), Helixer (Holst et al., 2025) and StringTie (Kovaka et al., 2019) using Mikado (Venturini et al., 2018) as an integrator. Across the 41 genomes, annotated gene counts ranged from 9,992 to 14,309. Consistent with their smaller genome size, *P. didactyla* isolates encoded 9,992 and 10,295 genes, whereas the tripartite species *P. leucophlebia* (13,556 genes) and *P. aphthosa* (14,210 and 14,309 genes) harboured the highest number of genes. Annotated proteome BUSCO completeness values ranged from 95.36% (*P. praetextata* Sim-1) to 98.09% (*P. rufescens* UB1), with every species reaching at least 97.66% in its best assembly close to the genome BUSCO values (Supplementary Table S4).

Functional annotations were generated by integrating eggNOG, InterProScan, SwissProt/BLAST, KEGG/KOfamScan and UniProt mappings against curated fungal references including *Saccharomyces cerevisiae* and *Aspergillus nidulans*. This yielded 9631 InterPro (Supplementary Table S6) and 25882 putative KEGG distinct terms. After filtering (Interproscan E-value of <10⁻^10^ ; KEGG terms < specific HMM score thresholds (Aramaki et al., 2020)) reduced these unique term numbers to 8371 and 3417, respectively (Supplementary Table S7). Thus, functional annotations were assigned to between 81.5% (Paph1, *P. aphthosa*) and 86.7% (*P. neckeri* Klett-1) of predicted proteins, substantially reducing the proportion of hypothetical proteins across all assemblies.

### Photobiont genome assemblies

Next, we recovered the genomes of associated photobionts from the metagenomic assemblies. For the cyanobacterial partner *Nostoc spp.*, single-contig assemblies were obtained for 40 of the 41 samples, with genome sizes ranging from 6.79 Mb (*P. aphthosa* Paph1) to 8.35 Mb (*P. aphthosa* Paph2) and a mean genome size of 7.71 Mb (Supplementary Table S8) in line with typical genome sizes (Gagunashvili and Andrésson, 2018; Pardo-De la Hoz et al., 2025). BUSCO completeness against the nostocales_odb12 dataset exceeded 99.5% for all single-contig assemblies, and the annotated proteome completeness ranged from 98.6% to 99.6%, confirming near-complete gene space recovery. The *Nostoc* genomes encoded between 5,670 and 7,085 protein-coding genes similar to previous *Nostoc* analyses (Gagunashvili and Andrésson, 2018; Huo et al., 2021). The sole exception was *P. rufescens* isolate Mis-2, whose *Nostoc* genome remained fragmented into 23 contigs with a substantially reduced BUSCO completeness of 77.8%, likely reflecting lower sequencing coverage of the photobiont in this single sample. To determine the phylogenetic placement of the newly sequenced *Nostoc* cyanobionts, we extracted the *rbcLX* region from each genome and classified the sequences according to the framework of (Pardo-De la Hoz et al., 2025). The majority of cyanobionts (n = 31) were assigned to *Nostoc* Section 3.1 and were associated with multiple mycobiont species. In contrast, all cyanobionts associated with *Peltigera praetextata* as well as the cyanobiont associated with *P. didactyla_3* belonged to Section 3.6. The cyanobiont associated with *P. leucophlebia_2* represented the only member of Section 3.4 in our dataset. Notably, the two cyanobionts associated with *P. aphthosa_5* were assigned to distinct phylogenomic sections. Paph1 clustered within Section 3.10 together with the cyanobiont of *P. hymenina* (Schwed-1), whereas Paph2 was assigned to Section 2.4. The placement of these cyanobionts in distinct phylogenomic sections may explain the observed differences in genome size (Supplementary Table S9).

For the two tripartite species *P. aphthosa_*5 and *P. leucophlebia_*2, algal *Coccomyxa spp* genomes were additionally recovered. To determine their likely taxonomic identity, ITS sequences were extracted assigning all algal photobionts to *Coccomyxa solorinae*. Near-complete assemblies were obtained for both *P. aphthosa* isolates, with genome sizes of 48.3 Mb (Paph1, 19 contigs, BUSCO 99.4%) and 47.9 Mb (Paph2, 23 contigs, BUSCO 98.2%). Analysis of telomeric repeat distributions, combined with comparisons to the chromosome-scale genome of *Coccomyxa elongata* (Kraege et al., 2025), which contains 19 complete chromosomes, indicated that the algal photobiont genome of Paph1 was recovered as a complete telomere-to-telomere (T2T) chromosome-scale assembly. The 19 contigs of Paph1 showed a high correspondence with the 19 nuclear scaffolds of *C. elongata* and were flanked by telomeric repeat motifs (CCCTAAA/TTTAGGG) at both ends. The Paph2 assembly also showed strong correspondence to the 19 *C. elongata* chromosomes. However, its 23 contigs indicate that several chromosomes were split across multiple contigs, resulting in a highly complete but not fully contiguous assembly. In contrast, the *Coccomyxa* assembly from *P. leucophlebia* (Pleu1) was more fragmented (95 contigs, 41.8 Mb) with a BUSCO completeness of 85.1%, indicating that a fraction of the algal genome was not captured in the assembly (Supplementary Table S10).

### Phylogenetic and Synteny analysis

To place the 41 newly generated mycobiont genomes within a phylogenomic context, we added these genomes to 100 other *Peltigera* genomes. Three genomes representing *Solorina* and *Pseudosolorina* were included to root the tree, for a total of 144 Peltigeraceae genomes (Supplementary Table S9). The tree was inferred using weighted ASTRAL based on 2,826 genes (Figure 2). Species identifications were largely consistent with phylogenomic clustering. All *Peltigera rufescens* samples formed a well-supported clade corresponding to *P. rufescens*_1 (Magain et al., 2018). The sample Noord-3, identified as *P. neorufescens*_5 formed a distinct clade together with the publicly available sample SRR14561385 (*Peltigera neorufescens*). This analysis also revealed that in total only one *P. ponojensis* thallus (Mis-1) was sampled. All *Peltigera monticola* samples formed one monophyletic group identified as *P. monticola*_2. The two specimens of *P. didactyla* represent two different lineages, provisionally named *P. didactyla*_2 and *P. didactyla*_3. All *Peltigera canina* samples were nested within a well-supported *P. canina*_1 clade. The *Peltigera leucophlebia* sample and the two *P. aphthosa* samples represent *P. leucophlebia_*2 and *P. aphthosa*_5, respectively (Pardo-De la Hoz et al., 2025). This phylogenomic analysis shows that the 41 samples comprised 33 specimens from section *Peltigera*, four from section *Horizontales*, two from section *Peltidea* and one each from sections *Polydactylon* and *Chloropeltigera,* representing eleven species.

**Figure 2:**
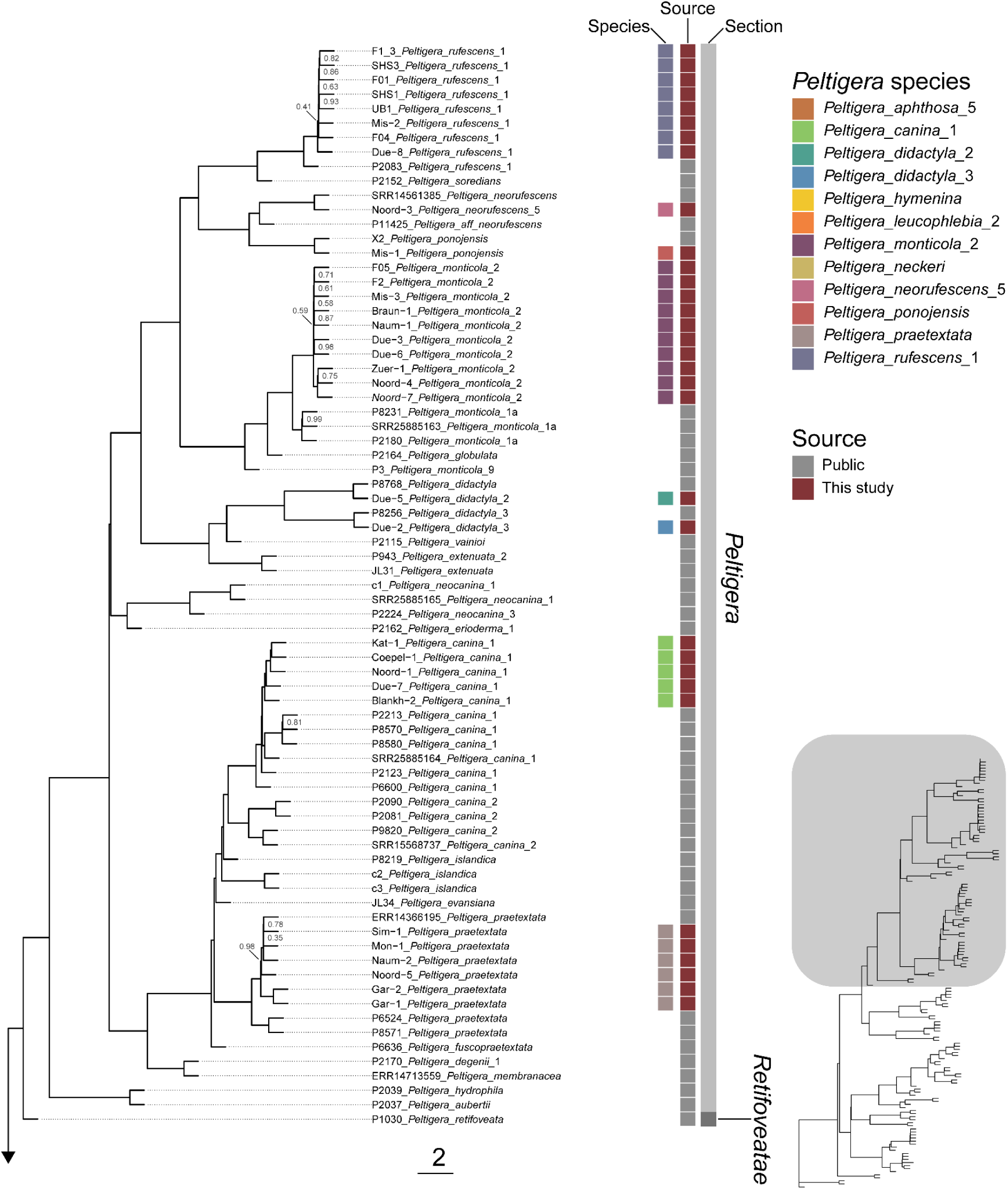
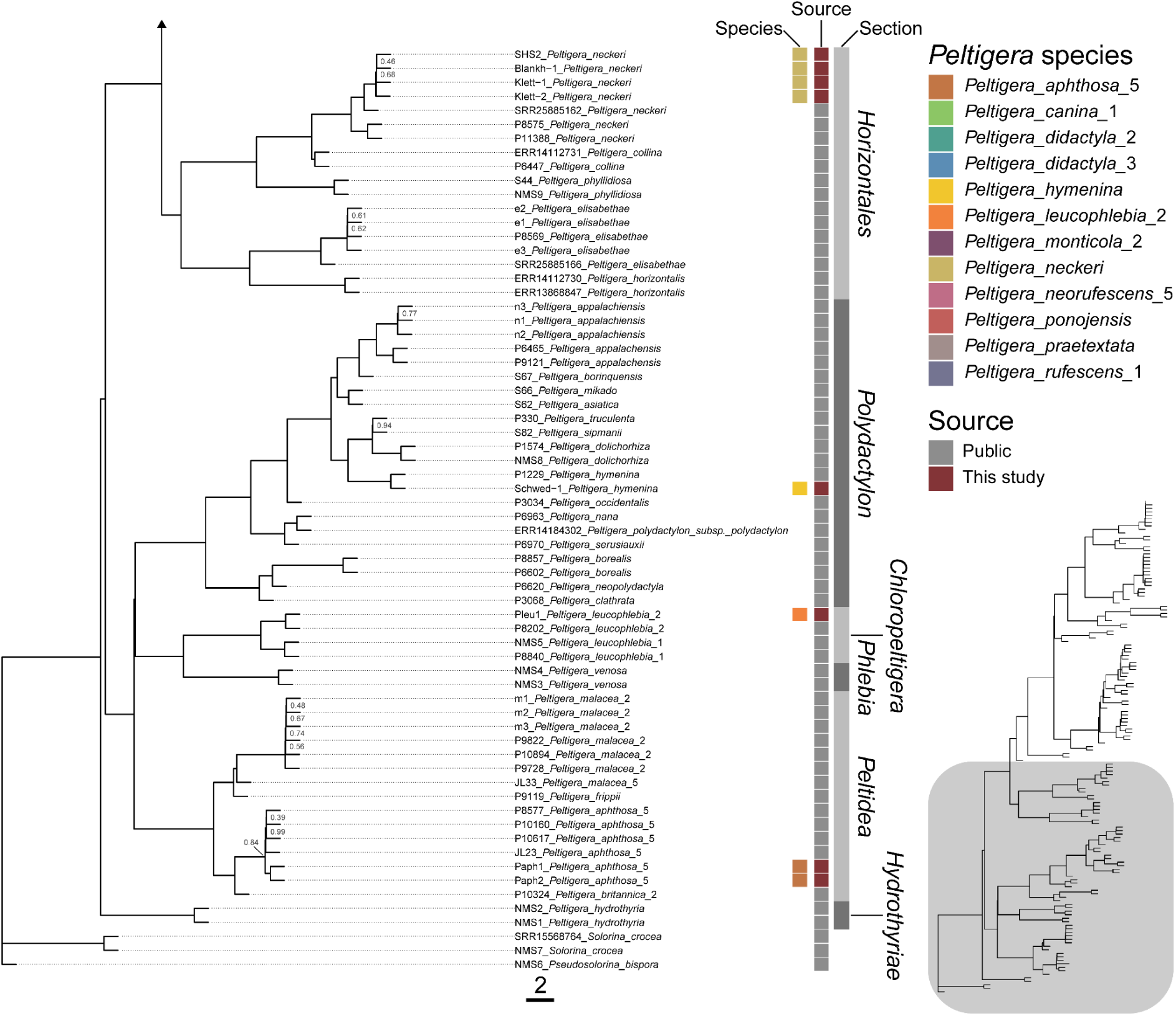
Phylogenomic tree of the genus *Peltigera*. Species tree based on 144 *Peltigeraceae* genomes, including three outgroup taxa representing *Solorina* and *Pseudosolorina* (Supplementary Table S9). The tree was inferred with weighted ASTRAL using 2,826 genes. Branch lengths and scale represent coalescent units. Numbers next to nodes are Local Posterior Probabilities (LPP); only LPPs lower than 1.00 are shown. Genome metadata is in Supplementary Table S11. The first column to the right of the tree indicates the species of the genomes generated by this study. The Source column indicates whether a genome was sequenced for this study or retrieved from public databases. The last column indicates the taxonomic section of *Peltigera* species. Numbers after species name indicate lineage designation based on previous phylogenetic studies (Magain et al., 2018; Miadlikowska et al., 2018; Pardo-De la Hoz et al., 2018).

To assess structural conservation and divergence across these genomes, we performed gene-based synteny analyses. Intraspecific comparisons across the nine chromosome-scale *P. monticola_*2 genomes revealed a high degree of gene collinearity across all 13 chromosomes (Tang et al., 2024) (Figure 3A, Supplementary Figure 1). Nucleotide-level synteny analysis (Coombe et al., 2025) broadly confirmed this conservation but revealed additional local rearrangements not apparent from gene order alone (Supplementary Figure 2). Interspecific comparisons using one representative per species showed broad conservation of syntenic blocks across chromosomes, but also pronounced structural rearrangements, especially involving more distantly related taxa such as *P. hymenina* from section *Polydactylon* (Figure 3B). For example, regions corresponding to chromosome 3 in *P. monticola_2* aligned with chromosome 4 in *P. didactyla_3*, indicating inter-chromosomal translocations. The extent of structural divergence scaled with phylogenetic distance, with closely related species retaining largely collinear chromosomes while more distant lineages showed progressively greater reorganization (Figure 3B).

**Figure 3:**
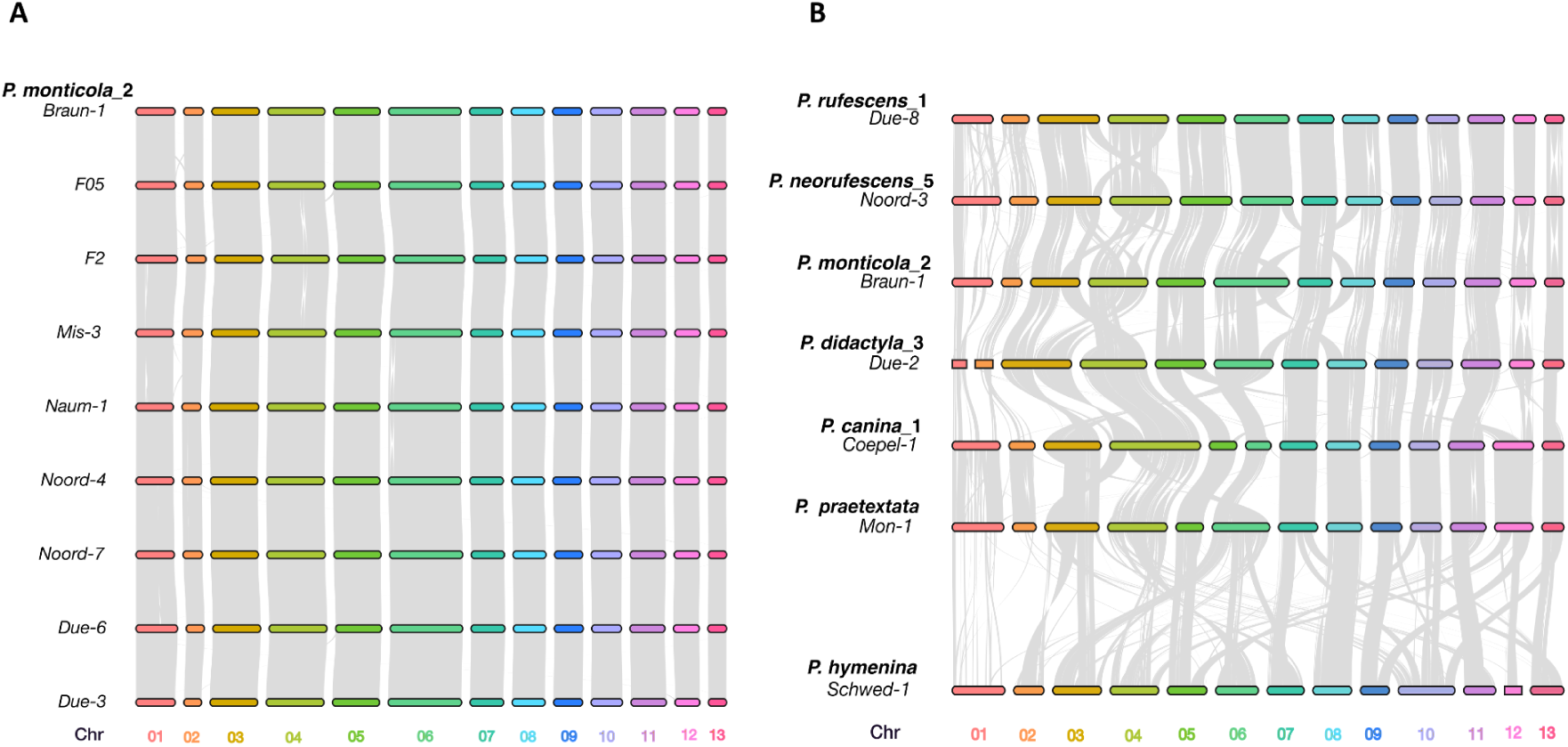
Intra- and inter-species synteny comparisons across mycobiont chromosomes. Gene-based synteny relationships are visualized as grey ribbons connecting homologous genomic regions between chromosomes. Intra-species synteny is shown for *Peltigera monticola_*2 (A), highlighting conserved gene order within the genome. Inter-species synteny comparisons include all *Peltigera* species for which chromosome-scale genome assemblies are available, illustrating the conservation and rearrangement of chromosomal regions across species (B). Each ribbon represents a gene-based syntenic link between corresponding loci.

### Transposable element landscape

Having established the genome assemblies and their phylogenomic relationships, we next investigated the repeat and transposable element (TE) content that underlies the observed variation in genome size. Using PanEDTA (Ou et al., 2024), we found substantial variation in repeat content among the *Peltigera* mycobiont genomes (Figure 1). Genome size was positively correlated with TE abundance across species (r=0.75, Spearman rho=0.377, p=0.02). The most striking case was *P. didactyla*, whose markedly smaller genomes (56 and 59 Mb for the two clades) also harboured the lowest proportion of repetitive elements (Figure 1). At the other extreme, the large genomes of *P. hymenina* (137.6 Mb) and the tripartite lichen *P. aphthosa* (145.5 Mb) were correspondingly enriched in TEs, suggesting that genome expansion in these lineages has been driven by TE accumulation.

LTR/Ty3/Gypsy elements represented the most abundant TE class in all genomes, but their relative abundance varied considerably among species. *P. hymenina* showed the highest Gypsy content, where these elements comprise nearly 30% of the genome, followed by *P. neckeri* at approximately 25%. All other species ranged from 4.75% to 18.39% (Supplementary Table S12). In addition, the tripartite lichen *P. leucophlebia_2* and *P. aphthosa_5* uniquely harboured piggyBac TIR elements. Although these occupied only approximately 0.02% of the genome, they were entirely absent from all bipartite species analyzed (Supplementary Table S12).

#### “Two-speed genome” traits

In filamentous fungal plant pathogens, secreted protein genes and particularly effectors are often located in gene-sparse, TE-rich genomic regions, giving rise to a “two-speed genome” architecture in which conserved core regions coexist with rapidly evolving plastic compartments (Raffaele and Kamoun, 2012; Seidl and Thomma, 2017; Cook et al., 2020). This organization has recently also been reported in a beneficial plant fungal symbiont (Oliveira et al., 2025), suggesting it may be a more general feature of fungi that maintain intimate interactions with other organisms. To test whether lichen-forming fungi display a similar architecture, we compared the distance from secreted protein genes to their nearest flanking TEs against that of other protein-coding genes across all 41 *Peltigera* genomes. Indeed, TEs were significantly closer to secreted protein genes than to non-secreted genes in all 41 genomes (Figure 4). Interestingly, when secreted protein genes were further categorized into genes that encode antimicrobial proteins (Mesny et al., 2026), those that encode carbohydrate-active enzymes and remaining secreted proteins that comprise putative effector genes, each of the three categories showed equally close association to TEs (Supplementary Figure 3).

**Figure 4.**
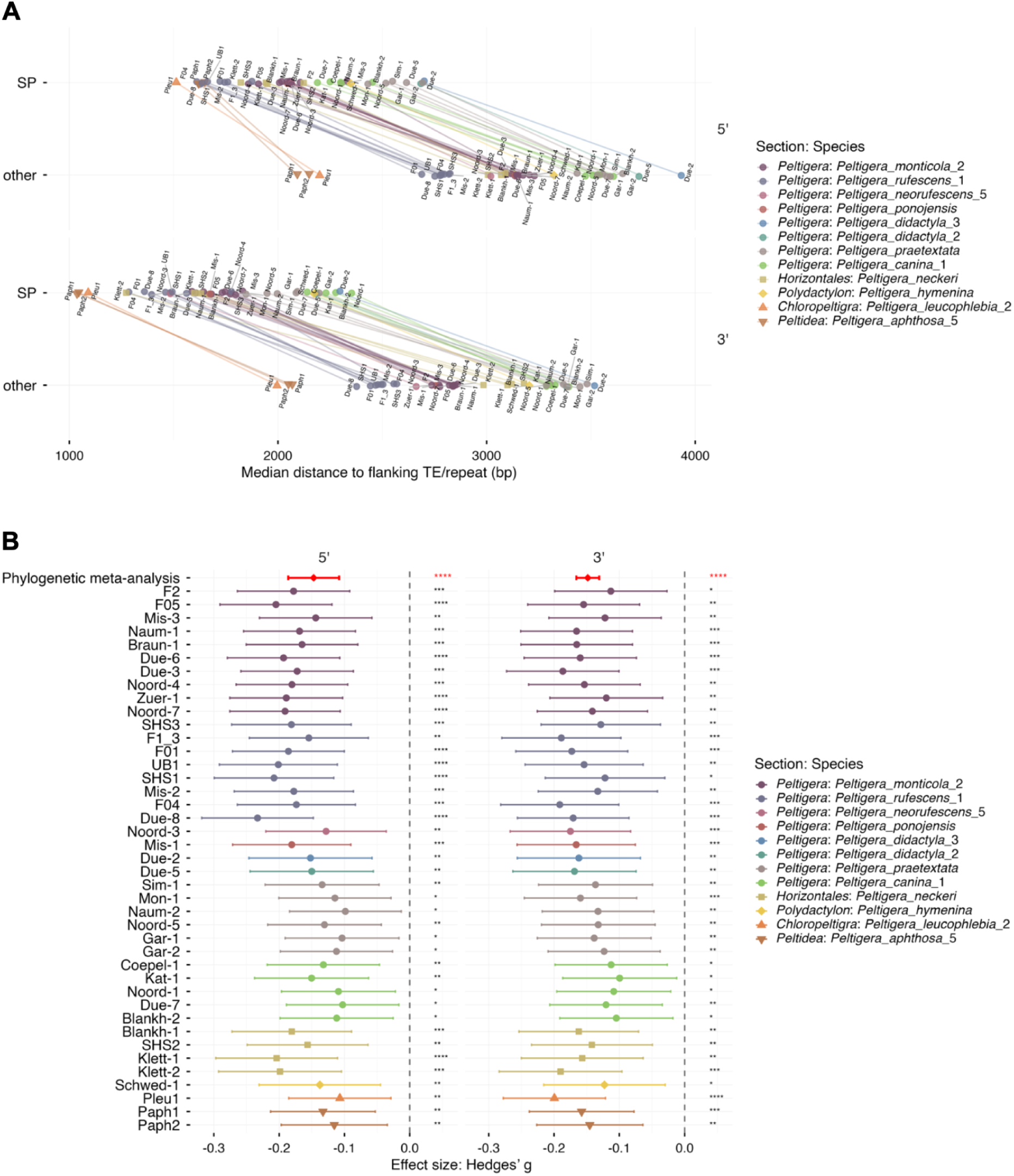
Secreted protein (SP) genes are located closer to transposable elements (TEs) than other protein-coding genes across the genus *Peltigera*. (A) Plot of the median distance to flanking TE/repeats for SP genes and other protein-encoding genes, shown separately for each sample and for the 5’ or 3’ side. Points represent median values, and lines connect the median values of SP genes and other protein-encoding genes within each *Peltigera* sample. (B) Forest plots showing the standardized mean differences (Hedges’ g) in genomic distances to the nearest TEs between SP and other protein-coding genes for each sample and the flanking direction (5’ and 3’). Each point and horizontal line represent the estimated effect size and its 95% confidence interval for individual samples. Red diamonds and bold red error bars indicate the summary effect sizes and 95% confidence intervals derived from a phylogenetic multi-level meta-analysis, which accounts for both phylogenetic relatedness and species-specific random effects. Statistical significance was determined using false discovery rate-corrected *p*-values (*: *P*<0.05, **: *P*<0.01, ***: *P*<0.001, ****: *P*<0.0001). Negative effect sizes indicate that the distance to the nearest TE is smaller in SP genes compared to other protein-coding genes.

Accordingly, secreted protein genes were enriched in gene-sparse regions (Supplementary Figure 4), which is statistically significant for all the 41 *Peltigera* genomes (Supplementary Figure 5). Together, these patterns are indicative of a typical “two-speed genome” organization that is similarly observed in various filamentous fungal pathogens.

#### *Starship*-like element annotation

Large mobile elements termed *Starships* have recently emerged as important drivers of genome plasticity in fungi, mobilizing dozens to hundreds of cargo genes via the activity of a tyrosine recombinase (YR) “captain” gene at their 5’ end (Bucknell and McDonald, 2023; Urquhart et al., 2023; Gluck-Thaler and Vogan, 2024; Urquhart et al., 2024). In the pathogenic fungus *Verticillium dahliae*, *Starships* have been shown to constitute a major component of the dynamic genomic compartment (Sato et al., 2025), and a *Starship*-like element termed *Tangerine* has been described in the lichen-forming genus *Xanthoria (Tagirdzhanova et al., 2025a)*. Given the TE-rich, compartmentalized architecture of *Peltigera* genomes, we investigated whether *Starship*-associated elements are also present in this genus.

We screened all 41 assemblies for captain-like YR genes and identified between one and 14 per genome, belonging to diverse navis and family classifications (Figure 5A). YR genes were ubiquitous across the genus, but their repertoires differed markedly between species. The Prometheus and Enterprise families were broadly distributed across most lineages, whereas Phoenix, Tardis, and Galactica YR genes showed a more restricted distribution. In contrast, the Arwing family YR gene was detected exclusively in *P. hymenina* (Figure 5A).

**Figure 5.**
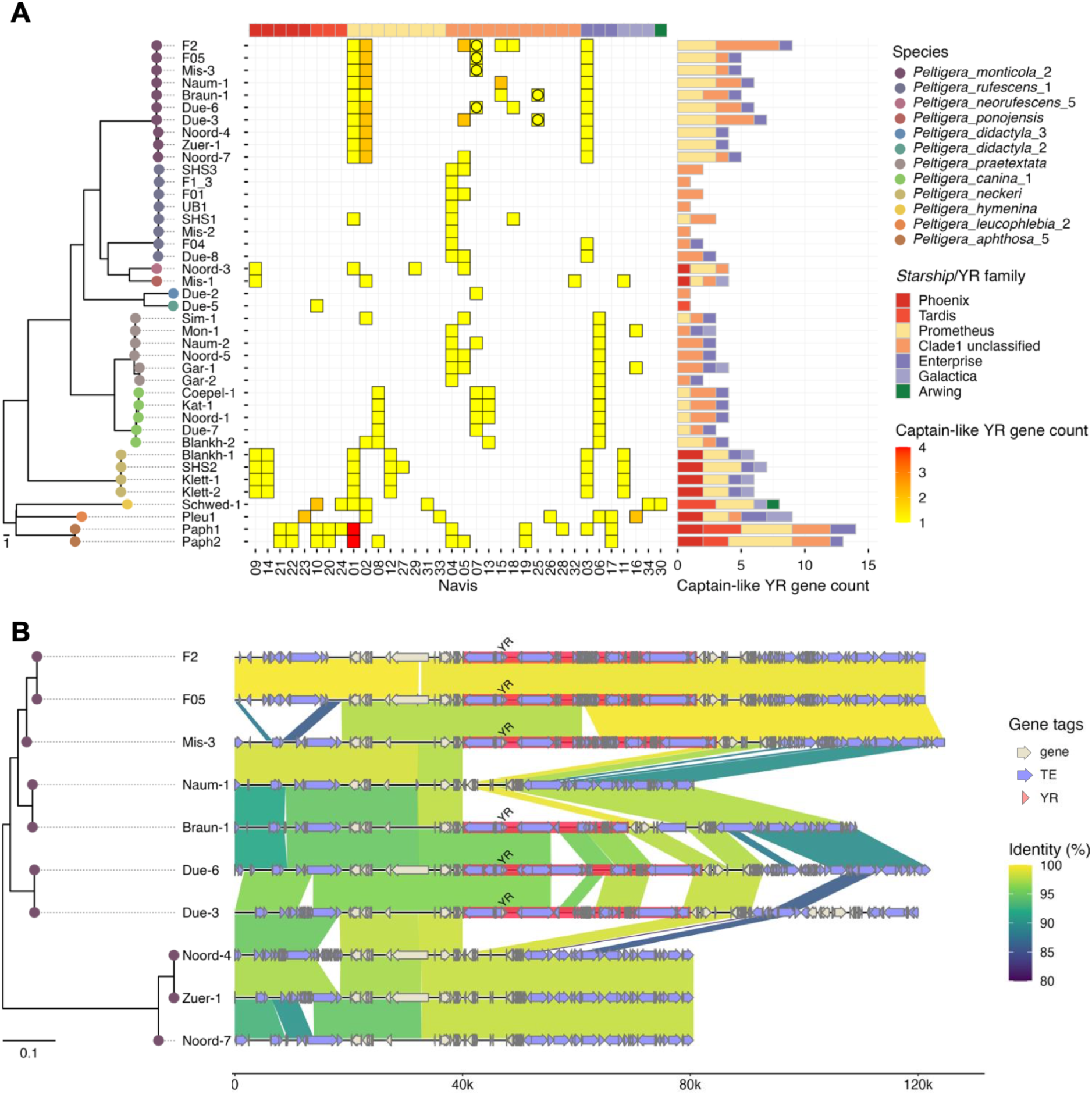
Variable *Starship*-like elements across the *Peltigera* genomes colocalize with transposable elements (TEs). (A) Repertoires of *Starship* captain-like tyrosine recombinase (YR) genes across the 41 *Peltigera* genomes. The tree on the left shows the phylogeny of the 41 *Peltigera* samples, as in Figure 2. The heatmap shows the number of captain-like YR genes in each navis and family, whereas the bar plots on the right show their counts per family. The circle in the heatmap indicates the Starship-like elements highlighted in B. (B) Synteny plots of genomic regions containing *Starship*-like elements in ten samples of *P. monticola_*2. The tree on the left shows the phylogeny of the 41 *Peltigera* samples, as in Figure 2. Syntenic regions with >80% nucleotide sequence identity over >1 kb are connected between genomes. Regions highlighted in red indicate *Starship*-like elements identified using starfish.

Using multiple genome alignments within *P. monticola_2*, we identified six *Starship*-like elements in six of the ten isolates (yellow circles in the main heatmap of Figure 5A). These elements ranged from 29 to 45 kb and were located within an otherwise syntenic chromosomal region, but showed striking insertion–deletion polymorphism across isolates (Figure 5B). The elements were dominated by TE sequences, and in each case contained a captain-like YR gene and only one additional protein-coding gene. However, the YR gene was not positioned directly at the 5’ end of the element but was instead separated from it by intervening TEs, representing a non-canonical *Starship* architecture. In isolates lacking the element (e.g., Noord-4, Zuer-1, Noord-7), the flanking syntenic regions were directly contiguous, confirming that the elements represent genuine insertions rather than assembly artefacts. The co-localization of *Starship*-like elements with other TEs in a variable genomic region reinforces the link between large mobile elements and the repeat-rich, rapidly evolving compartments identified in the two-speed genome analysis above.

### Pangenomic analyses

To systematically characterize gene content variation across the *Peltigera* pangenome, we clustered predicted proteomes into orthogroups using OrthoFinder v3.1.3 with one representative per species and two clades for *P. didactyla*, along with *Solorina crocea* as an outgroup. We then checked for missing orthogroup members by checking for their occurrence in individual genomes. This analysis identified 13,290 orthogroups.

The pangenome was partitioned into core orthogroups (present in all species), soft-core (present in 10–11 of 12 species), shell (present in 2–9 species), and species-specific orthogroups (present in a single species) (Supplementary Figure 6).

The core set comprised approximately 6,963 orthogroups, representing the conserved functional backbone of *Peltigera* genomes. Rarefaction analysis showed that the core genome has largely stabilized at the current sampling depth, whereas the number of species-specific orthogroups continues to increase with each added genome (Supplementary Figure 6, inset), indicating an open pangenome in which lineage-specific gene content remains incompletely sampled across the genus.

A total of 526 species-specific orthogroups were identified. Of these, 58.35% lacked any clear functional annotation, meaning that none of the proteins within these orthogroups could be assigned a putative function based on InterProScan annotations. The amount of private orthogroups broadly mirrored genome size variation. The largest species-specific fractions were observed in *P. aphthosa_5* (158 orthogroups), *P. hymenina* (118 orthogroups) followed by *P. leucophlebia_2* (114 orthogroups). In contrast, species with the smallest genome sizes, including *P. didactyla_2* and *P. didactyla_3*, contained only 11–22 species-specific orthogroups. This comparatively low number of private orthogroups may indicate reduced lineage-specific diversification within sections *Peltigera* and *Horizontales*. (Supplementary Figure 6, Supplementary Table S13).

Beyond core and species-specific content, we identified orthogroups which were shared exclusively among *P. aphthosa_5*, *P. leucophlebia_2*, and *P. hymenina*. Other shared orthogroup sets linked phylogenetically adjacent species within section *Peltigera*. To identify specific biochemical functions that distinguish lineages, we examined InterPro domain presence–absence across species(Blum et al., 2021). Several domains were restricted to the tripartite lichens and *P. hymenina.* These comprised GTP-cyclohydrolase (IPR022163) as well as the virulence factor CipC-like domain (IPR022234) and D-galactonate dehydratase (IPR023592) where the latter also showed a region of similarity also detectable in *P. neckeri*. At the single-species level, Hemicentin/VWA7 galactose-binding domain-like (IPR056475) and Hemicentin-1-like von Willebrand factor A domain (IPR056861) were found exclusively in *P. leucophlebia_2*, while a galactose-binding lectin domain (IPR052487) was unique to *P. aphthosa_5*.

#### Biosynthetic gene clusters

*Peltigera* might show a reduced metabolite richness compared to other members of the Lecanoromycetes, but higher accumulation of specific compounds such as triterpenoids (Ramírez et al., 2025). Terpenoids have been used to classify *Peltigera* species into taxonomic sections (Holtan-Hartwig, 1993; Vitikainen, 1994; Miadlikowska and Lutzoni, 2000). We thus identified biosynthetic gene clusters (BGCs) using antiSMASH (Blin et al., 2025) and grouped these into gene cluster families (GCFs) using BiG-SCAPE for cross-genome comparison. Across most bipartite lichens, we observed a relatively stable repertoire of 33–43 predicted BGCs per genome (Supplementary Table S14). Consistent with its reduced genome size, *P. didactyla* which likely produces little or no terpenoids harboured only 28–29 clusters. In contrast, the tripartite lichens showed a marked expansion: *P. leucophlebia_*2 contained 61 clusters and *P. aphthosa_*5 69 and 71 in its two isolates. *P. hymenina* showed an intermediate position again with the highest BGC count among bipartite species (43 clusters). In line with the metabolic analyses (Ramírez et al., 2025) terpene-associated clusters constituted the dominant BGC class across all genomes, and this was particularly pronounced in *Peltigera* species with tripartite thalli, where terpene BGCs accounted for the majority of the expanded repertoire (Supplementary Figure 7). Despite clear differences in BGC abundance and composition, only a limited number of clusters per genome showed similarity to previously characterized clusters (typically ≤10), suggesting that the majority of *Peltigera* BGCs represent novel or poorly characterized biosynthetic pathways.

Among these, one biosynthetic gene cluster potentially associated with clavaric acid, a lanostane-type triterpenoid, was observed across all 41 genomes. Most genomes harboured a single clavaric acid-like BGC, whereas all *P. rufescens_1* and *P. didactyla* genomes contained two such clusters. In contrast, *P. monticola_2* genomes possessed between three and five BGCs with high similarity to the putative clavaric acid cluster, representing the highest copy numbers observed among the sampled species.

Another notable example was a BGC showing high similarity to the 1,3,6,8-tetrahydroxynaphthalene biosynthetic cluster. 1,3,6,8-Tetrahydroxynaphthalene serves as a precursor in the polyketide pathway (Austin et al., 2004). This cluster was detected exclusively in all samples of *P. rufescens_1*, *P. neorufescens_5*, *P. ponojensis*, and *P. monticola_2*. As these species are phylogenetically close this restricted distribution may represent a phylogenomic signal (Figure 2).

BiG-SCAPE clustering (Draisma et al., 2026) identified 219 GCFs in total. To reduce the influence of sample-specific variation, GCFs detected in only a single sample were excluded, except for species represented by a single genome (*P. leucophlebia_2*, *P. hymenina*, *P. ponojensis*, and *P. neorufescens_5*), resulting in a final dataset of 167 GCFs. These were categorized according to their distribution across genomes into a conserved core of seven GCFs, 31 shared GCFs present across multiple species, 59 species-specific GCFs, and 70 variable GCFs. (Supplementary Figure 7). The presence–absence matrix revealed a marked contrast between the small, universally conserved core and the extensive species-specific and variable fractions, which together accounted for the majority of GCF diversity. BGC repertoires were consistent across isolates of the same species which was visible in the deeply sampled *P. monticola_*2 and *P. rufescens*_1 isolates which showed nearly identical GCF compositions. The species-specific GCFs were concentrated in *P. aphthosa_*5, *P. leucophlebia_*2, and *P. hymenina*, linking the genomic expansion in these lineages to the acquisition of novel cluster families.

#### The genetic inventory of the *Peltigera* mycobiont highlights expanded repertoires of secreted antimicrobial proteins

The molecular mechanisms governing the establishment and maintenance of microbial communities within the lichen thallus remain largely unresolved, particularly regarding the respective contributions of the mycobiont and photobiont to these processes. We hypothesized that antimicrobial proteins (AMPs) play a role in structuring these communities and therefore predicted AMP catalogues for *Peltigera* mycobionts using AMAPEC (Mesny et al., 2026). Across the 41 mycobiont genomes, we identified an average of 306 AMPs per genome, ranging from 27 to 800 amino acids in length, with most proteins falling within 200–400 amino acids and an overall mean length of 321 amino acids (Figure 6A; Supplementary Figure 8). Consistent with the patterns observed for genome size and BGC content, the largest AMP catalogues were identified in the tripartite lichens represented by *P. leucophlebia*_2 (403 AMPs) and *P. aphthosa_*5 (357 and 349 AMPs), whereas the smallest were detected in *P. didactyla_2/3* (252 and 248 AMPs).

**Figure 6.**
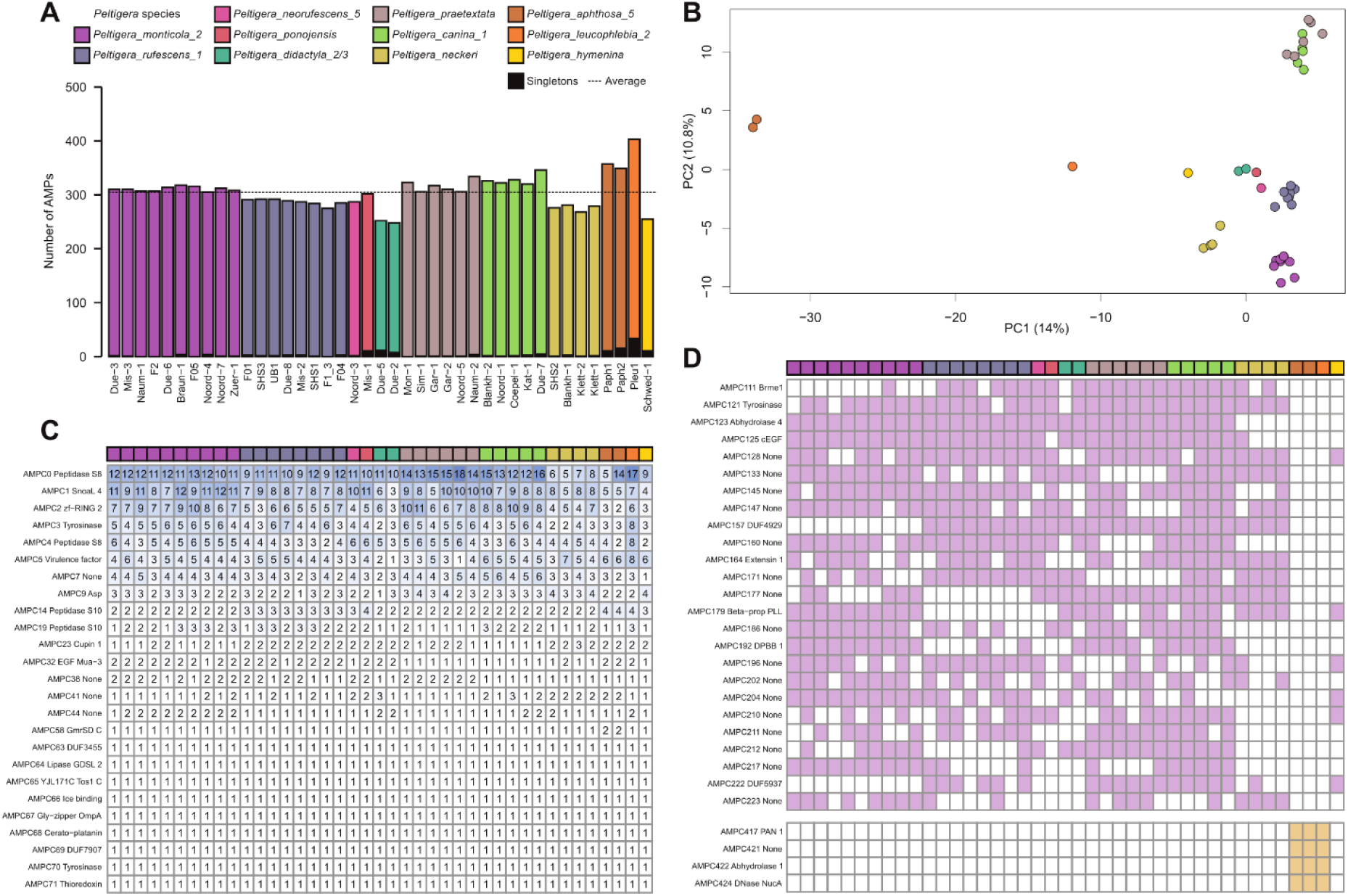
AMP repertoire composition and conservation in *Peltigera* mycobionts. (A) AMP counts and orthogroup assignment across isolates. Bars show total AMP numbers per sample, coloured by *Peltigera* species. Black segments indicate unassigned AMPs, while coloured segments represent AMPs assigned to orthogroups. The dashed line indicates the mean across all 41 samples. (B) Principal component analysis (PCA) of AMP repertoires. Each point represents an isolate based on orthogroup composition and is coloured by *Peltigera* species (color legend in Figure 6A). (C) Conserved clusters across mycobionts. Heatmap showing copy numbers of AMPCs present in all isolates. Colour intensity (white to blue) reflects copy number per genome. Functional annotations are indicated on the left. (D) AMPCs found only in bipartite lichens (top) or tripartite lichens (bottom). Purple indicates presence of an AMPC in a bipartite sample while beige indicates the presence of an AMPC in the tripartite sample and white indicates the absence of an AMPC.

Although AMP numbers differed between species, they were generally stable across isolates of the same species (Figure 6A).

To determine whether AMP repertoires differ between samples, we performed a principal component analysis (PCA). For this, a total of 12,696 predicted AMPs were grouped into Antimicrobial orthologous Clusters (AMPC). Only 199 proteins were unique to a single isolate, while the remaining 12,497 proteins shared sequence similarity with at least one other protein in the dataset, resulting in 505 AMPCs. Most unassigned genes originated from species represented by only one or two samples (Figure 6A). The PCA revealed species-specific AMP profiles, as samples from the same species clustered together. As expected based on phylogenetic relationships of *Peltigera* species sampled for this study (Figure 2), the AMP repertoires of the closely related (Magain et al., 2018) *P. canina_1* and *P. praetextata* clustered closely and were distinct from other species. Tripartite lichens displayed separate clustering patterns, with *P. aphthosa*_5 showing the strongest divergence (Figure 6B).

Among the 25 conserved AMPCs detected across all samples, 16 contained multiple AMP copies in at least one sample. Amongst these, several orthogroups displayed substantial differences in copy number, indicating lineage-specific expansions. For example, AMPC3 (tyrosinase) was present in all genomes but ranged from two to three copies in *P. didactyla_2/3* to eight in *P. leucophlebia*_2 (Figure 6C). Beyond copy-number variation, cluster presence–absence also differed between symbiotic strategies: Four AMPCs were found exclusively in mycobiont genomes of tripartite thalli, whereas 25 were absent from tripartite thalli but present in at least 50% of bipartite thalli (Figure 6D). At the species level, *P. didactyla_2/3*, *P. neorufescens_5*, *P. ponojensis*, and *P. hymenina* lacked unique clusters, whereas *P. aphthosa*_5 harboured 32 (Supplementary Figure S9, Supplementary Table S15).

As antimicrobial proteins can retain similar folds despite extensive sequence divergence, we next assessed whether mycobiont AMPs display conserved folds despite low sequence similarity by clustering predicted structures and comparing the resulting clusters to sequence-based AMPCs. Only ∼1% of structural clusters (26 of 2827) contained proteins from multiple AMPCs, indicating that structural and sequence-based relationships were concordant. The few mixed clusters mainly grouped proteins with similar or unknown functional annotations; for example, representatives of two AMPCs annotated as ricin-B lectins shared low sequence similarity (<25% identity) but high structural similarity (TM-score = 0.71). Only a small number of mixed clusters contained proteins with differing functional annotations (Supplementary Figure S10).

#### *Peltigera* genomes encode abundant orthologs of pathogen–host interaction genes

Because the genome architecture and gene repertoires of *Peltigera* spp. are reminiscent of those of pathogens, we queried the Pathogen–Host Interactions Database (PHI-base; (Urban et al., 2020) to determine whether *Peltigera* genomes indeed harbor orthologs of genes known previously functionally characterized to be involved in pathogen–host interactions. To this end, we reassigned ortholog groups using proteins from the 41 *Peltigera* samples together with proteins from PHI-base. This analysis revealed 2,862 ortholog groups that included PHI-base proteins, indicating that a substantial number of *Peltigera* proteins are orthologous to proteins involved in pathogen–host interactions. These orthologs include many of the genes in BGCs and genes encoding antimicrobial proteins, other secreted proteins, GPCRs, and lectins (Supplementary Table S16).

Next, we analyzed the association of these genes with *Starships*. Our current approach identified only a few *Starship*-like elements in the 41 *Peltigera* genomes (Figure 5B), despite the presence of many captain-like YR genes (Figure 5A). Because *Starship* annotation relies on insertion/deletion polymorphisms flanked by highly conserved syntenic regions around the insertion site, potential *Starships* may have been overlooked in the repeat-rich and variable *Peltigera* genomes. Therefore, we examined genes located within 50 kb downstream of the captain-like YR genes, as these regions may contain potential cargo genes of overlooked Starship-like elements. This analysis revealed that some captain-like YR genes are followed by genes in BGCs and genes encoding antimicrobial proteins, other secreted proteins, and GPCRs (Supplementary Table S17). These genes include orthologs of pathogen genes previously shown to contribute to virulence, such as an antimicrobial gene orthologous to *lipA* in *Xanthomonas campestris* (Tamir-Ariel et al., 2012) and a BGC gene orthologous to ABC3 in *Magnaporthe grisea* (Sun et al., 2006)(Supplementary Table S17). Although it remains to be determined whether these genes are true *Starship* cargo, the proximity of captain-like YR genes to virulence-associated genes is reminiscent of patterns observed in pathogen genomes (Gourlie et al., 2022; Bucknell et al., 2025; Sato et al., 2025).

#### F-Lectin repertoires and fucose-mediated recognition

Lectins are carbohydrate-binding proteins that mediate cell-cell and cell-environment interactions through recognition of surface-exposed glycoconjugates. To define the lectin repertoires of *Peltigera*, we screened the Interproscan functional annotations (Supplementary Tables S6) as well as the Interproscan member databases of the *Peltigera* genomes for lectin-associated InterPro domains. This yielded multiple lectin-related domain families, including Ricin B-, Galectin-, Jacalin-, Legume-, ConA-like, Bulb-type, and Fucose-specific lectins. (Supplementary Table S18, S19). By further filtering data based on sequence similarity to the domains, we identified a core lectin repertoire dominated by Fucose-specific lectins, Ricin B-type lectins, and Galectins, which were present in all examined genomes. Fucose-specific lectins represented the most abundant lectin family across the *Peltigera* genomes, with a total of 355 genes identified among the 41 analyzed genomes. Individual genomes contained between 5 and 17 Fucose-specific lectin genes, highlighting this family as a prominent and highly conserved component of the *Peltigera* lectin repertoire.

To link lectin repertoires to potential binding targets in the photobionts we performed cell-wall carbohydrate compositional analysis of the photobiont and medulla layers in three *P. rufescens_1* thalli (F04, SHS3, and UB1). Mannose, glucose, and galactose were abundant in both layers, but fucose was consistently detected only in the photobiont layer and was absent from the medulla (Supplementary Figure 11A; Supplementary Table S20, S19). This is supported by prior observations where the *Nostocs* exopolysaccharides contain fucose (De Philippis et al., 2000) as well as fucosylated carotenoids (Takaichi et al., 2009) to test whether fungal tissues produce proteins capable of binding fucose, we stained cross sections of an intact *Peltigera* thallus with a fluorescent fucose glycoconjugate recognized by such fucose-binding lectins. Fucose based fluorescence bound predominantly to the cortex and medulla layers, while the photobiont layer showed weaker signals (Supplementary Figure 11B). This might suggest that fucose-specific lectins may mediate recognition of fucose-containing epitopes at the mycobiont-photobiont interface.

#### An extensive G-Protein coupled receptor repertoire is found across *Peltigera* isolates

G protein–coupled receptors (GPCRs) represent a central class of eukaryotic signaling proteins that translate extracellular cues into intracellular responses and are therefore key determinants of environmental sensing and developmental regulation. In filamentous fungi, GPCRs are particularly important for coordinating growth, morphogenesis, and interactions with other organisms, yet their diversity and evolutionary dynamics in lichen-forming fungi remain poorly characterized (Heinen et al., 2025). We therefore generated a comprehensive overview of GPCRs across all *Peltigera* isolates using a structure based approach shown to substantially increase GPCR recovery in fungi (Mendoza-Rojas et al., 2025). To validate this strategy for lichen-forming ascomycetes, we first re-analyzed the GPCR repertoire of *Cladonia grayi* and compared our results to previous annotations (Liu et al., 2025). Using our structural pipeline, we identified 128 GPCR candidates, greatly exceeding the previously reported set of 26 GPCRs and indicating that sequence-based approaches also substantially underestimate receptor diversity in lichen fungi (Supplementary Table S21 and S22).

We subsequently applied the pipeline to all *Peltigera* genomes using a multi-threshold TM-score cascade and identified a total of 4,214 putative GPCR candidates across all 41 genomes. Individual genomes encoded, on average, approximately 103 GPCRs (range: 84–119), indicating a remarkably conserved receptor repertoire across the sampled thalli. Together with the results from *C. grayi*, these findings suggest that lichen-forming fungi generally possess substantially larger and comparatively stable GPCR inventories than previously recognized.

To classify these GPCR candidates, we used representative structures from the 17 GPCR classes defined by Liu et al. (Liu et al., 2025). Of all predicted GPCRs, 3,860 (91.6%) were resolved unambiguously to a single GPCR class, while 354 candidates (8.4%) remained structurally ambiguous across all thresholds (Figure 7A). Most classes were represented by only a few candidates per genome. However, based on the Liu et al. (Liu et al., 2025) classification scheme, 2,934 of 3,860 resolved candidates (76%) were assigned to Class 14 (Pth11-like/CFEM-domain-containing receptors), with per-genome copy numbers ranging from 37 to 95 (Figure 7B, Supplementary Tables S23, S24). This marked expansion is consistent with observations across the Pezizomycotina, where Pth11-like receptors constitute the largest and most dynamic fungal GPCR family (Xu et al., 2017; Armaleo et al., 2019; Liu et al., 2025). Considering receptor numbers reported for other fungi, however, available genome annotations have likely underestimated the full extent of Pth11 repertoires (Liu et al., 2025).

**Figure 7:**
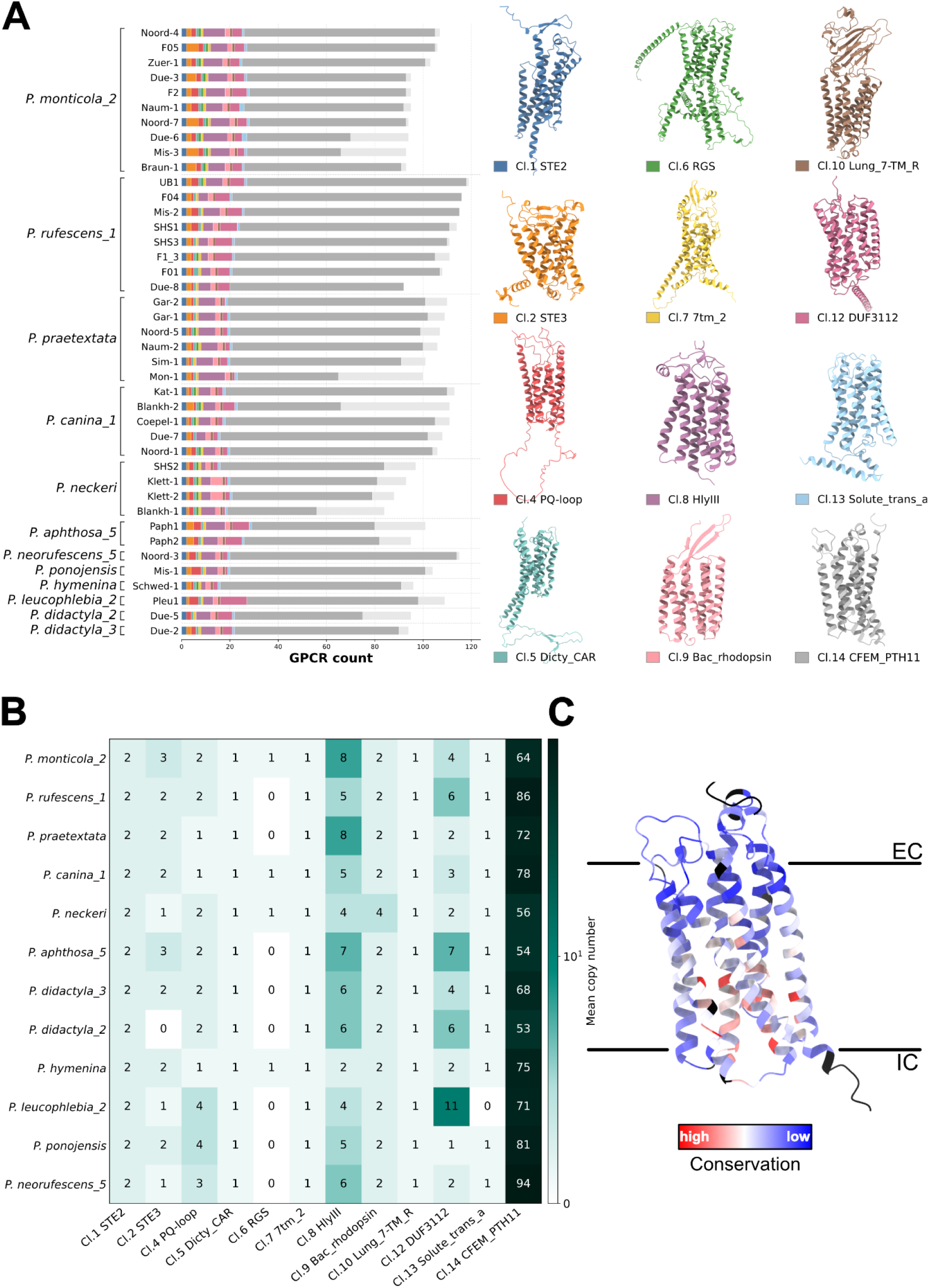
GPCR repertoire composition, class distribution, and structural conservation across *Peltigera* lichen mycobionts. (A) Stacked bar charts depicting the GPCR class composition (Cl.1–Cl.14, color-coded) per sampling site across 11 *Peltigera* species. Total GPCR count is displayed on the x-axis (0–120), with representative structural models for each class shown to the right in the corresponding color. (B) The heatmap displays the mean GPCR copy numbers per genome per class and species. Values depicted are rounded to the next full number, SymLog scale coloring was chosen to keep resolution for lower copy number classes, while still showcasing CFEM_Pth11 expansion correctly. (C) A representative 7-TM GPCR structural model colored by sequence conservation (blue = variable; red = conserved) was derived from a multi-species MSTA of all putative Cl. 14 CFEM_Pth11 receptors, with one representative genome from each species. The color scale highlights conserved transmembrane core regions relative to variable extra- and intracellular loops, with the extracellular (EC) and intracellular (IC) faces indicated.

Beyond class 14 (Pth11-like GPCRs), classes 8 (HlyIII) and 12 (DUF3112) showed increased copy numbers, encompassing potential orthologs of *Aspergillus* and *Neurospora* PTH- and NOP-type receptors and proteins of the DUF3112 family, respectively. For some species, we observed increased overlap between these class assignments, suggesting that the structural boundaries between classes 8, 12, and 14 might be relatively subtle. This ambiguity may reflect either evolutionary relatedness among these classes or limitations of current fungal GPCR classification schemes.

Given the striking abundance of Pth11-like GPCRs in *Peltigera*, our data suggest that, as in plant-pathogenic fungi, these receptors likely play a central role in environmental sensing and in mediating interactions within the lichen symbiosis. To further examine this receptor family, we structurally superposed a broad repertoire of Pth11-like receptors, generated a multiple structural alignment (MSTA) using FoldMason (Gilchrist et al., 2026), and mapped residue conservation onto one representative candidate from *P. rufescens_1* (Figure 7C). Despite pronounced structural conservation across the Pth11 family revealing a common architecture, this analysis highlighted a noticeable sequence diversity explaining why sequence-based approaches to identify GPCRs have missed a large portion of the Pth11 candidates. Mapping sequence conservation onto this structural scaffold showed that residues lining the extracellular-facing surface are highly divergent, whereas a subset of residues within the transmembrane core remains conserved. This pattern is consistent with diversification of ligand-recognition surfaces combined with conservation of the structural core required for receptor stability and signal transduction (Venkatakrishnan et al., 2013; Hauser et al., 2021) (Figure 7C).

In addition to the expanded Pth11-like receptor repertoire, we observed a highly conserved presence of two slightly different Ste2/class 1 pheromone receptors across all *Peltigera* isolates (Figure 7A, B). These paralogs were consistently located on chromosomes 7 and 10, respectively, indicating that the duplication predates the diversification of the sampled *Peltigera* lineages and has been maintained over substantial evolutionary time (Supplementary Figure 12). The conservation of both paralogs across all isolates suggests that this duplication is unlikely to represent recent copy-number variation or assembly artefact. Instead, it points to long-term retention, potentially reflecting functional diversification within the pheromone-sensing pathway. Such diversification could involve differences in ligand specificity, expression pattern, mating-type association, or additional roles outside canonical mating, as observed for the Ste2 pheromone receptor *Fusarium oxysporum (Vitale et al., 2019)*.

Collectively, our results reveal a receptor landscape in *Peltigera* that is dominated by extensive expansion of Pth11-like GPCRs, accompanied by more limited diversification of additional receptor classes. In disagreement with previous studies, we reveal a strong expansion of Pth11 GPCRs in lichen-forming fungi suggesting that expansion is not limited to pathogens but likely reflects a general evolutionary theme in Pezizomycotina fungi.

### Spatial transcriptomics reveals layer-specific deployment of interaction genes

To test whether the gene families identified through comparative genomics are specifically expressed in different layers of the lichen thallus, we performed layer-specific transcriptome profiling. An additional *P. rufescens* specimen from the UB1 sampling site was dissected into a combined cortex/photobiont layer, the medulla, and the rhizines (Figure 8A), followed by RNA-sequencing. Differential expression analysis identified 1,256 differentially expressed genes between the cortex/photobiont layer and the medulla, 2,350 between the cortex/photobiont layer and the rhizines, and 564 between the medulla and rhizines (Supplementary Table S25). InterPro domain enrichment analysis of genes upregulated in the cortex/photobiont layer relative to both the medulla and rhizines revealed a pronounced enrichment of PTH11-like integral membrane proteins, Rhodopsin domains (both corresponding to GPCR class 14), and Peptidase S8/S53 family proteins (Figure 8C). PTH11-like receptors represented the most strongly enriched domain family in both comparisons suggesting a role in signalling and interaction. Examination of the top 70 differentially expressed genes between the cortex/photobiont layer and the medulla (Figure 8B, Supplementary Table S26) showed a similar pattern: among the most strongly upregulated genes were two Pth11 GPCRs and an RGS domain-containing protein, alongside several AMP protein candidates including Peptidase encoding genes. Focussing on the spatial expression of antimicrobial protein candidates indeed revealed that multiple Peptidase S8/S53 domain proteins were strongly and specifically upregulated in the cortex/photobiont layer, whereas other AMP candidates, including a SnoaL-like domain protein, an aspergillopepsin-like endopeptidase, and tyrosinase copper-binding domain proteins, showed higher expression in the medulla or rhizine layers (Figure 8E; Supplementary Table S27).

**Figure 8:**
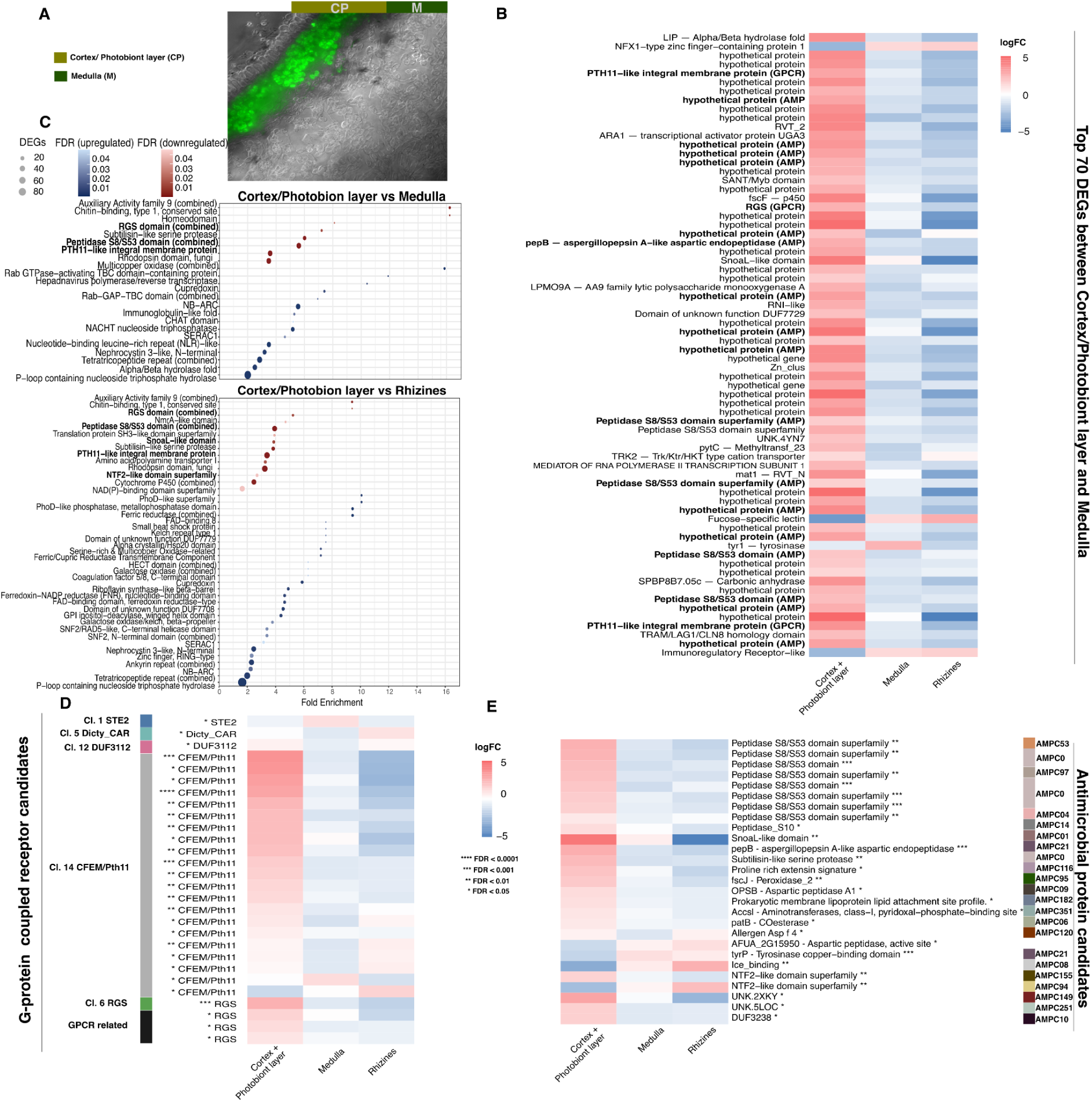
Layer-specific transcriptome analysis of the *Peltigera* thallus. Representation of the dissection strategy of the *Peltigera* thallus into defined layers. Due to the compact organization of the upper cortex and photobiont layer, both were combined for downstream analyses (A). Additional layers include the medulla and rhizines (not shown). (B) Heatmap of the top 70 significantly differentially expressed genes (DEGs) between the cortex/photobiont layer and the medulla. Bold terms indicate proteins belonging to subcategories of antimicrobial peptides (AMPs) and G-protein–coupled receptors (GPCRs). (C) Overrepresentation analysis of functional categories among DEGs. Bold terms highlight categories associated with potential AMP and GPCR functions.(D) Magnified view of GPCR candidates showing significantly differentially expressed genes between the cortex/photobiont layer and the medulla. Assigned GPCR classes are indicated by colored bars. RGS labels include one candidate corresponding to a class 6 RGS GPCR, whereas the remaining candidates represent GPCR-related RGS proteins lacking the characteristic seven-transmembrane helix structure. Stars indicate statistical significance: **** FDR < 0.0001, *** FDR < 0.001, ** FDR < 0.01, * FDR < 0.05. (E) Visualization of significantly differentially expressed antimicrobial protein candidates. Assigned AMPCs are represented by colored bars. Significant AMP-related DEGs annotated as “hypothetical protein” are not shown. Stars indicate statistical significance: **** FDR < 0.0001, *** FDR < 0.001, ** FDR < 0.01, * FDR < 0.05

Additional genes associated with extracellular interaction and secondary metabolism were also enriched in the cortex/photobiont layer. These included tyrosinases (*tyr1*, *tyrP*), a cytochrome P450 monooxygenase, galactose oxidase (*GAOA*), and the siderophore biosynthesis enzyme *SidA*. Extracellular enzymes implicated in protein and polysaccharide degradation included subtilisin-like serine proteases, an AA9 lytic polysaccharide monooxygenase (*LPMO9A*), and a glycoside hydrolase family 18 enzyme. Transporters, including the sugar: H⁺ symporter *mstC* and the Trk-type cation transporter *TRK2*, further pointed to altered nutrient acquisition and ion homeostasis across thallus layers. Interestingly, a substantial fraction of the most highly differentially expressed genes encoded hypothetical proteins (Figure 8B), underscoring that much of the transcriptional activity at the symbiotic interface remains functionally uncharacterized.

We next assessed whether potential lectin-encoding genes exhibited layer-specific expression patterns. For this analysis, we used the filtered lectin dataset (Supplementary Table S19) and evaluated differential gene expression. Among the identified lectins, we detected homologs of the previously described LEC-1 and LEC-2 genes from *Peltigera membranacea*. These galectin/galactose-binding lectins have been proposed to mediate recognition of ligands produced by the *Nostoc* photobiont (Miao et al., 2012). The gene mikado.UB1_chr02G942.1 showed 97% query coverage and 47.0% amino acid identity to the LEC-1 protein (AGC12988.1). Notably, this gene was significantly upregulated in the Cortex/Photobiont layer (FDR < 0.01) and downregulated in rhizines, consistent with a potential role in photobiont recognition and interaction.

A second lectin candidate, mikado.UB1_chr02G287.1, showed strong similarity to LEC-2 (AGC54691.1) but was not expressed in our dataset. In contrast, another lectin candidate displaying similarity to both a galectin from *Sphaerosporella brunnea* and *P. membranacea* LEC-2 showed elevated expression in the cortex/photobiont layer and reduced expression in rhizines, albeit no statistically significant differences. Given the high abundance of fucose-specific lectins in the *Peltigera* mycobiont genomes, we next examined their expression patterns in more detail. Among the nine predicted fucose-specific lectins in *P. rufescens_1* genome UB1, one candidate displayed elevated expression in rhizines and reduced expression in the Cortex/Photobiont layer. A similar pattern was observed among the top 70 differentially expressed genes (Fig. 8B), where the only detected fucose-specific lectin exhibited the same trend. In contrast, a second high-confidence fucose-specific lectin candidate showed the opposite expression pattern, with increased expression in the Cortex/Photobiont layer and reduced expression in rhizine. All fucose-specific lectins recognize fucosylated epitopes, but their binding specificity is influenced by neighboring, subterminal sugar moieties, such as galactosyl and/or N-acetylglucosamine residues (Vasta et al., 2017). Therefore, the different fucose-specific lectins might recognize distinct glycan epitopes present only in the mycobiont/photobiont and/or provide a first line of recognition for non-symbiontic microbes. A comparable divergence in expression profiles was also observed among Ricin B lectins, with different candidates exhibiting contrasting layer-specific expression patterns. Together, these results suggest that, unlike the galectins, which displayed a consistent enrichment in the Cortex/Photobiont layer, other lectin families such as fucose-specific lectins and Ricin B lectins may fulfill more diverse and potentially specialized functions within the lichen symbiosis. (Supplementary Table S28).

Finally, we analyzed the spatial deployment of GPCR signaling components. InterPro domain enrichment revealed that PTH11-like integral membrane proteins and Rhodopsin domains, both corresponding to GPCR Class 14, were among the most enriched domain families in genes upregulated in the cortex/photobiont layer relative to both the medulla and rhizines (Figure 8C). Mapping GPCR expression across thallus layers showed that most differentially expressed Pth11-class GPCRs were upregulated in the cortex/photobiont layer (Figure 8D; Supplementary Table S29), consistent with a role in sensing local symbiotic or environmental signals. By contrast, the two Ste2 pheromone receptor paralogs and RGS regulators showed contrasting layer-specific expression patterns, suggesting functional differentiation among GPCR classes and between duplicated canonical signaling components.

Together, the layer-specific transcriptome supports the spatial deployment of the interaction-associated repertoires identified by comparative genomics. Secreted proteins and AMP candidates, lectins, GPCRs, extracellular enzymes, transporters, and secondary metabolism-associated genes are differentially expressed across the cortex/photobiont layer, medulla, and rhizines, indicating that the molecular interaction toolkit of the *Peltigera* mycobiont is organized not only at the genomic level but also shows defined transcriptional profiles across the differentiated architecture of the lichen thallus.

## Discussion

### Chromosome-scale genomes establish a foundational resource for lichen genomics

Here, we present a comprehensive resource for the lichen genus *Peltigera*, comprising 41 isolates representing 11 different species. We provide chromosome-scale assemblies for thirty samples representing seven species including telomere-to-telomere (T2T) assemblies for five isolates representing four species. To achieve this, we combined high-accuracy PacBio HiFi and long Nanopore sequencing data, complemented by Pore-C chromosome conformation data for multiple samples relying on integrating the strength of the different sequencing platforms (van Rengs et al., 2022). Extensive intraspecific sampling for *P. rufescens*_1 and *P. monticola*_2 revealed high levels of within species synteny, further supporting assembly accuracy and structural consistency. Our assembly exhibited high per-base accuracy and gene completeness. Merqury QV scores frequently exceeded 60 and reached 76.17 for *Peltigera rufescens_*1 F1_3, while at least one assembly per species exceeded 98.2% BUSCO completeness.

Compared with currently available lichen genome resources, our assemblies substantially improve contiguity and completeness. The largest comparative dataset to date (Resl et al., 2022) broadly sampled 46 Lecanoromycetes but mostly relied on short-read sequencing, resulting in relatively fragmented assemblies with N50 values typically below 300 kb. More recent individual lichen long read assemblies, including *Ramalina farinacea (Llewellyn et al., 2023)* or *Bacidia gigantensis* (Allen et al., 2021), improved continuity but remained below chromosome scale which were reached only recently in some assemblies from *Acarospora socialis* and *Cladonia uncialis (Adams et al., 2023; Dong et al., 2026)*.

Our precise assemblies revealed that *Peltigera* genomes are substantially larger than those of many other lichen-forming fungi, with most species reaching approximately 120 Mb, far exceeding the commonly observed 20–60 Mb range. Genome size variation among *Peltigera* genomes was however pronounced, ranging from only 56–59 Mb in *P. didactyla* to 137.6 and 145.5 Mb in *P. hymenina* and *P. aphthosa_*5, respectively. The latter variation seems to represent TE contraction in the *P. didactyla* lineage (Figure 1). This might be driven by the specialized lifestyle of *P. didactyla* which is relatively fast growing, always features apothecia and produces soredia (i.e. propagules containing both myco- and cyanobiont). Potentially this leads to increased meiosis rate which can e.g. trigger repeat induced point mutations to inactivate TEs (Horns et al., 2012; Gladyshev, 2017) or simply based on recombination events, as non-recombing mate type chromosomes have been shown to accumulate TEs (Duhamel et al., 2023).

Furthermore the TE driven genome size variation is consistent with the established role of TEs as major determinants of fungal genome size evolution (Möller and Stukenbrock, 2017). Interestingly we also detected piggyBac TIR transposable elements exclusively in tripartite genomes which was also supported by the interpro analysis (Supplementary Table S6), while piggyBac TIRs are not widespread in the ascomycota, they have been detected in some *Fusarium* isolates (Laxmi et al., 2023; Srivastava et al., 2023).

In terms of genes, we found that the small genome of *P. didactyla* had somewhat fewer genes (9,992–10,295) compared to 10,835-14,309 for other *Peltigera* species, and lacked species-specific (private) AMPs. On the other hand, the two tripartite species *P. leucophlebia*_2 and *P. aphthosa*_5 exhibited expanded molecular inventories across multiple gene categories. Their genomes harboured the highest numbers of biosynthetic gene clusters with 71 and 69 in *P. aphthosa*_5 and 61 in *P. leucophlebia*_2 versus 33–43 in bipartite *Peltigera* lichens, with a pronounced expansion of terpene-associated clusters and general high diversity reflecting the diversity in terpenes produced by most species from the genus *Peltigera (Miadlikowska and Lutzoni, 2000; Ramírez et al., 2025)*.

Tripartite *Peltigera* lichens encoded the largest AMP catalogues in our dataset (349–403 predicted AMPs compared with a mean of 306 across all samples). In addition, they possessed several AMP orthogroups that were absent from all bipartite *Peltigera* species, indicating a distinct AMP repertoire associated with this symbiotic strategy. More broadly, the extensive AMP repertoires identified across *Peltigera* suggest that antimicrobial proteins may contribute to the antimicrobial activity previously reported for members of this genus and may influence the composition of their associated bacterial communities. These protein-based mechanisms could complement the diverse antimicrobial secondary metabolites known from *Peltigera* lichens (Ingólfsdóttir et al., 1985; Leiva et al., 2016; Ramírez et al., 2025). The biological basis of these expansions might be driven by the increased complexity of the tripartite symbiosis.

### Diversified molecular interaction repertoires distinguish lichen symbiotic strategies

*Peltigera* mycobionts showed pronounced interspecific variation ranging from genome size, TE content to gene content and functions, with the tripartite species consistently harbouring the most expanded repertoires. The BGC landscape mirrors the genomic patterns where tripartite thalli harbour nearly twice as many clusters as *Peltigera* species forming bipartite thalli, a pattern driven primarily by terpene BGCs.

The AMP repertoires identified using the recently developed AMAPEC selector (Mesny et al., 2026), in *Peltigera* suggest that antimicrobial proteins may represent an important and previously overlooked component of the molecular toolkit used by lichen. AMP profiles were highly conserved within species yet differed markedly among species, indicating that these repertoires are evolutionarily stable traits rather than highly variable genomic features. The presence of lineage-specific clusters and copy-number expansions further suggests that AMP repertoires have undergone diversification during *Peltigera* evolution, potentially reflecting adaptation to distinct ecological niches or microbial associates. The strong correspondence between sequence- and structure-based clustering indicates that this diversification extends beyond simple sequence divergence and largely involves structurally distinct protein families. The lineage specific AMPs further highlights the uniqueness of these repertoires and suggests that lichen-forming fungi may represent a largely unexplored reservoir of antimicrobial protein diversity. Given the proposed role of AMPs in mediating microbial interactions in other symbiotic systems, these findings raise the possibility that species-specific AMP repertoires contribute to the assembly and maintenance of lichen-associated microbiomes. Indeed, we found that these profiles seem to differ between symbiotic strategies: four orthogroups are exclusive to tripartite thalli, while 25 are restricted to species forming bipartite thalli.

*P. hymenina* as the closest bipartite relative of the tripartite clade and sharing a common ancestor with the tripartite group, consistently was placed between the two groups with the largest genome and highest BGC count among bipartite species, and a large AMP compendium. This might suggest that the genomic expansions associated with tripartite symbiosis accumulated gradually rather than arising at the transition itself, or that the common ancestor was tripartite and some species lost their algal associations. In both cases this makes this clade and potentially the whole *Polydactylon* section very interesting for further studies to understand tripartite versus bipartite lichens.

Our structure-based annotation pipeline identified approximately 103 GPCRs per *Peltigera* genome and 128 GPCRs in *Cladonia grayi*, greatly exceeding the previous sequence-based estimates of 26 GPCRs (Liu et al., 2025). Approximately three quarters of the classified receptors belonged to the Pth11-like family, and inclusion of receptors related to classes 8 and 12 increased this proportion to more than 90% of the total GPCR repertoire. Historically, Pth11-like receptors were discovered in the rice blast fungus Pth11 (DeZwaan et al., 1999) and subsequently attracted attention because large repertoires appeared characteristic of filamentous plant pathogens. They were originally considered important to mediate host and host surface sensing but presence in saprophytes and entomopathogens suggested further roles in general environmental sensing (Kulkarni et al., 2005; Gruber et al., 2013; Xu et al., 2017). The extensive receptor repertoires identified in lichenized fungi now demonstrate that expansion of this family is not restricted to pathogenic lifestyles. Instead, our findings suggest that previous sequence-based approaches substantially underestimated the abundance of these highly divergent receptors. The predominance of Pth11-like GPCRs in *Peltigera* and their expression pattern supports a broader role in sensing complex extracellular environments and mediating interactions with symbiotic partners, which also suggests that the Pth11 repertoires of other Pezizomycotina fungi might be larger than anticipated so far.

Residues predicted to participate in intracellular signal transduction were considerably more conserved than extracellular regions, suggesting that receptor diversification primarily affected signal recognition while preserving downstream signalling mechanisms. Such a pattern is reminiscent of metazoan GPCR evolution, where diversification of ligand-binding interfaces is accompanied by conservation of G-protein coupling determinants (Strotmann et al., 2011; Zhou et al., 2019). Residues revealed by our analysis might therefore be involved in transmitting conformational dynamics within the seven-transmembrane core.

Our results raise the possibility that Pth11-like receptors are not merely an expanded GPCR family of Pezizomycotina, but instead represent their principal receptor lineage suggesting that the majority of GPCR-mediated signalling in filamentous ascomycetes may ultimately be attributable to the diversification of the Pth11 family. Therefore, we propose that the diversification of the Pth11 family represents a central evolutionary strategy by which Pezizomycotina fungi adapts extracellular sensing to distinct ecological niches while maintaining a conserved intracellular signalling framework.

### Two-speed genome architecture and mobile elements contribute to symbiotic adaptation in *Peltigera*

Taking our results together, within the *Peltigera* mycobiont genomes, we identified a typical hallmark of “two-speed genomes”: the localization of secreted protein genes in gene-sparse, TE/repeat-rich regions. *Peltigera* genomes are highly enriched in TEs/repeats, resembling those of some filamentous fungal plant pathogens (Raffaele and Kamoun, 2012). Together with the occurrence of a similar “two-speed genome” organization in the plant-beneficial AM symbiont (Oliveira et al., 2025), our results suggest that this organization is associated with various types of symbioses. Although the evolutionary mechanisms underlying the preferential proximity of TEs to secreted protein genes in the lichen-forming fungi remain to be elucidated, this pattern may reflect potential molecular inventory races between the *Peltigera* mycobiont and its symbionts, analogous to the molecular arms races between pathogens and their hosts (Seidl and Thomma, 2017). Indeed, our study reveals a wealth of antimicrobial proteins in the mycobionts, suggesting that symbiotic partners may need to employ immune surveillance to balance their symbiotic relationships. Microbial immune systems targeting other microbes remain poorly understood compared with mammalian and plant immune systems. Thus, lichen-forming fungi may provide a unique platform for studying potential arms races in microbial symbiosis and their relevance to genome compartmentalization. In addition to secreted protein genes, TEs have also been found near genes associated with abiotic stress responses and get activated in *P. britannica* upon heat stress (Almer et al., 2023), rather than near genes essential for cellular activity, in diverse Peltigerales fungi (Cameron et al., 2026). Together with our findings, this further supports genome compartmentalization and its potential relevance to environmental adaptation in lichen mycobionts.

In line with this symmetry to some pathogenic fungi, our study reveals *Starship*-like elements in which captain-like YR genes are followed by abundant TEs in *Peltigera* mycobionts. These elements are variable among the samples within the same species, analogous to the localization of *Starships* in the plastic genomic regions of filamentous pathogenic fungi infecting animals or plants (Gluck-Thaler et al., 2025; Sato et al., 2025). Although it remains unclear whether these elements are mobile as intact *Starships*, owing to the non-canonical position of captain-like YR genes (i.e., not immediately at the 5’ end), our data show an association between *Starships* and other TEs, analogous to patterns observed in filamentous fungal plant and animal pathogens (Sato et al., 2025; Griem-Krey et al., 2026). In these fungal pathogens, active TEs have been proposed to be introduced via “hitchhiking” on horizontally transferred *Starships*. Similarly, a *Starship* in the lichen-forming fungus *Xanthoria parietina* also carries TEs and has been implicated in horizontal transfer between its symbiotic fungi (Tagirdzhanova et al., 2025a). Because we did not identify canonical *Starships* with a captain gene precisely at the 5’ end, the exact *Starship* cargo contents and their relationship with two-speed genome architecture in *Peltigera* remain to be clarified. However, detecting *Starships* in the repeat-rich and highly variable *Peltigera* genomes is likely challenging, as current *Starship* identification depends on presence–absence patterns in conserved syntenic regions surrounding insertion sites (Gluck-Thaler and Vogan, 2024), which are typically obscured by subsequent genomic rearrangements that are frequent in TE- and *Starship*-containing regions (Faino et al., 2016; Gluck-Thaler et al., 2022; Sato et al., 2025). Nevertheless, the widespread occurrence of lineage-specific captain-like YR genes suggests that additional, currently overlooked *Starships* associated with plastic genomic regions, may be present in *Peltigera* genomes. Hence in total we detected broad interaction repertoires in mutualistic fungi which might evolve through genomic principles similar to those known from pathogens.

## Material and Methods

### Lichen Sampling and Preparation

Three lichen samples were obtained from previous studies. Lichen samples of the genus *Peltigera* were collected from 16 sites across five European countries. Samples were collected in the field using sterile scalpel blades and individually transferred to labeled sterile bags. Geographic coordinates and sampling metadata were recorded at the time of collection (Supplementary Table S1). Samples were transported to the laboratory in a cooled insulated container and processed immediately. In the laboratory, lichen thalli were rinsed with running water to remove adhering soil and debris. Residual moss and stones were removed using tweezers. Cleaned material was blotted dry using laboratory tissue, flash-frozen in liquid nitrogen, and stored at −80 °C until further analysis.

### Oxford Nanopore and PacBio Hifi Sequencing

Approximately 0.5–1 g of frozen lichen thallus material was pulverized to a fine powder using a ball mill. Prior to grinding, steel milling containers were decontaminated with RNase Away (Thermo Fisher Scientific, Waltham, MA, USA) to remove residual nucleic acids. High-molecular-weight (HMW) DNA was extracted using the NucleoBond HMW DNA extraction Kit (Macherey-Nagel, Germany) as described previously for plants (van Rengs et al., 2022; Ziegler et al., 2025). DNA integrity was assessed by 0.8% agarose gel electrophoresis. DNA purity was measured using a NanoDrop spectrophotometer, and DNA concentration was determined using the Qubit DNA assay kit and Qubit fluorometer (Thermo Fisher Scientific, Waltham, MA, USA). The quality and quantity of extracted DNA were assessed using a NanoDrop spectrophotometer (Thermo Fisher Scientific, Waltham, MA, USA), the Qubit BR DNA assay (Thermo Fisher Scientific, Waltham, MA, USA), and the Femto Pulse system (Agilent Technologies, Santa Clara, CA, USA) with the FP-1002 protocol (165 kb) to evaluate fragment size distribution. When DNA purity metrics fell below recommended thresholds, additional clean-up steps were performed. DNA was precipitated using 0.15× volume of 3 M sodium acetate (pH 5.5) and 2× volumes of 100% ethanol, followed by centrifugation and three wash steps with 80% ethanol, each including centrifugation. If contamination persisted, further purification was performed using the DNeasy PowerClean Pro Cleanup Kit (Qiagen, Hilden, Germany). For Oxford Nanopore Sequencing approximately 1.5 µg of DNA was used for library preparation. DNA repair and 3′-adenylation were performed using the NEBNext FFPE DNA Repair Mix and the NEBNext Ultra II End Repair/A-Tailing Module (New England Biolabs, Ipswich, MA, USA). Libraries were prepared using the SQK-LSK114XL kit and sequenced on FLO-PR0114M flow cells (Oxford Nanopore Technologies, UK) for 100 h. For sample F1_3, three sequencing runs were performed using the SQK-LSK112 kit on FLO-PRO112 flow cells (Oxford Nanopore Technologies, UK). PacBio Hifi Sequencing was performed on the Sequell and Revio system (Pacific Biosciences of California, Inc., California, USA).

### Oxford Nanopore cDNA library construction and Sequencing

To support annotation of the mycobiont genome, cDNA sequencing was performed. cDNA libraries were generated from three *Peltigera rufescens*_1 samples (F1_3, F01, F04) and two *Peltigera monticola*_2 samples (F2, F05). In addition, a separate *P. rufescens*_1 thallus collected at the Blankenwasser Ufer B sampling site was used. Total RNA was extracted from all samples using TRIzol reagent according to the manufacturer’s instructions. RNA integrity was assessed by 0.8% agarose gel electrophoresis. RNA purity was measured using a NanoDrop spectrophotometer, and RNA concentration was determined using the Qubit RNA assay kit and Qubit fluorometer (Thermo Fisher Scientific, Waltham, MA, USA). cDNA libraries were prepared using the SQK-PCS111 kit selecting for polyA tailed RNA and sequenced on an FLO-MIN116 flow cell (Supplementary Table S30).

### Oxford Nanopore PoreC library construction and Sequencing

To generate chromosome-scale genome assemblies of the mycobiont, Pore-C sequencing data were generated for species with sufficient biomass. Pore-C data were obtained for *P. rufescens*_1 (F1_3, F01, F04), *P. monticola*_2 (F2, F05), *P. praetextata* (Sim-1), *P. canina*_1 (Coepel-1), *P. neorufescens_5* (Noord-3), and *P. neckeri* (Blankh-1). For each sample, 1-2 g of whole lichen thallus material was cut into ∼2 cm² fragments and subjected to vacuum infiltration. Pore-C libraries were prepared following the Plant Pore-C protocol (Oxford Nanopore Technologies, UK). DNA purity was assessed using a NanoDrop spectrophotometer, and DNA concentration was determined using the Qubit DNA assay kit and Qubit fluorometer (Thermo Fisher Scientific, Waltham, MA, USA). When DNA purity metrics fell below recommended thresholds, additional clean-up steps were performed as described in section Oxford Nanopore and PacBio Hifi Sequencing. Libraries were constructed using the SQK-LSK114XL sequencing kit and sequenced on FLO-PRO114M flow cells. When sufficient library material was available, flow cells were washed using the EXP-WSH004 (Oxford Nanopore Technologies, UK) wash kit and reloaded for a second sequencing run.

### Mycobiont genome assembly construction

Sequencing data were basecalled using different versions of the Guppy/Dorado basecaller. Details on Guppy/Dorado versions and basecalling parameters used for individual sequencing runs are provided in (Supplementary Table S31)

First, we aimed to assemble the mycobiont (fungal partner) of the lichen symbiosis. To achieve this, we generated draft genome assemblies using either NECAT v0.0.1 (Chen et al., 2021a) or Hifiasm (v0.24.0 or v0.25.0) (Cheng et al., 2026). The parameters and input data used for each individual sample are provided in Supplementary Table S32.

Pore-C data were generated for nine samples representing a total of six different *Peltigera* species. The corresponding draft assemblies were used for the incorporation of the Pore-C data. The Pore-C analysis was performed using a customized pore-c pipeline, based on the ONT Pore-C workflow (https://github.com/epi2me-labs/wf-pore-cwas).

In brief, chimeric Pore-C reads were first bioinformatically digested using pore-c-py digest with the parameter --max_monomers 250. The resulting BAM file was converted to FASTQ format using samtools v1.20 fastq with the parameter -T “*”. The digested reads were then mapped to the corresponding draft assembly using minimap2 v2.28 with the parameters -ay -x lr:hq. Subsequently, contact links were annotated using pore-c-py annotate with the parameters --monomers --stdout --summary --paired_end --filter_pairs --paired_end_minimum_distance 100 --paired_end_maximum_distance 200. The resulting BAM file was sorted using samtools v1.20 sort with the parameters -m 2g -u - followed by name-based sorting using samtools v1.20 sort -n. The file was then preprocessed using rename_reads_for_bam.py, converted using samtools v1.20 view with the parameters -S -h -b -F 3340, and subsequently filtered with filter_bam.py. Genome scaffolding was performed using YaHS v1.2.1 (Zhou et al., 2023) with default parameters. Hi-C contact maps were generated using juicer pre v1.9.9, and visualization was carried out with Juicebox. Where necessary, assemblies were manually curated within Juicebox, and the reviewed assemblies were exported using juicer post v1.9.9.

The corresponding reference assemblies were then used to compare the draft-level assemblies of the remaining samples in order to identify chromosome-scale contigs. Matching contigs were extracted using seqtk subseq v1.4.

For samples in which chromosomes were fragmented in the draft assembly, we attempted to improve assembly contiguity by reducing the background of reads not originating from the mycobiont. To achieve this, matching contigs were first identified using D-GENIES and extracted for each sample that did not initially yield chromosome-scale assemblies. These contigs were used to generate a prefiltered assembly containing only contigs assigned to the mycobiont. This pre-filtered assembly was then used as a reference to map ONT Q15 reads and raw HiFi reads using minimap2 v2.28, with the parameters -ax map-ont for ONT reads and -ax map-hifi for HiFi reads. Subsequently, only reads that mapped to the filtered assembly were retained. Unmapped reads, secondary alignments, and supplementary alignments were removed using samtools view v1.20 with the flags -b -F 2308. The resulting BAM files were converted to FASTQ format using samtools bam2fq v1.20. The filtered reads were then used to generate new assemblies The resulting contigs were again compared to the reference Pore-C chromosome-scale assemblies using D-GENIES, and contigs corresponding to chromosomes were extracted using seqtk subseq v1.4.

Samples that did not reach chromosome-scale assemblies were retained at the draft level and subsequently filtered to retain only the mycobiont-derived contigs. For this purpose, one reference assembly per species was selected, and the contigs were aligned against it using minimap2 v2.30 with the parameter -x asm5.

Alignments corresponding to unmapped reads, secondary alignments, or supplementary alignments were removed, and only mappings with a minimum mapping quality of 30 were retained using samtools view v1.22 with the parameters -F 3332 -q 30. The contig which met these criteria were then extracted from the draft assembly using seqtk subseq v1.4.

To correct potential structural errors in the assemblies, Inspector v1.3 (Chen et al., 2021b) was used. For samples where HiFi reads were available, these were used for the correction; otherwise, ONT reads with a Q score of 15 were used. First, inspector was executed with the parameter --datatype ont or --datatype hifi, depending on the sequencing data. Subsequently, assemblies were corrected using the fixed version of inspector-correct.py (issue #36) with either --datatype nano-hq or --datatype pacbio-hifi, respectively.

To screen for potential contamination, the FCS-GX (Astashyn et al., 2024) pipeline was applied. First, run_gx was executed with the parameter --tax-id 48861. Subsequently, contaminants were removed using gx clean-genome with the parameters --action-report, --contam-fasta-out, and --min-seq-len.

Genome assembly statistics and completeness were evaluated using QUAST v5.2.0 (Gurevich et al., 2013) and Compleasm v0.2.7 (Huang and Li, 2023) with the ascomycota_odb12 dataset. Completeness and potential contamination were additionally assessed using EukCC v2.1.3 with the parameter --DNA and the database eukcc2_db_ver_1.1, K-mer–based completeness and assembly quality were assessed using Merqury v1.3 with k = 23, as well as CRAQ v1.0.9 using the -sms option to assess Regional Accuracy (R-AQI) and Structural Accuracy (S-AQI). For chromosome-scale assemblies, telomeric repeat sequences with the motif “TTAGGG” or its reverse complement “CCCTAA” were manually inspected at the beginning and end of each contig.

### Mycobiont genome annotation

Repetitive elements and transposable elements were identified using RepeatMasker v2.2.0. The genomes were soft-masked using the parameters -pa 250 -nolow -xsmall for subsequent gene prediction with BRAKER3, and hard-masked using the parameters -pa 250 -nolow -norna for use with Helixer. ONT cDNA data from *P. rufescens*_1 and *P. monticola_*2 were basecalled using Guppy/Dorado (Supplementary Table S31).The corresponding read statistics are provided in Supplementary Table S30. All cDNA reads from *P. rufescens* samples were combined into a single dataset and filtered for a minimum quality score of Q15. The same procedure was applied to all *P. monticola_*2 samples. Subsequently, the ONT cDNA sequences were mapped to the mycobiont genome assemblies using Minimap2 v2.30 with the -ax splice setting. For *P. monticola*_2 samples, the *P. monticola_*2 cDNA dataset was used, and for *P. rufescens*_1 samples, the *P. rufescens* 1 dataset was applied. For all other species, both the *P. monticola*_2 and *P. rufescens*_1 datasets were mapped to the respective genomes.

STRINGTIE2 version v3.0.0 (Kovaka et al., 2019) was used to assemble transcripts using the parameters -c 3 -j 3 -f 0.1. In addition, both HELIXER version 0.3.3 (Holst et al., 2025) with a fungi-specific model (fungi_v0.3_a_0100.h5), and BRAKER3 (Gabriel et al., 2024) version 3.0.8 with protein homology information from a UniProtKB/Swiss-Prot Ascomycota protein sequence set (status 2025_08_28) were used using the parameters --busco_lineage=ascomycota_odb10, --filterOutShort, --gff3 and --fungus were used to predict genes *ab-initio*.

To integrate gene predictions from the three approaches, Mikado v2.3.3 was used. Prior to integration, splice junctions were validated using Portcullis v1.2.4 to retain only high-confidence junctions. Read alignments flagged as unreliable by Portcullis were removed, and the resulting set of validated junctions was supplied to Mikado. For transcript scoring, the configuration file scerevisiae.yaml was applied. During the scoring procedure, transcript models generated by StringTie were assigned a weight of 2, whereas predictions from Helixer and BRAKER were each assigned a weight of 1. The resulting annotation file was then filtered for the longest isoform using Agat v1.4.0 script agat_sp_keep_longest_isoform.pl. Proteome completeness was assessed using Compleasm prot v0.2.7 (Huang and Li, 2023) with the ascomycota_odb12 dataset.

Functional annotation was performed using InterProScan v5.73-104.0 with the options -f tsv, -goterms, -iprlookup, and -pa. In addition, EggNOG-mapper was applied using the eggnog7_annotator.sh script with default parameters. For similarity-based annotation, DIAMOND v2.1.9 was used to construct protein databases and perform BLASTP searches against the UniProt Swiss-Prot, *Saccharomyces cerevisiae* and *Aspergillus nidulans* protein datasets. Searches were conducted using the --more-sensitive mode with an e-value threshold of 1e-5. Up to five target sequences were retained for Swiss-Prot searches, whereas a maximum of one target sequence was retained for *Saccharomyces cerevisiae* and *Aspergillus nidulans* searches. KEGG Orthology assignments were generated using KofamScan v1.3.0 with the exec_annotation workflow and default profile databases. Finally, all annotation results were integrated and merged using FungalMercator.

To assess the transposable element (TE) landscape of the genus *Peltigera*, TE detection was performed on the mycobiont genome assemblies using EDTA v2.2.2. Initially, LTRs, TIRs, and Helitrons were detected separately, followed by the final EDTA annotation step. Subsequently, a pan-EDTA library was constructed from the EDTA outputs using the panEDTA.sh script included in EDTA v2.2.2.

### SPALN-Based Recovery and Validation of Orthogroup Members

Orthogroups were inferred using OrthoFinder v3.1.3 with one representative genome per species and default settings. Gene copies not identified by OrthoFinder were subsequently recovered by aligning orthogroup representative proteins against genome assemblies using SPALN. To reduce false-positive predictions, recovered loci were evaluated using five independent validation criteria. First, protein similarity was assessed by DIAMOND searches against original orthogroup members, requiring a bitscore above an orthogroup-specific threshold. Second, splice-site quality was evaluated by requiring major GT–AG splice junctions in at least 50% of introns (or no introns). Third, predicted exon counts were required not to exceed the maximum observed among original orthogroup members. Fourth, local syntenic support was assessed based on the presence of orthogroup-associated genes within a ±10 gene neighborhood. Fifth, predicted proteins were required to initiate with methionine, lack internal stop codons, and fall within 0.5–2.0× the length of the orthogroup representative protein. Only loci passing all five criteria were retained as high-confidence recoveries for downstream analyses.

### Biosynthetic gene cluster analysis

Biosynthetic gene clusters (BGCs) were identified using antiSMASH v8.0.4. As the analysis focused on mycobiont genomes, assembled genome sequences were analysed together with their corresponding gene annotations, specifying the fungal taxon using the --taxon fungi option. To ensure that the existing gene annotations were retained, gene prediction was disabled by using the parameters --genefinding-gff3 and --genefinding-tool none.To enhance cluster detection and functional annotation, additional analysis modules were enabled, including --cassis for motif-based cluster boundary prediction, --clusterhmmer and --fullhmmer for domain-based annotation, --asf for active site identification, and --cb-knownclusters for comparison against the MIBiG database of characterized biosynthetic gene clusters. For downstream analyses, BGCs were summarized based on their predicted core biosynthetic type as reported by antiSMASH. Clusters were classified into PKS (T1PKS and T3PKS), NRPS (including NRPS-like), terpene (including terpene precursor clusters), hybrid clusters (containing multiple biosynthetic classes, indicated by comma-separated annotations), and other categories (e.g. indole or isocyanide). Each region was counted once, and hybrid clusters were not additionally assigned to individual categories. Clusters showing similarity to entries in the MIBiG database were recorded as known hits, with high-confidence matches defined according to antiSMASH confidence annotations.

To enable cross-sample comparisons, BGC identifiers were standardized by adding sample-specific prefixes prior to clustering. Sequence similarity relationships among BGCs were inferred using BiG-SCAPE (v2.0) with default parameters (including a similarity cutoff of 0.3). BiG-SCAPE constructs a sequence similarity network based on domain composition, sequence similarity, and gene cluster architecture, and partitions this network into Gene Cluster Families (GCFs), representing groups of homologous BGCs. For the presence-absence analysis GCFs were categorized based on their distribution across genomes as follows: core (present in all genomes), shared (present in all genomes of multiple species), species-specific (present in all genomes of a single species) and variable (all other patterns). For species represented by a single genome, singleton GCFs were classified as species-specific.

Biosynthetic classes were assigned to each GCF based on antiSMASH annotations of constituent BGCs. Cluster types were harmonized into the following categories: NRPS, PKS, terpene, and hybrid classes (NRPS–PKS, NRPS–terpene, PKS–terpene), as well as other for clusters lacking clear classification. For each GCF, class assignment was determined based on the combined domain architecture of all member BGCs.

### Structure-based putative GPCR identification and classification pipeline

Putative GPCR candidates were identified from the predicted proteomes of the 41 *Peltigera* genomes. To isolate proteins with the characteristic seven-transmembrane (7TM) architecture, we utilized TMHMM 2.0 (Krogh et al. 2001), a hidden Markov model-based algorithm that predicts transmembrane helices and differentiates between soluble and membrane proteins with high accuracy. Candidate sets were restricted to proteins with 6 to 8 predicted transmembrane helices, retaining likely GPCR candidates while accommodating minor uncertainties in sequence-based topology prediction. Because fungal GPCRs exhibit extreme primary sequence divergence that often defeats traditional sequence homology searches (Heinen et al., 2025), we employed a structure-guided classification approach. For each candidate passing the topological filter, a high-confidence, three-dimensional structural model was generated using AlphaFold2 (Jumper et al., 2021).

To prepare these models for robust structural comparison, their termini were trimmed inward from both the N- and C-termini until a residue with a predicted local distance difference test (pLDDT) score greater than 70 was reached. This step effectively removes low-confidence, unstructured regions that could introduce noise or false similarities during structural alignment. To classify the newly predicted *Peltigera* receptors, trimmed query structures were combined with 55 representative reference structures from the Liu et al. (Liu et al., 2025) classification framework and searched against themselves using FoldSeek (van Kempen et al., 2024) (foldseek-linux-avx2 @ 24020d2) in exhaustive all-vs-all mode (--alignment-type 1, --exhaustive-search 1, -c 0.85, --cov-mode 5), outputting pairwise query and target TM-scores.

Using Foldseek’s TM-score output metric, the resulting pairwise structural alignments were converted into a distance matrix defined as 1 − TM-score. Finally, hierarchical agglomerative clustering (HAC) was performed on this distance matrix utilizing the SciPy library (Virtanen et al., 2020) in Python. The resulting dendrograms were cut at a default distance threshold to resolve each dataset into discrete structural clusters. To assign GPCR classes, a multi-threshold cascade resolution strategy was applied across five different thresholds (T = 0.9, 0.8, 0.7, 0.6, 0.5), evaluated in order from most to least stringent. Cluster identities were subsequently annotated by mapping the structural groupings to the previously defined Liu et al. reference representatives, allowing us to classify and infer the highly divergent *Peltigera* GPCR candidates based strictly on their conserved 7TM fold. Candidates that remained multi-class at all thresholds were retained as ambiguous and reported separately.

### Synteny analysis of mycobiont chromosomes

Synteny analyses of the mycobiont chromosomes were performed using two complementary approaches. Structural similarities between chromosomes were first assessed using ntSynt (v1.0.2) based on nucleotide-level genome alignments. Analyses were performed exclusively within species and not across species. ntSynt was executed with a divergence threshold of -d 5, while default parameters were retained for -k 24 and -w 1000. Results were visualized using ntsynt_viz.py.

Gene-based syntenic relationships were subsequently identified using JCVI (v1.5.6). This approach detects conserved syntenic regions across species based on orthologous genes, as well as within species through duplicated genes, by integrating gene order and sequence similarity. For interspecies comparisons, chromosomes were renamed according to the *P. rufescens_*1 T2T reference (Due-8) to ensure consistent chromosome nomenclature across species.

Annotation files were converted into a JCVI-compatible BED format using jcvi.formats.gff bed --type=mRNA --key=ID --primary_only. Orthologous anchors were then generated using jcvi.compara.catalog ortholog with the --no_strip_names option and subsequently simplified with jcvi.compara.synteny screen --simple. Karyotype plots were generated using jcvi.graphics.karyotype. To ensure correct orientation in the visualization, inverted chromosomes were marked with a “-” in the Seq-ID files.

### Phylogenetic tree reconstruction

103 Peltigeraceae metagenomes were obtained from public databases (97 short-read and six long-read; 100 *Peltigera,* two *Solorina,* and one *Pseudosolorina;* Supplementary Table S11). *Solorina* and *Pseudosolorina* metagenomes were used as outgroup taxa in our phylogenetic analysis (Miadlikowska et al., 2014; Widhelm et al., 2021; Zheng et al., 2025). Data was trimmed with fastp v0.23 (Chen et al., 2018) to trim reads with a 15-bp sliding window, removing windows in which fewer than 80 % of bases had a Phred quality ≥ Q18 or that contained at least five N bases. After trimming, reads shorter than 100 bp were discarded. The short-read metagenomes were assembled with SPAdes v4.2.0 (Nurk et al., 2017) and long-read metagenomes with Flye v2.9.6-b1802 (--meta option) (Kolmogorov et al., 2019). Short- and long-read contigs were then binned together with VAMB v5.0.4 (Nissen et al., 2021) using a co-binning approach. The resulting bins were the basis for recovering the mycobiont metagenome-assembled genomes (MAGs).

To identify candidate mycobiont MAGs, taxonomy to each contig in every bin was assigned using MMseqs2 v15.0.0 (Steinegger and Söding, 2017) against the UniRef90 database (downloaded 13 January 2026). Subsequently, hidden Markov models (HMMs) of genes found exclusively in the mycobiont, cyanobiont, or microbiome members of cyanolichens were built and used to further refine the identity of each bin (Eddy, 2011). This information was used to remove non-mycobiont contigs inside the proposed mycobiont MAGs. The completeness and contamination of each mycobiont MAG was checked with BUSCO (Simão et al., 2015). This yielded 103 mycobiont MAGs with > 85 % completeness and < 5 % contamination.

### Taxonomic identification of newly obtained *Peltigera* genomes

For phylogenetic placement of the new *Peltigera* MAGs, we profiled HMMs for six loci (ITS, LSU, *β-tubulin, RPB1, COR1B, COR16*) (Miadlikowska et al., 2014; Magain et al., 2018) to search the 41 *Peltigera* genomes newly sequenced in this study (Eddy, 2011; Garfias-Gallegos et al., 2025). To build the profile HMMs, alignments for these six loci were retrieved from the *Peltigera* reference using the T-BAS platform (Carbone and White, 2019; Chagnon et al., 2019). For this, 25 genomes spanning a range of fungal clades (Supplementary Table S33) were scanned to establish bitscore thresholds for distinguishing true positives from false positives. Custom scripts that integrated HMMER and SeqKit v2.13.0 (Shen et al., 2024) were used to identify and extract the nucleotide sequences of target loci whose hits exceeded the threshold from the 41 *Peltigera* genomes. Finally, the extracted loci were uploaded to the T-BAS online platform to phylogenetically place the 41 genomes within the *Peltigera* reference tree (Carbone et al., 2019).

To define a bitscore threshold for the HMMER searches, the 25-genome panel (Supplementary Table S34) was partitioned into four sections: positive, negative_close, negative_mid, and negative_far. All 25 genomes were searched with the six HMM profiles and the bitscore of the best hit per locus per genome was recorded. The identity of each extracted sequence was then verified using NCBI BLAST (Altschul et al., 1990). For the genomes in the positive section (five *Peltigera* MAGs assembled from public long-read data and curated in this study), the lowest bitscore value across the five genomes was taken as the positive_minimum value. Across all negative sections combined, the highest bitscore was used the noise_maximum. The bitscore threshold for each locus was then set as the midpoint between positive_minimum and noise_maximum (Supplementary Table S34).

### Phylogenetic analysis

To infer a species tree of the family *Peltigeraceae*, the phylogenomic pipeline described in previous studies (Garfias-Gallegos et al., 2025) was followed. In brief 2,826 BUSCO v5.8.2 (Simão et al., 2015) genes using the ascomycota_odb12 (Tegenfeldt et al., 2025) were extracted from the 144 genomes (Supplementary Table S11). Protein sequences were aligned with MAFFT v7.525 (Katoh et al., 2002) and back-translated to codon alignments with PAL2NAL v14 (Suyama et al., 2006). The resulting alignments were trimmed with trimAl v1.5.1 (Capella-Gutiérrez et al., 2009), retaining only sites present in at least 95 % of taxa. Only genes represented in at least 95 % of the genomes in the dataset were retained, resulting in 2,561 genes. The obtained filtered genes were used to infer maximum likelihood gene trees with IQ-TREE v3.1.1 (Wong et al., 2026). The resulting gene trees were then used as input to infer a species tree with weighted-ASTRAL (Zhang et al., 2025). The species tree was visualized with the R package ggtree v.3.4.4 (Yu et al., 2017).

### Photobiont genome assembly and annotation

#### Cyanobacterial photobiont

To assemble the second major component of the lichen symbiosis, we aimed to reconstruct the cyanobacterial photobiont. As the cyanobacterial partner of *Peltigera* belongs to the genus *Nostoc*, an estimated genome size of 6–8 Mb was expected. Therefore, all previously generated assemblies were screened for circular contigs within this size range. The corresponding contigs were extracted using seqtk subseq version 1.4 and compared to the *P. membranacea* cyanobiont reference genome (GCA_002949735.1) using D-GENIES (Cabanettes and Klopp, 2018). The assembly strategies from which the photobionts were recovered are listed in Supplementary Table S35.

Genome statistics and completeness were assessed using QUAST v5.2.0 (Gurevich et al., 2013) and Compleasm v0.2.7 (Huang and Li, 2023) with the nostocales_odb12 model (Supplementary Table S8). For comparative purposes, the genomes were restarted at the *dnaA* gene region using SeqKit v2.9 with the restart parameter. Prior to this, the *dnaA* gene region was identified through genome annotation using Prokka v1.14.5 (Seemann, 2014) with default settings. After successful restarting of each genome, the annotation step was repeated. Proteome completeness was additionally assessed using Compleasm protein v0.2.7 with the nostocales_odb12 model. Taxonomic identities of the cyanobionts were inferred from *rbcLX* loci extracted from the assembled genomes. The extracted sequences were classified using the T-BAS phylogenomic framework. Assignments followed the hierarchical classification scheme implemented in T-BAS, comprising sections, species complexes, and phylogroups. Because genomic boundaries within some *Nostoc* lineages remain unresolved, certain strains could not be confidently assigned to a phylogroup and were instead classified at the species complex level. For each genome, the finest confidently supported taxonomic designation was retained and reported as the corresponding *Nostoc* operational taxonomic unit (OTU).

#### Algal photobiont

As *P. aphthosa_5* (Paph1, Paph2) and *P. leucophlebia_*2 (Pleu1) are tripartite lichen and harbor a green alga of the genus *Coccomyxa* as their primary photobiont, we additionally assembled the corresponding algal genome. For this purpose, the primary draft assembly was mapped against the *Coccomyxa* sp. Obi reference genome (GCA_020887355.1) as well as the *Coccomyxa viridis* SAG 216-4 genome (Kraege et al., 2025) using D-Genies (Cabanettes and Klopp, 2018). Matching contigs were extracted using seqtk version 1.4 with the subseq parameter. Genome statistics were calculated with QUAST version 5.2.0 (Gurevich et al., 2013), and completeness was assessed using Compleasm run version 0.2.7 (Huang and Li, 2023) with the chlorophyta_odb12 model (Supplementary Table S10). For taxonomic classification, the ITS loci from all *Coccomyxa* genomes were extracted and examined. Based on this result, we additionally compared the genomes with the *Coccomyxa elongata* SAG 216-3b reference genome (Kraege et al., 2025), which represents a more appropriate phylogenetic comparator than the genomes used in our initial analyses to validate the previous selection of algal contigs. To screen for potential contamination, the decontamination pipeline fcs-gx was applied using the parameter --tax-id 41891. Detected contaminant contigs were subsequently filtered out from the assembly. To assess assembly completeness, all contigs of the *P. aphthosa*-associated algal photobiont genomes were manually inspected for the presence of a telomeric repeat motif (CCCTAAA/TTTAGGG) at both ends.

### Identification of Lectin candidates

Lectin candidates were identified from the InterProScan annotations of 41 Peltigera mycobiont genomes. First, all InterPro accessions associated with lectins were collected by querying the InterPro database for entries containing the term “lectin” in the domain name or description. This search yielded 619 lectin-related accessions from InterPro and its member databases (Supplementary Table S36). These accessions were then searched against the InterProScan results of all genomes. When multiple lectin-associated accessions matched the same gene id, only the lectin-related hit with the lowest e-value was retained. The resulting candidate set was subsequently filtered using a stringent e-value threshold of ≤ 1 × 10⁻^10.^ To reduce false-positive assignments, the strongest overall InterProScan hit for each candidate was subsequently evaluated. Candidates whose best-supported annotation corresponded to non-lectin functions were removed, whereas ambiguous cases were further assessed using FungalMercator functional annotations, which integrate evidence from multiple annotation resources.

Remaining unresolved candidates were manually curated based on their combined InterProScan and FungalMercator annotations. Proteins containing LCCL and PAN1/GLEYA domains were also retained because both domain types have been associated with carbohydrate-recognition functions (Supplementary Table S36).

### Dissection of thallus layers and layer-specific RNA sequencing

To validate the expression patterns of the target gene families (GPCRs, AMPs, and F-type lectins) across different thallus regions, fresh lichen thallus material was collected from the UB1 sampling site at the same location as the original specimen. Thalli were cleaned of debris as described above. To define thallus layers laser-based microscopy was performed on a Zeiss Axio Observer.Z1 equipped with a Prime BSI Express sCMOS camera (Visitron Systems, Puchheim, Germany). Excitation was provided by a VS-LMS4 Laser-Merge-System (Visitron Systems, Puchheim, Germany) using a solid-state laser targeting GFP (488 nm, 50 or 100 mW), and emission was detected using a 520 nm filter. Image acquisition was performed using VisiView software (Visitron Systems, Puchheim, Germany). Due to their thinness and difficulty of separation, the cortex and photobiont layer were combined into a single layer (CP), while the underlying medulla (M) and rhizines (R) were defined as separate layers.

For each layer, at least 20 mg of material was collected in three replicates using a binocular microscope, fine tweezers, and scalpels. Dissected samples were immediately shock-frozen in liquid nitrogen and stored at −80°C until further processing. RNA extraction and sequencing library preparation were performed as described in Section Oxford Nanopore cDNA library construction and Sequencing using the SQK-PCS114 Kit (Oxford Nanopore Technologies, UK). Libraries were sequenced on a FLO-MIN114 flow cell using the MinION Mk1B device for 100h.

#### Differential gene expression analysis

Transcript-level count data were obtained from Oarfish (v0.9.0) quantification outputs for nine samples representing three lichen tissue layers: Cortex/photobiont layer (CP), Medulla (M), and rhizines (R) with three replicates per condition. Count matrices were imported into R (v4.5.3) and processed using the *edgeR* package (v4.8.2). Lowly expressed transcripts were removed using filterByExpr with default parameters, retaining transcripts exceeding an internally determined CPM threshold (approximately 10 counts in the smallest library) in at least one experimental group. Library size differences were normalized using the trimmed mean of M-values (TMM) method. Normalized expression values were transformed to log2 counts per million (logCPM) for downstream visualization and summary statistics. Principal component analysis (PCA) and multidimensional scaling (MDS) were performed to assess sample clustering and overall expression variance structure. Differential expression was assessed using a quasi-likelihood negative binomial generalized linear model (GLM) framework. Quasi-likelihood models were fitted using glmQLFit with default settings, and differential expression was tested using glmQLFTest using pairwise contrasts. Genes with an Benjamini Hochberg false discovery rate < 0.05 were considered differentially expressed.

#### Heatmap visualization

Differentially expressed genes (DEGs) were visualized using heatmaps generated with the pheatmap package (v1.0.13). Expression matrices were derived from log2 counts per million (logCPM) values and aggregated to per-group averages. Relative expression patterns were represented by row-wise mean-centering.

#### InterPro domain enrichment analysis

InterPro domain enrichment was assessed using one-sided Fisher’s exact tests comparing DEG sets to the full gene universe. A binary gene–domain matrix was constructed from IPR annotations. DEGs were further subdivided into up- and downregulated sets based on log fold. Domains represented by fewer than three differentially expressed transcripts were excluded from testing to avoid unstable enrichment estimates. Resulting p-values were adjusted for multiple testing using the Benjamini-Hochberg method, and domains with an adjusted FDR < 0.05 were considered significantly enriched. Fold enrichment was calculated as the ratio of domain frequency among differentially expressed transcripts to its frequency in the background universe. To reduce redundancy, closely related IPR annotations describing the same functional feature at different levels of specificity (such as domain, superfamily, and conserved site entries belonging to the same protein family) were manually consolidated into representative domain categories, retaining the entry with the lowest FDR as the group representative. Enrichment results were visualized as bubble plots using ggplot2 (v4.0.2).

### Cell wall composition analysis

Intact thalli of *Peltigera rufescens_1* were manually cut into cross sections, and the photobiont layer was separated from the medulla layer under a Leica S6D Greenough stereo microscope at 20x magnification. From each sample, three replicates per layer were collected. The material was freeze-dried overnight in a Coolsafe™ system (Scanvac) and subsequently milled into a powder with 2x 5 mm steel balls for 2x5 min in a MM400 mixer mill (Retsch). Alcohol-insoluble residue (AIR) was prepared following the protocol of (Foster et al. 2010) with the modifications described below. The powdered tissue was resuspended in 70 % ethanol (1 mL) and milled again for 5 min. After centrifugation for 10 min at 14000 rpm, the ethanol was replaced with chloroform methanol (1:1 (v:v)), followed by another cycle of milling and centrifugation. Acetone was then added, again followed by milling and centrifugation. The pellet was resuspended in 200 µL acetone and transferred to a pre-weighed tube before the acetone was evaporated under constant airflow.

AIR material (0.5 mg) was hydrolyzed with 2 M TFA at 121°C for 1.5 h, according to (Foster et al., 2010). The resulting monosaccharide mixtures were then separated on a HPAEC system (Knauer Azura) according to (Wang et al., 2023). Ribose was used as an internal standard. The results are depicted as the weight percentage (wt%) of each monosaccharide of the total carbohydrate fraction.

### Fluorescent fucose glycoconjugate labelling of *Peltigera* section

A thallus from *Peltigera neckeri* was manually sectioned and incubated in 200 µL phosphate-buffered saline containing 2 µg α-L-Fucose-C3-PAA-fluorescein (Sigma Aldrich Cat. Nr. 0027-FP). The mixture was incubated for 1 h at RT, afterwards the liquid was replaced with 200 µL PBS and the thallus subsequently imaged with a Leica DM2000 microscope equipped with a CoolLED pE-300 white illumination system and imaged with a Leica MC170HD camera. Fluorescein fluorescence was imaged with an L5 filter (bandpass 480/40 nm) and autofluorescence imaged with a Texas Red ET filter set.

### Prediction of secreted proteins and functional annotation

Signal peptides were predicted for annotated proteins from both mycobiont and photobiont genomes using SignalP 6.0 (Teufel et al., 2021). Proteins predicted to be secreted were collectively defined as the secretome. Carbohydrate-active enzymes (CAZymes) were annotated using dbCAN v3 (Zheng et al., 2023). Proteins containing transmembrane helices were identified using TMHMM 2.0 (Krogh et al., 2001). Pfam functional annotation was performed using HMMER 3.3 (Eddy et al., 2011).

### Structure prediction and antimicrobial protein clustering

To assess the antimicrobial potential of the secretome, protein structures were predicted using ESMFold v1.0.3 (Lin et al., 2023). CAZymes, transmembrane-containing proteins and proteins larger than 800 amino acids were excluded from the analysis. The predicted structures were subsequently analysed using AMAPEC v1.0b (Mesny et al., 2026) to identify candidate antimicrobial proteins (AMPs). Orthologous relationships among predicted AMPs were inferred using OrthoFinder, grouping proteins into orthogroups based on sequence similarity. To further assess structural similarity beyond sequence-based clustering, Foldseek v9.427df8a (van Kempen et al., 2021) was applied using the easy-cluster workflow with a TM-align threshold of ≥ 0.5.

### Analysis of “two-speed genome” traits

The panEDTA outputs, excluding rRNA and tRNA, were defined as TEs/repeats for the two-speed genome analysis. The distances of genes to flanking TEs were examined using ‘closest’ command of bedtools v2.31.1(Quinlan, 2014) with the options ‘-D ref’ and either ‘-id’ or ‘-iu’, by comparing two BED files containing gene or TE annotations. The distances between flanking genes were examined using bedtools ‘closest’ with the options ‘-D ref’, by comparing two BED files containing gene annotations or annotations of intergenic regions identified using bedtools ‘subtract’ with BED files of genomes and genes. Effect sizes were quantified as the standardized mean difference, Hedges’ g (Hedges, 1981) For each individual sample, *P*-values were calculated from Z-scores and subsequently corrected for multiple testing using the false discovery rate (FDR) method (Benjamini and Hochberg, 1995) within each flanking direction. To obtain a consolidated estimate of the effect size across the *Peltigera* genus while accounting for the evolutionary relationships among sampled taxa, we performed a phylogenetic multi-level meta-analysis using the rma.mv function in the metafor package (Viechtbauer, 2010). A rooted phylogenetic tree of the *Peltigera* samples was used to construct a phylogenetic variance-covariance matrix under a Brownian motion model of trait evolution. To account for both phylogenetic signals and taxon-specific variation not captured by the phylogeny, we fitted a multilevel random-effects model (Nakagawa and Santos, 2012) including a random effect linked to the phylogenetic covariance matrix and an independent random effect for each sample. Summary effect sizes and 95% confidence intervals were estimated using restricted maximum likelihood (Harville, 1977). The statistical significance of the meta-analytic summary effect was determined after FDR correction. The statistical analyses were performed in R version 4.5.3, and plots were visualized using the R package ggplot2 v4.0.2 (Wickham, 2016).

### Identification of *Starships* and captain genes

*Starship*-like elements and captain-like YR genes were identified using starfish v1.1.1 (Gluck-Thaler and Vogan, 2024), as described previously. *Starship*/YR navis and family classifications were assigned according to definitions in previous studies (Gluck-Thaler and Vogan, 2024; Urquhart et al., 2024). Genomic synteny was identified by genome alignment using ‘nucmer’ command of MUMmer v4.0.1 (Marçais et al., 2018) with the options ‘--maxmatch -L 1000’, followed by filtering of hits based on nucleotide identity using ‘delta-filter’ command with the options ‘-1 -i 80’. Genomic regions containing *Starship*-like elements were visualized using the R package gggenomes v1.1.3 (Hackl et al., 2024). The phylogeny of the *Peltigera* genomes was visualized with the R package ggtree v4.0.5 (Yu et al., 2017). Genes located within 50 kb downstream of the captain-like YR genes were identified using the bedtools ‘window’ function by comparing a BED file of the YR genes with a BED file of all genes, using the options ‘-sw -l 0 -r 50000’. Genes orthologous to proteins in PHI-base v4.17 (Urban et al., 2020) were identified using OrthoFinder v3.1.0 (Emms and Kelly, 2019).

## Supporting information

All Supplementary Tables are included in a single file

## Data availability

HiFi, Pore-C and ONT sequencing data has been uploaded to the European NucleotideArchive (ENA) under project code: PRJEBXX. The genome assemblies and gene annotations can be found at the ENA under accessions XX to XX. Data including analyses is also available at https://doi.org/XXXX hosted at the PLANTdataHub (Weil et al., 2023).

## Acknowledgments

This study was supported by Deutsche Forschungsgemeinschaft within the framework of CRC1535 to BU, FA, MF, MP, BT project no. 458090666. MF, FA, MP, BT, and BU also thank CEPLAs EXC 2048/2 project no. 390686111. SW was supported by DFG grants nr. 405889398 and 531156553. The data and results provided by CPDH, DGG, FL, and JM for this study resulted from funding by the National Science Foundation (BEE 1929994). We thank Nele Grünig, Peter Braubach, Julius Trebbien and Bernd Gliwa for providing samples. VJ-R also acknowledges Pilzforum.eu for enabling communication and public outreach.

## Supplementary Figures

**Supplementary Figure 1:**
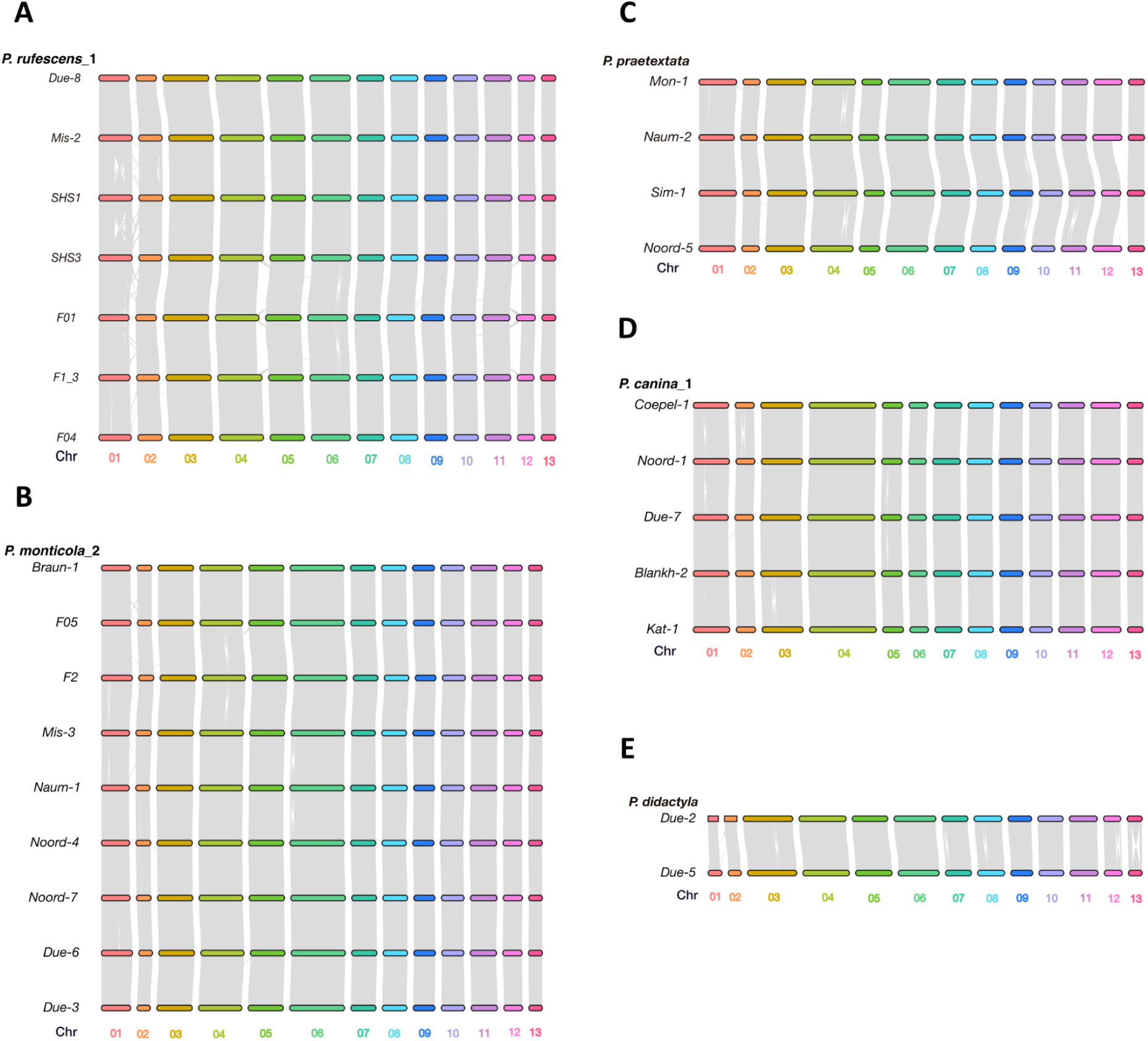
Gene-based synteny within *Peltigera* species. Synteny based on gene-coding regions was inferred using the jcvi toolkit. Chromosomes are represented as linear tracks, and connections indicate collinear gene blocks between homologous regions. (A) Intraspecific synteny in *Peltigera rufescens*, (B) *Peltigera monticola*, (C) *Peltigera praetextata*, (D) *Peltigera canina* and (E) *Peltigera didactyla*. These comparisons highlight the conservation of gene order across chromosome-scale assemblies within a species.

**Supplementary Figure 2:**
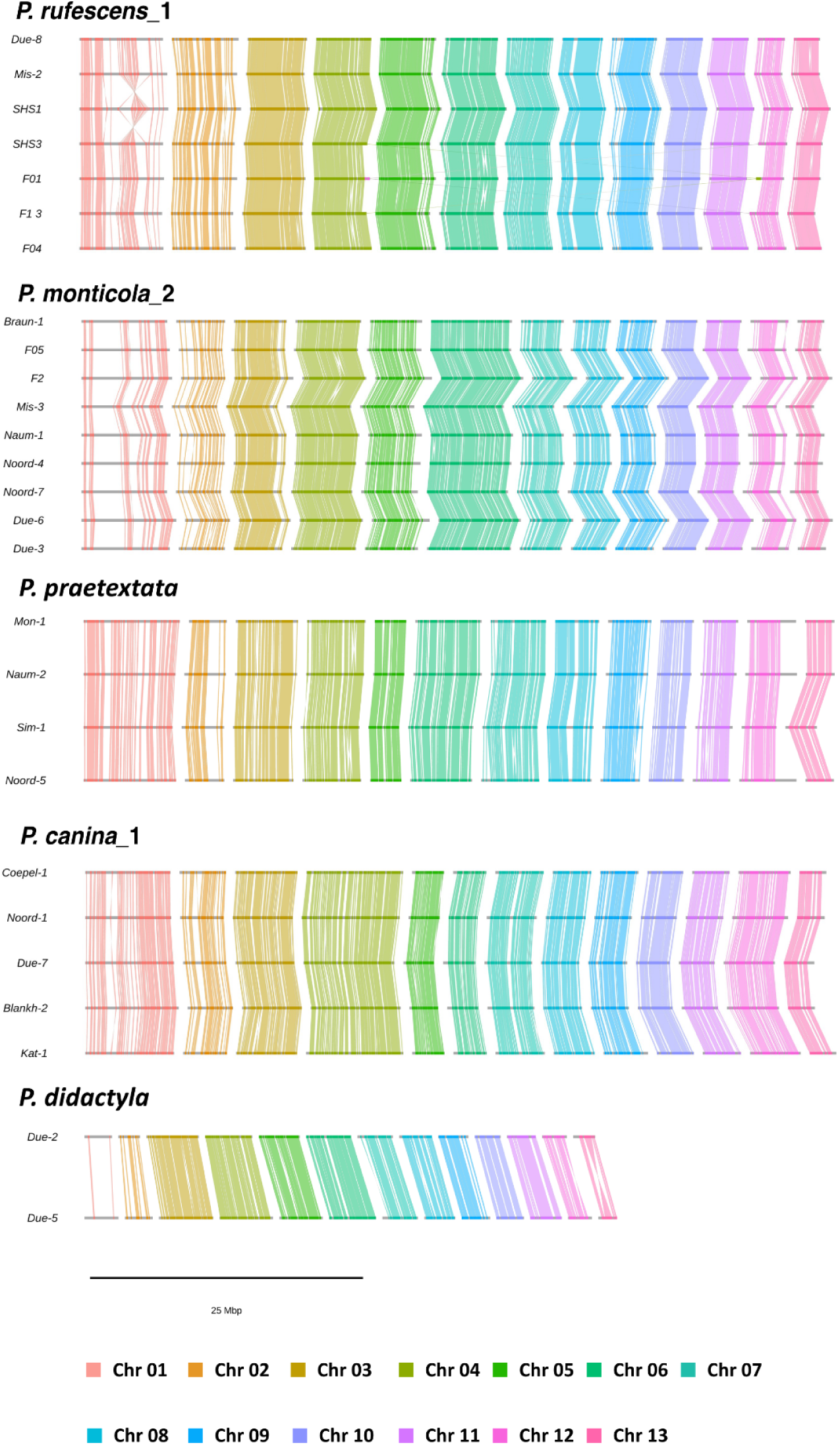
Sequence-based synteny across *Peltigera* species. Sequence-based synteny was inferred between *Peltigera* species for which at least two chromosome-scale genome assemblies are available, using ntsynt. Chromosomes are shown as linear representations, and colored ribbons connect syntenic regions between genomes. These ribbons highlight conserved genomic segments and reveal structural rearrangements such as translocations and inversions across chromosomes.

**Supplementary Figure 3.**
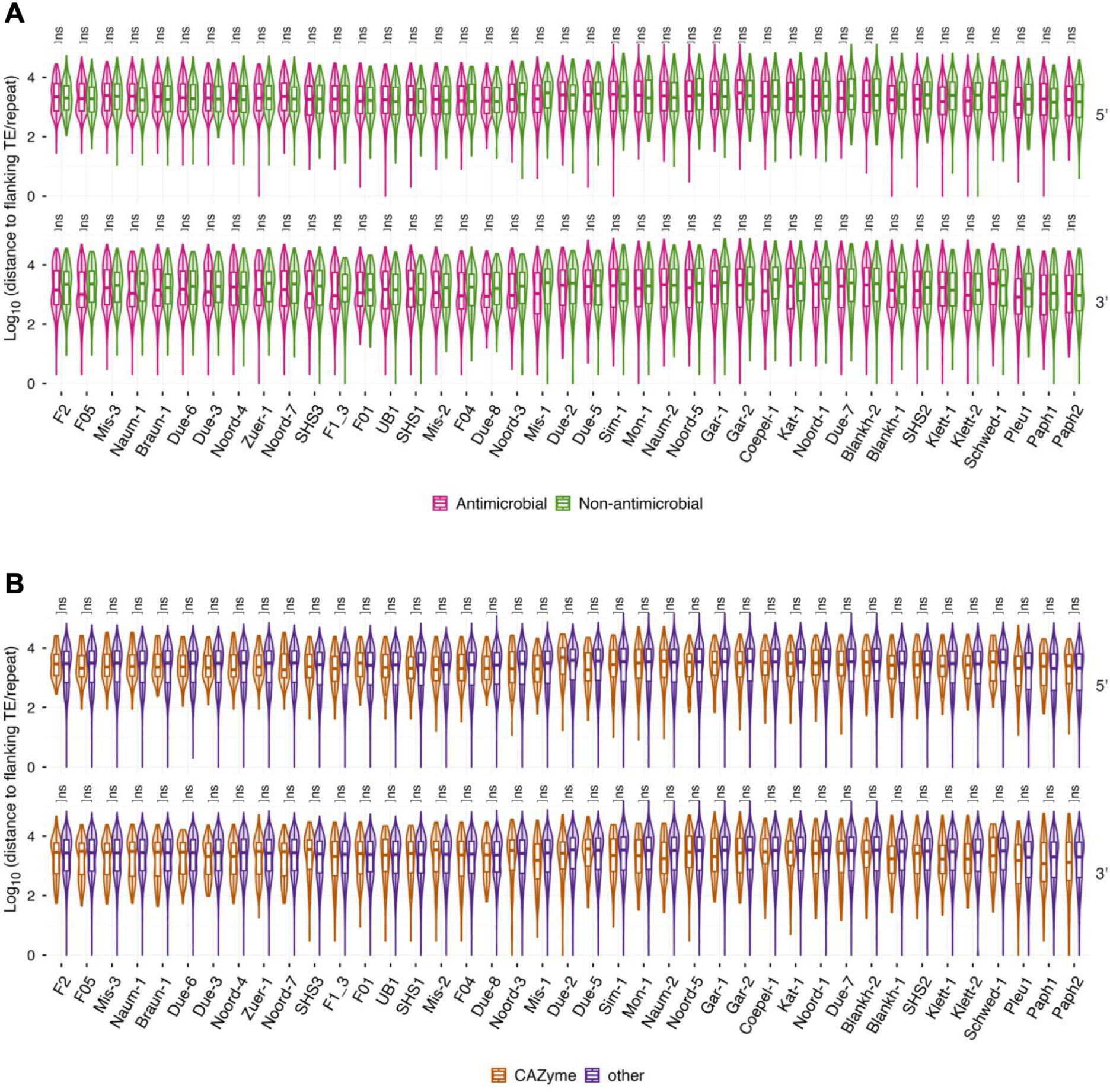
Distances to transposable elements (TEs) do not differ significantly between antimicrobial/CAZyme genes and other secreted protein (SP) genes. Violin and box plots depict log-transformed distances from the 5’ and 3’ ends, respectively, of antimicrobial genes and other SP genes (A), and of carbohydrate-active enzyme (CAZyme) genes and other SP genes (B), to the nearest flanking TE/repeat, with gene categories shown in different colors. In the box plots, the center, upper, and lower lines denote the median, third quartile, and first quartile, respectively. Asterisks indicate significant differences based on Wilcoxon rank-sum tests with Benjamini–Hochberg correction for multiple testing (**** *p* < 0.0001, *** *p* < 0.001, ** *p* < 0.01, * *p* < 0.05, ns: not significant).

**Supplementary Figure 4.**
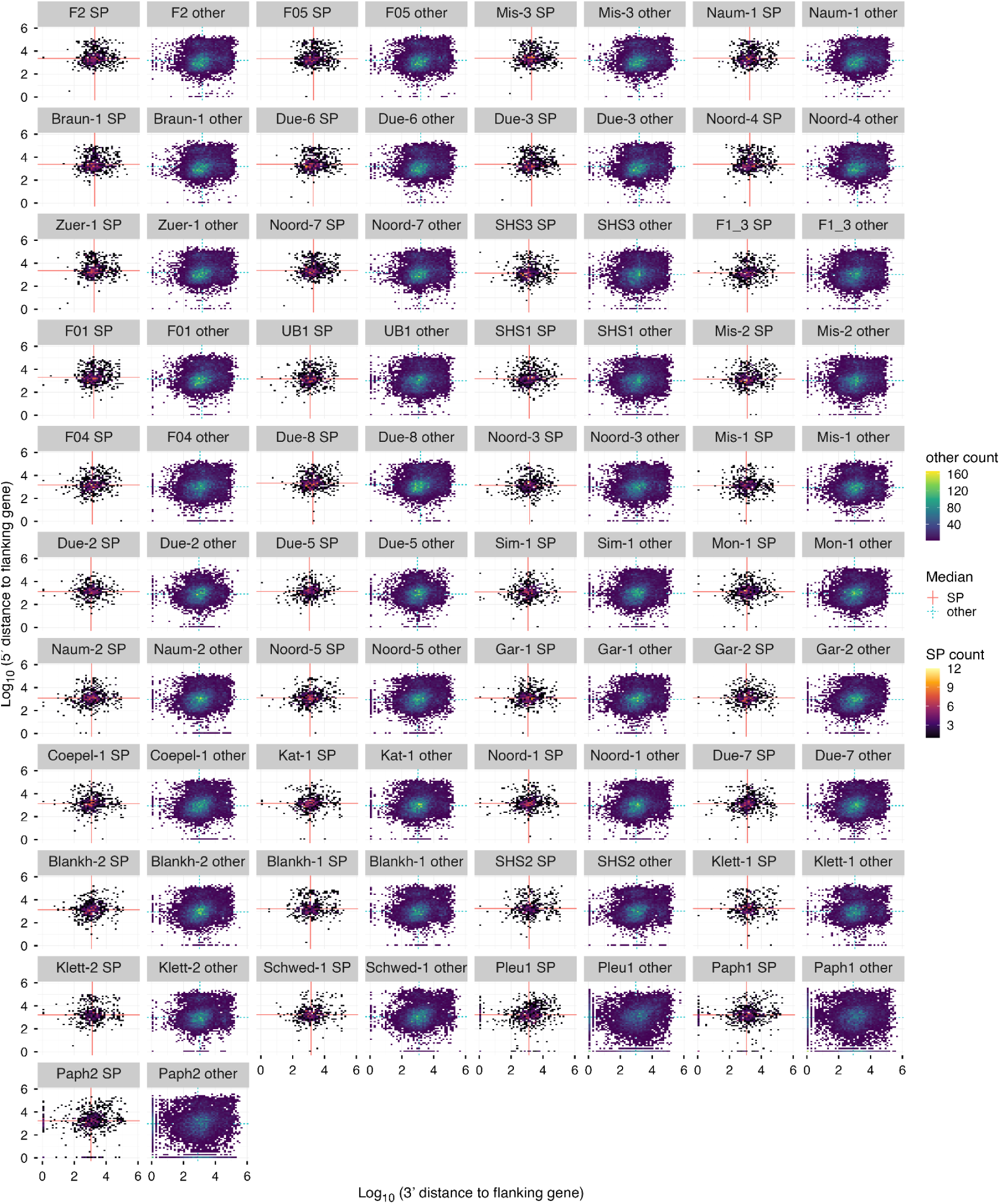
Secreted protein (SP) genes tend to localize in gene-sparse regions across the 41 *Peltigera* genomes. The heatmaps show the counts of SP genes and other protein-coding genes in separate panels, with the X- and Y-axes each divided into 50 bins. The X- and Y-axes represent the distances from the 5’ and 3’ ends of each gene, respectively, to the nearest flanking protein-coding gene. Thus, the lower-left region indicates gene-rich regions, whereas the upper-right region indicates gene-sparse regions. Red solid and blue dashed lines indicate the median values for SP genes and other protein-coding genes, respectively, in each *Peltigera* sample. The statistical differences are shown in Supplementary Figure 5.

**Supplementary Figure 5.**
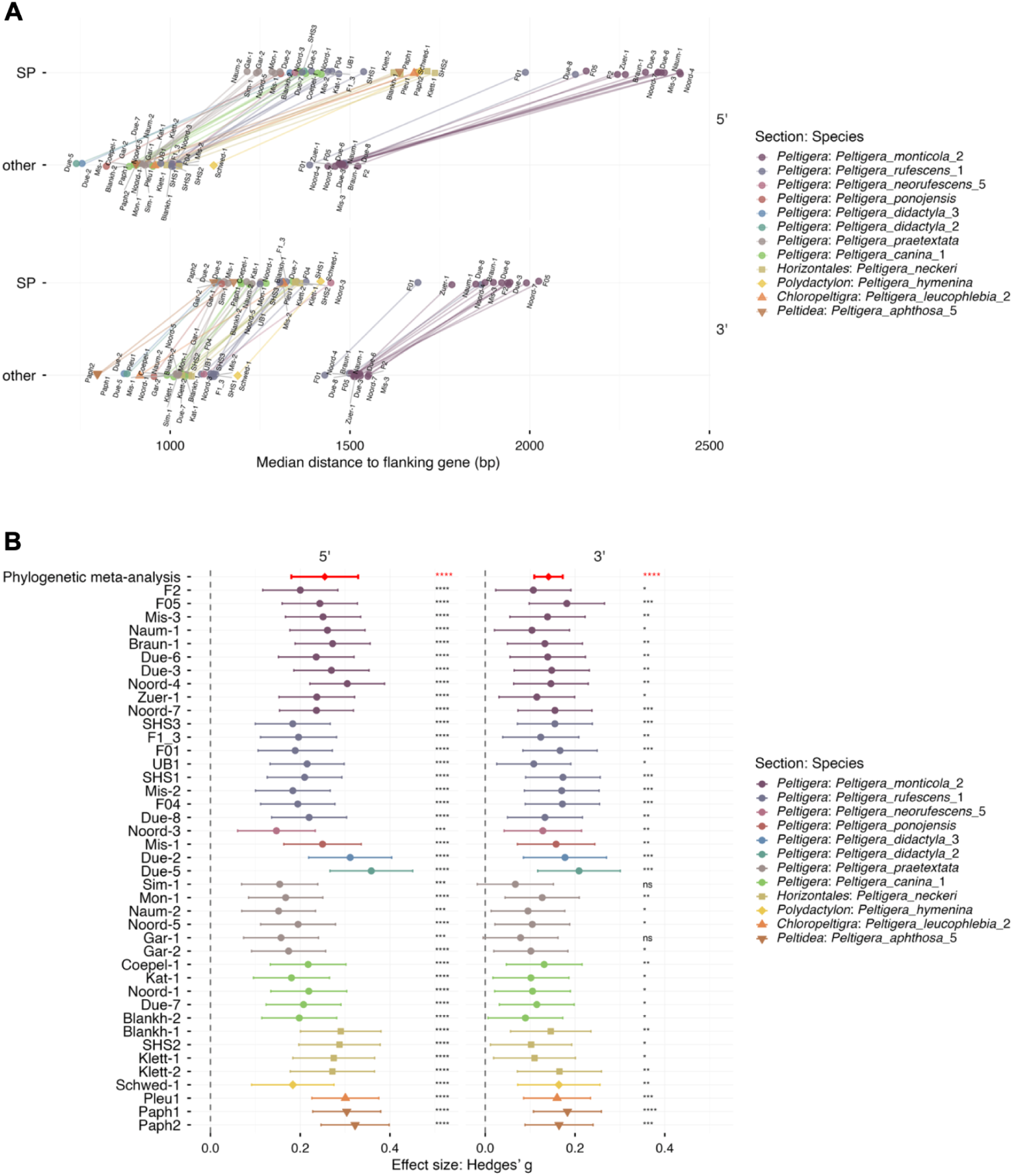
Secreted protein (SP) genes are more distant from flanking protein-coding genes than other protein-coding genes across *Peltigera* spp. (A) Scatter plot of the median distance to the nearest protein-coding genes for SP genes and other protein-encoding genes, shown separately for each sample and for the 5’ or 3’ side. Points represent median values, and lines connect the median values of SP genes and other protein-encoding genes within each *Peltigera* sample. (B) Forest plots illustrate the standardized mean differences (Hedges’ g) in intergenic distances (normalized by log_10_ (length + 1)) between SP and other protein-coding genes for each sample and the flanking direction (5’ and 3’). Each point and horizontal line represent the estimated effect size and its 95% confidence interval for individual samples. The red diamonds and bold red error bars indicate the summary effect sizes and 95% confidence intervals derived from a phylogenetic multi-level meta-analysis, which accounts for both phylogenetic relatedness and species-specific random effects. Statistical significance was determined using false discovery rate-corrected p-values (*: *P*<0.05, **: *P*<0.01, ***: *P*<0.001, ****: *P*<0.0001; ns: not significant). Positive effect sizes indicate longer distances for SP genes compared to other protein-coding genes.

**Supplementary Figure 6:**
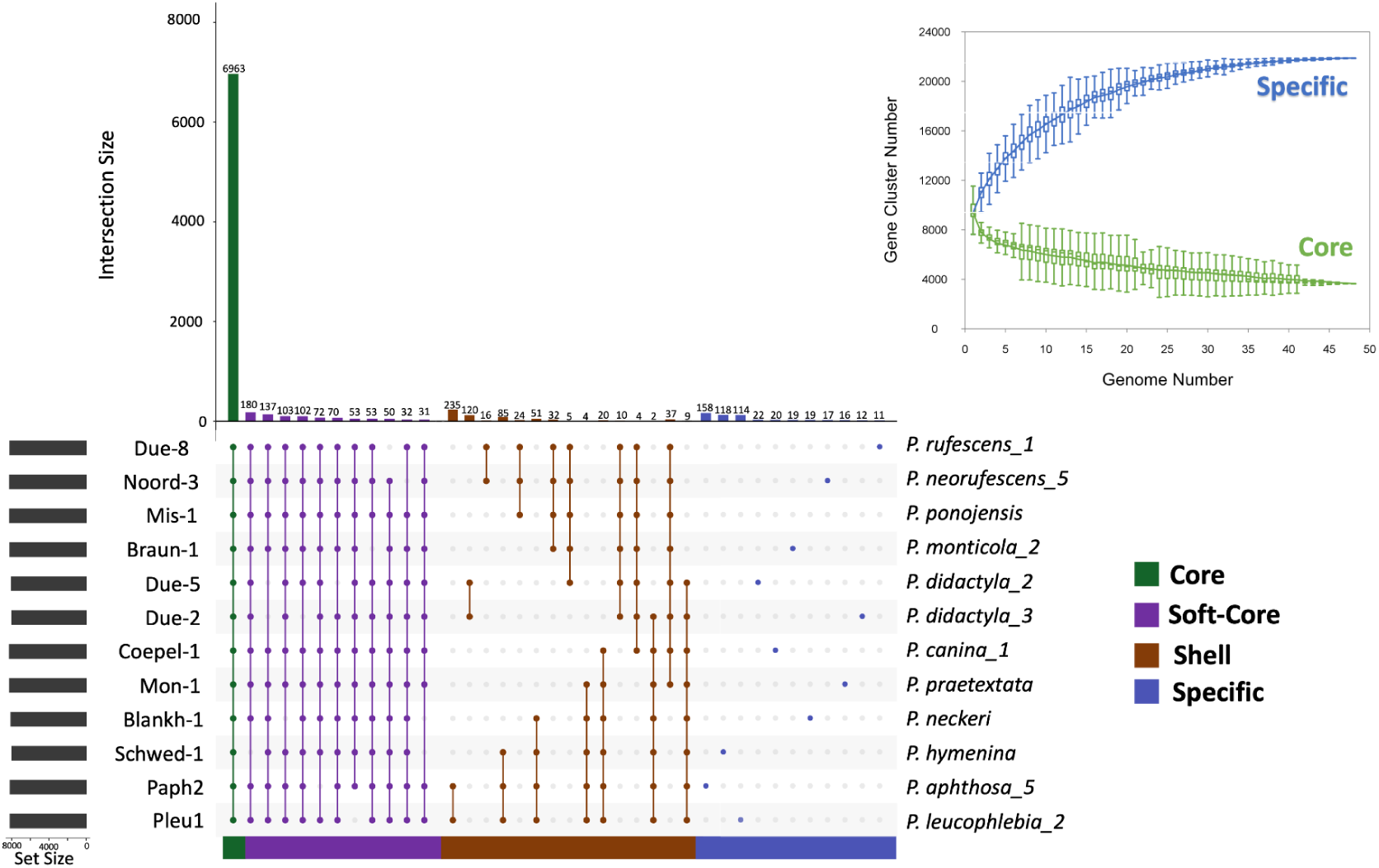
Analysis of orthologous group composition across the *Peltigera* super-pangenome. UpSet plot of orthologous group distributions across the analyzed *Peltigera* species. Core orthogroups were defined as present in all representative species, soft-core orthogroups as present in 10–11 species, shell orthogroups as present in 2–9 species, and species-specific orthogroups as present in only a single species. The pangenome analysis was constructed using all genomes generated in this study together with publicly available *Peltigera* genomes. The insert (upper right corner) shows the number of core genes (green) and specific genes (blue) when adding new genomes across all species.

**Supplementary Figure 7:**
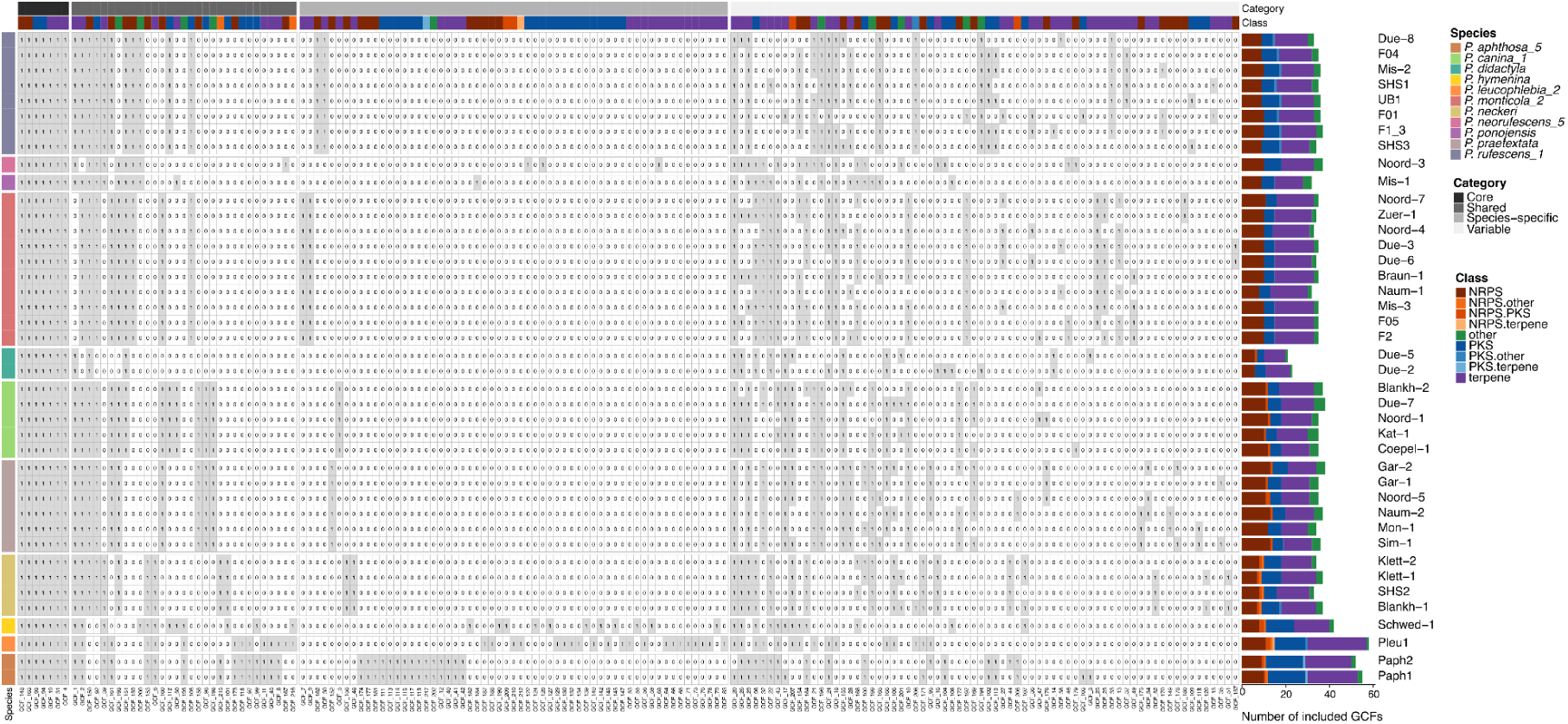
**Biosynthetic gene cluster family diversity across *Peltigera*** Presence–absence matrix of biosynthetic gene cluster families (GCFs) inferred using BiG-SCAPE. Rows represent genomes and columns correspond to GCFs arranged by distribution pattern. GCFs are classified as core, shared, species-specific or variable. Filled cells indicate presence. Genome labels are colored by species, and GCFs are annotated by biosynthetic class based on antiSMASH domain architecture. b) Stacked bar plots showing the class composition of the same GCF set across genomes. Bars indicate the number of GCFs per genome, partitioned by biosynthetic class, with genome order matching panel (a).

**Supplementary Figure S8.**
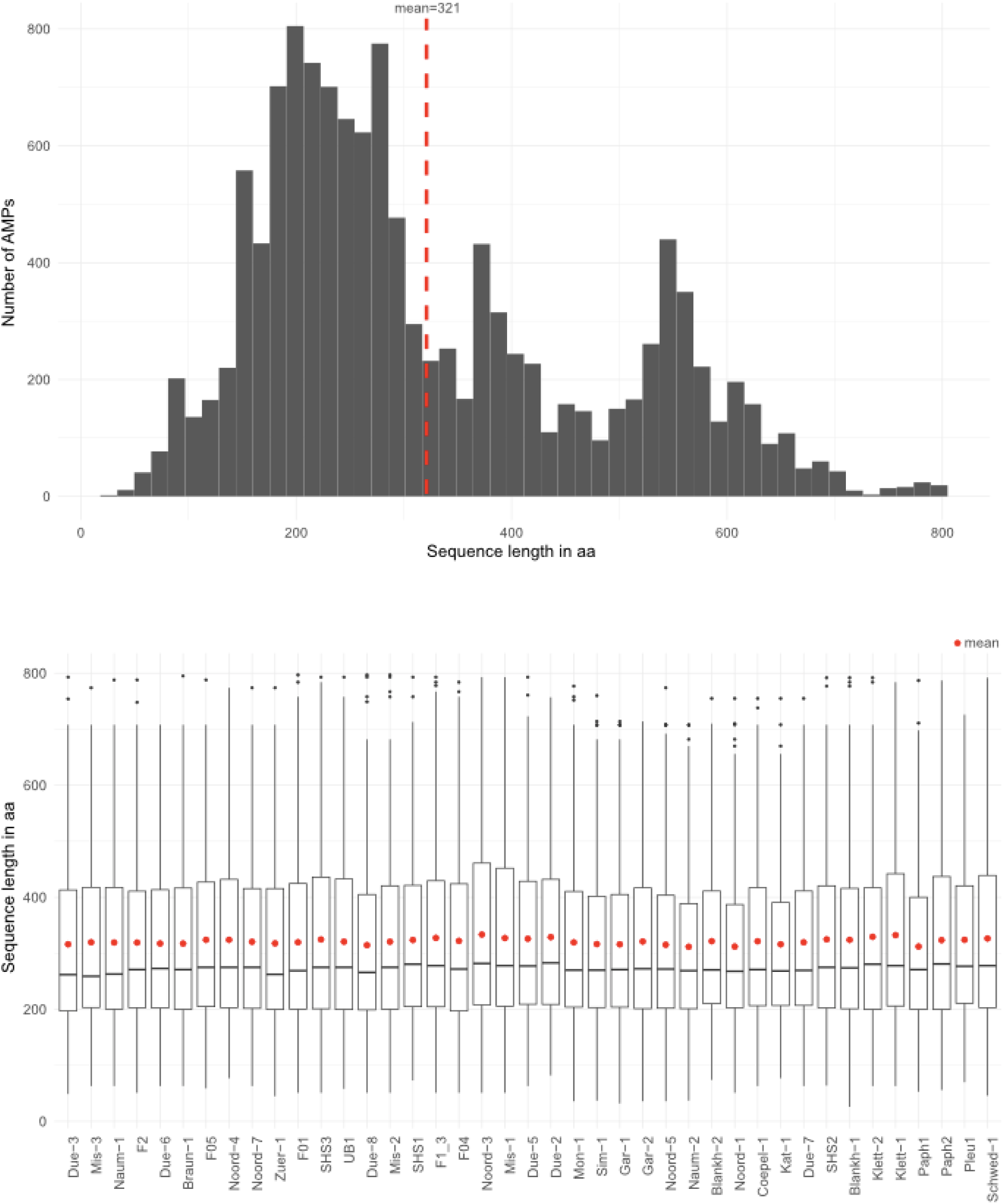
AMP length distribution across all mycobionts and per isolate. (Top) Distribution of AMP sequence lengths across all 41 mycobiont isolates. (Bottom) AMP length distributions shown for each individual isolate. Mean represented by red line or red dot.

**Supplementary Figure S9.**
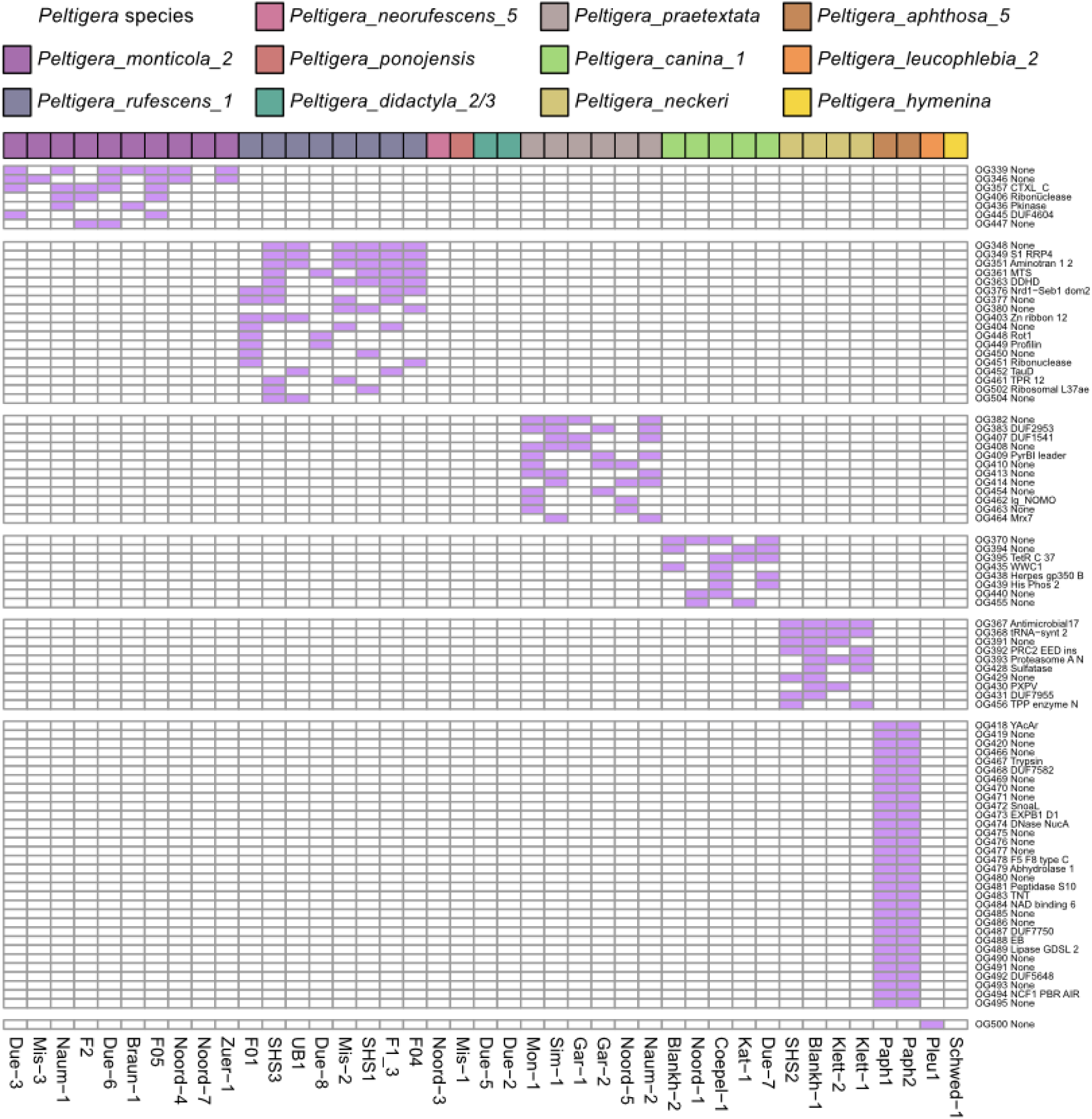
Species-specific AMP orthogroups of *Peltigera* mycobionts. Presence/absence heatmap of species-specific orthogroups. Purple indicates presence, white indicates absence. Orthogroups are functionally annotated and isolates are grouped by species which are represented by the different colours at top of the heatmap.

**Supplementary Figure S10.**
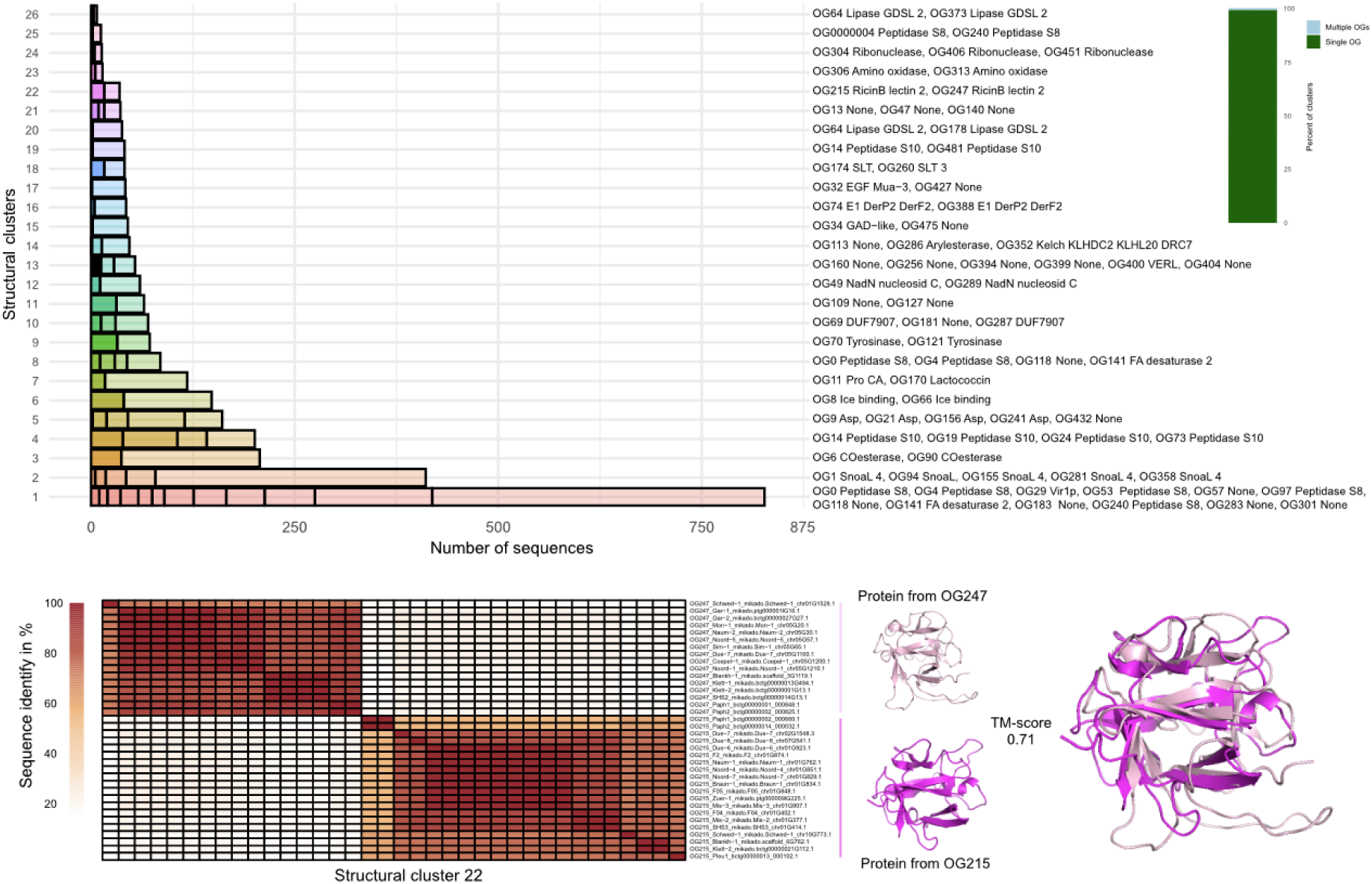
Comparison of sequence-based orthogroups and structure-based clustering of mycobiont AMPs. Top: Bar plot showing the proportion of structural clusters containing proteins from single (green) or multiple orthogroups (blue). Bar plot showing annotation of the 26 mixed structural clusters containing multiple orthogroups with annotation on the right. Bottom: Example of a mixed cluster (cluster 22) comprised of proteins from OG215 (magenta) and OG247(pale pink). Heatmap with sequence-based grouping with red representing high sequence similarity and white representing low sequence similarity. Structural alignment of one representative protein from each of the two orthogroups.

**Supplementary Figure 11:**
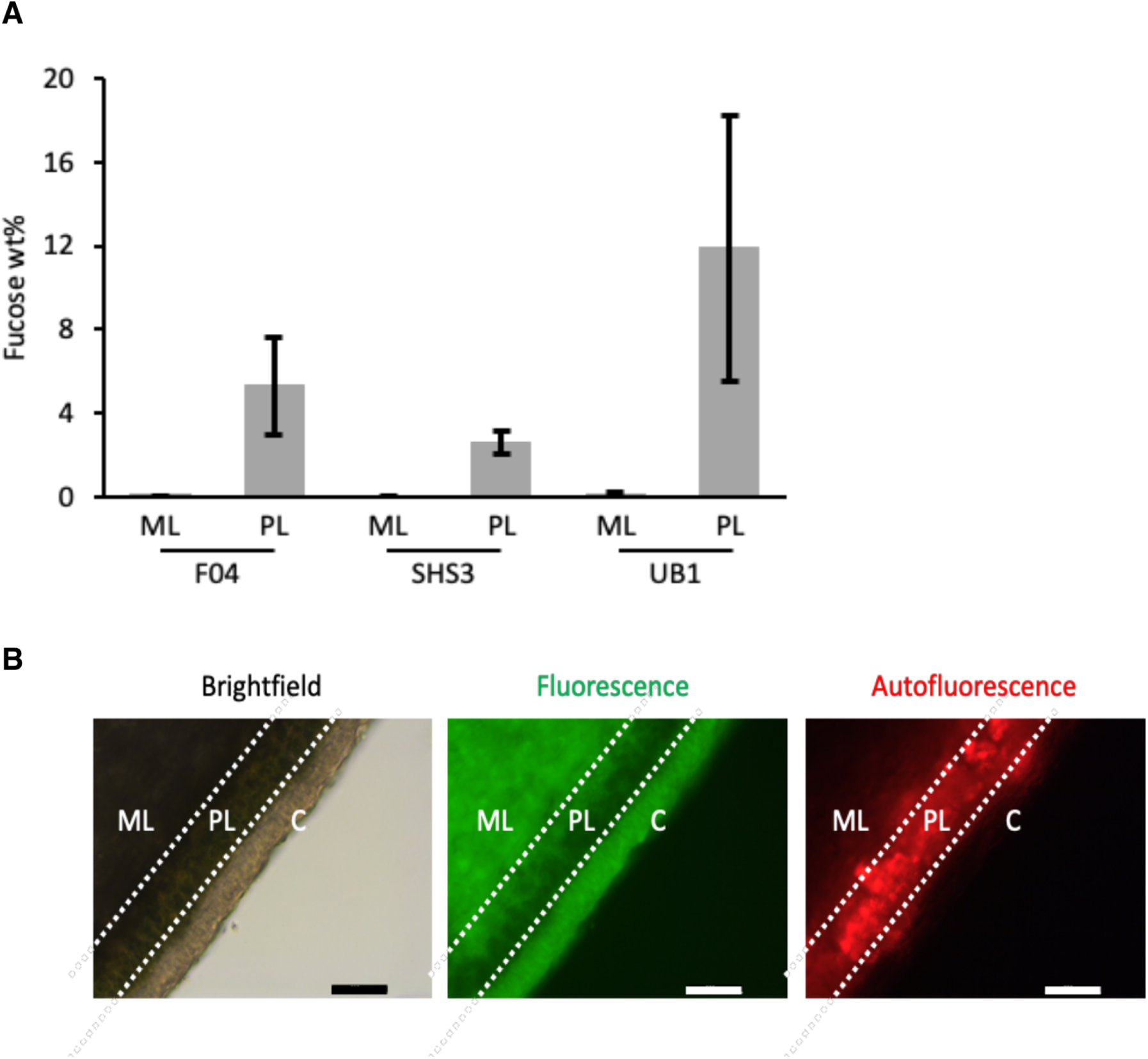
Cell Wall and Lectin analysis. A) Fucose (in weight % of total carbohydrates) in the medulla (ML) and photobiont (PL) layer of three *P. rufescens*_1 samples (F04, SHS3, UB1). Values are depicted as the average ± standard deviation of three replicates. B) Analysis with FITC labeled fucose coupled polymer to detect fucose binding lectins. ML denotes the medulla PL the photobiont layer and C the upper cortex layer. The scale bar represents 200 µm.

**Supplementary Figure 12.**
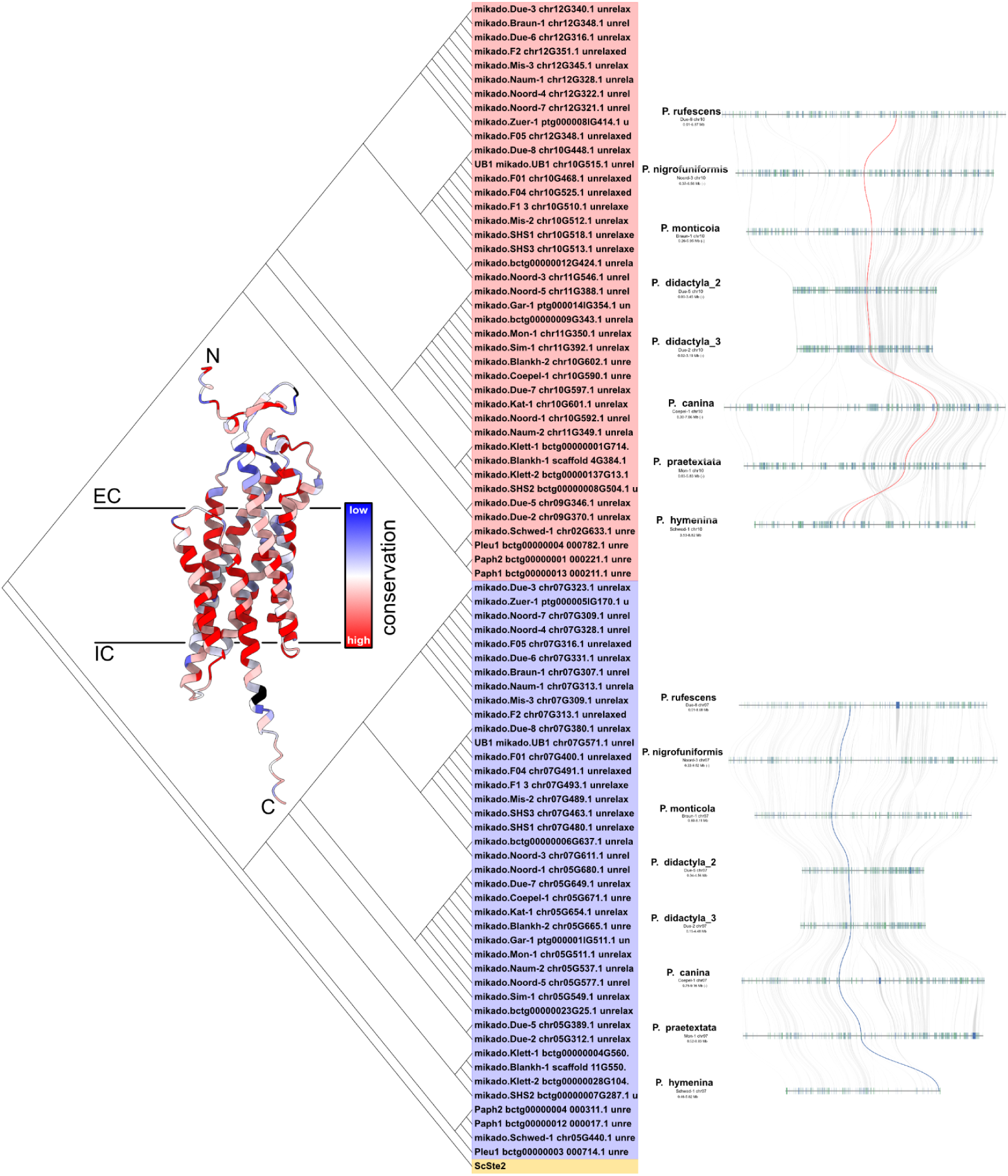
Conservation analysis of Ste2-like GPCRs reveals two distinct protein clades across all *Peltigera* species. The sequence-similarity-tree was calculated using Jalview Neighbour Joining algorithm on a MSA generated using the MUSCLE web tool with sequences of all Ste2-likes identified in GPCR analysis (two per sampled genome) plus *S. cereviseae* Ste2 as an outgroup representative. The tree was rooted to the outgroup. Grouping of Peltigera Ste2-likes into two distinct clades is based solely on sequence conservation. Position of both Ste2-like GPCRs on chromosome 7 and 10 respectively reveals conservation of specific Ste2-like type per chromosome, as well as conservation of genomic position across all depicted species. Mapping of all Ste2-like GPCRs onto representative structure from UB1 reveals sequence variability in extracellular regions of Ste2-likes.

**Supplementary Figure 13:**
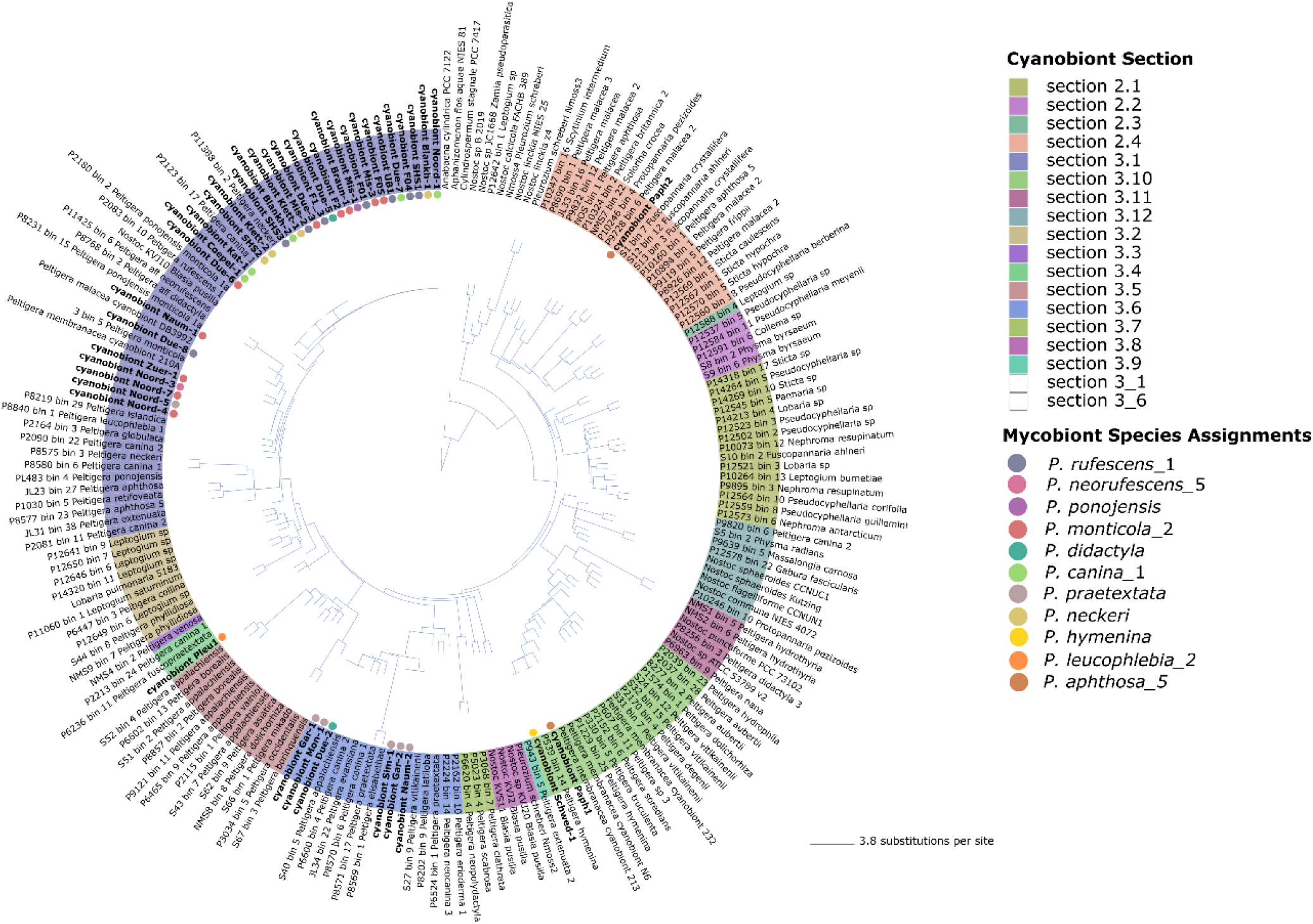
Phylogenomic tree of cyanobionts. The phylogenomic tree includes the 41 cyanobiont genomes newly assembled in this study, highlighted as bold text. Background colors indicate the assigned cyanobiont sections, while colored inner dots denote the associated mycobiont species assignments.

## Supplementary Tables

Supplementary Table S1: Overview of *Peltigera* samples, sampling sites, and associated metadata

Supplementary Table S2: Sequencing and Read Statistics

Supplementary Table S3: EukCC-estimated completeness and contamination of *Peltigera* mycobiont genomes

Supplementary Table S4: Genome assembly and annotation statistics of *Peltigera* mycobionts

Supplementary Table S5: Craq Genome Quality Metrics

Supplementary Table S6: Presence–Absence Matrix of InterPro IPR Domains and KEGG terms Across 41 Genomes

Supplementary Table S7: Presence–Absence Matrix of filtered InterPro IPR Domains and KEGG terms Across 41 Genomes

Supplementary Table S8: Cyanobacterial photobiont genome and proteome statistics

Supplementary Table S9: Phylogenetic assignments of cyanobacterial photobionts inferred from rbcLX sequences.

Supplementary Table S10: Algal photobiont genome statistics

Supplementary Table S11 Metadata for the 141 lichen-forming fungi *Peltigera* genomes used in the study

Supplementary Table S12: Transposable element and repeat composition of mycobiont genomes identified using PanEDTA

Supplementary Table S13: Presence–Absence Matrix of Orthologous Groups Identified by OrthoFinder

Supplementary Table S14: Biosynthetic gene clusters (BGCs) identified using antiSMASH

Supplementary Table S15: Peltigera AMP gene clusters

Supplementary Table S16: Peltigera genes with orthologs in the Pathogen–Host Interactions Database (PHI-base)

Supplementary Table S17: Genomic proximity context of captain-like YR genes in Peltigera mycobiont genomes

Supplementary Table S18: Presence/absence matrix of lectin-associated InterPro (IPR) numbers in Peltigera mycobiont functional annotations

Supplementary Table 19: Lectin candidates and Lectin associated candidates retained after multiple filtering steps

Supplementary Table S20: Cell wall composition

Supplementary Table S21: Cladonia G Protein assignment

Supplementary Table S22: Cladonia G Protein assignment comparison to Li et al.

Supplementary Table S23: G Proteins identification in the genomes including thresholds

Supplementary Table S24: G Proteins classes per genome

Supplementary Table S25: Gene expression data

Supplementary Table S26: Top Differentially Expressed genes

Supplementary Table S27: AMP Gene expression

Supplementary Table S28: Lectin Gene expression

Supplementary Table S29: GPCR Gene expression

Supplementary Table S30: Statistics of Oxford Nanopore cDNA reads

Supplementary Table S31: Basecaller information for the sequencing data generated in this study.

Supplementary Table S32: Assembly strategies and parameter settings of *Peltigera* mycobiont genomic assemblies

Supplementary Table S33. Information of the 25 genomes used for the bitscore threshold calculation for the loci used in the phylogenetic placement analysis

Supplementary Table S34. Calculated bitscore thresholds for the HMMER search

Supplementary Table S35: Assembly strategies and parameter settings of genomic assemblies from which Nostoc photobiont contigs were extracted

Supplementary Table S36. InterPro accessions used for the identification and filtering of lectin candidates.

